# Mapping Cellular Interactions from Spatially Resolved Transcriptomics Data

**DOI:** 10.1101/2023.09.18.558298

**Authors:** James Zhu, Yunguan Wang, Woo Yong Chang, Alicia Malewska, Fabiana Napolitano, Jeffrey C. Gahan, Nisha Unni, Min Zhao, Rongqing Yuan, Fangjiang Wu, Lauren Yue, Lei Guo, Zhuo Zhao, Danny Z. Chen, Raquibul Hannan, Siyuan Zhang, Guanghua Xiao, Ping Mu, Ariella B. Hanker, Douglas Strand, Carlos L. Arteaga, Neil Desai, Xinlei Wang, Yang Xie, Tao Wang

**Affiliations:** Quantitative Biomedical Research Center, Peter O’Donnell Jr. School of Public Health, University of Texas Southwestern Medical Center, Dallas, TX, 75390, USA; Division of Pediatric Gastroenterology, Hepatology and Nutrition, Cincinnati Children’s Hospital Medical Center, Cincinnati, OH, 45229, USA; Department of Pediatrics, University of Cincinnati, OH, 45221, USA; Department of Urology, University of Texas Southwestern Medical Center, Dallas, TX, 75390, USA; Harold C. Simmons Comprehensive Cancer Center, University of Texas Southwestern Medical Center, Dallas, TX, 75390, USA; Department of Internal Medicine, University of Texas Southwestern Medical Center, Dallas, TX, 75390, USA; Department of Pathology, University of Texas Southwestern Medical Center, Dallas, TX, 75390, USA; Department of Computer Science and Engineering, University of Notre Dame, Notre Dame, IN, 46556, USA; Department of Radiation Oncology, University of Texas Southwestern Medical Center, Dallas, TX, 75390, USA; Department of Bioinformatics, University of Texas Southwestern Medical Center, Dallas, TX, 75390, USA; Department of Molecular Biology, UT Southwestern Medical Center, Dallas, TX, 75390, USA; Hamon Center for Regenerative Science and Medicine, UT Southwestern Medical Center, Dallas, TX, 75390, USA; Department of Mathematics, University of Texas at Arlington, Arlington, TX, 76019, USA; Center for Data Science Research and Education, College of Science, University of Texas at Arlington, Arlington, TX, 76019, USA

**Keywords:** spatially resolved transcriptomics, cellular interaction, spacia, MERSCOPE, GeoMX

## Abstract

Cell-cell communication (CCC) is essential to how life forms and functions. However, accurate, high-throughput mapping of how expression of all genes in one cell affects expression of all genes in another cell is made possible only recently, through the introduction of spatially resolved transcriptomics technologies (SRTs), especially those that achieve single cell resolution. However, significant challenges remain to analyze such highly complex data properly. Here, we introduce a Bayesian multi-instance learning framework, spacia, to detect CCCs from data generated by SRTs, by uniquely exploiting their spatial modality. We highlight spacia’s power to overcome fundamental limitations of popular analytical tools for inference of CCCs, including losing single-cell resolution, limited to ligand-receptor relationships and prior interaction databases, high false positive rates, and most importantly the lack of consideration of the multiple-sender-to-one-receiver paradigm. We evaluated the fitness of spacia for all three commercialized single cell resolution ST technologies: MERSCOPE/Vizgen, CosMx/Nanostring, and Xenium/10X. Spacia unveiled how endothelial cells, fibroblasts and B cells in the tumor microenvironment contribute to Epithelial-Mesenchymal Transition and lineage plasticity in prostate cancer cells. We deployed spacia in a set of pan-cancer datasets and showed that B cells also participate in *PDL1*/*PD1* signaling in tumors. We demonstrated that a CD8^+^ T cell/*PDL1* effectiveness signature derived from spacia analyses is associated with patient survival and response to immune checkpoint inhibitor treatments in 3,354 patients. We revealed differential spatial interaction patterns between γδ T cells and liver hepatocytes in healthy and cancerous contexts. Overall, spacia represents a notable step in advancing quantitative theories of cellular communications.

## INTRODUCTION

Various types of cells form complex structures and communication networks in the tissue micro-environment, and the signaling between these cells is central to normal organ development and diseased physiological processes. Elucidating cell-cell communication (CCC) in the tissue microenvironment in different biological systems is of vital importance. Many experimental and informatics approaches have attempted to address this question ^1–4^.

Since the introduction of single-cell RNA-sequencing (scRNA-seq) data, many informatics approaches to infer CCCs based on scRNA-seq data have been developed, such as CellChat ^5^, NicheNet ^6^, CellphoneDB ^7^, NATMI ^8^, SingleCellSignalR ^9^, *etc* (some reviewed in **Sup. Table 1**). Despite their popularity and their significant contributions to the field, these methods suffer from a number of major caveats, due to the limited information provided by scRNA-seq. To begin with, many of these tools only infer interactions between cell types rather than interactions at the single-cell level, thus losing single cell resolution. Secondly, CCC is usually context-specific, and the common approach of mapping the data to pre-defined interaction pathway databases, regardless of the cellular context, inevitably results in low resolution and low specificity in the detection of true CCCs in the tissue sample of interest. Lastly, most of these tools rely on the co-expression of ligand-receptor gene pairs in signal-sending and receiving cells to claim detection of CCC. However, the expression of the receptor gene itself is not necessarily impacted by the expression of the ligand. But rather, the ligand’s impact can only be revealed by the transcriptomic regulation of (some of) the receptor’s downstream genes.

Fortunately, emerging spatially resolved transcriptomics (SRT) technologies potentially provide promising data to address these problems, by providing the extra spatial modality to aid the dissection of CCCs. Some bioinformatics tools have been developed for inferring CCCs from such data (reviewed in **Sup. Table 1**). However, earlier SRT methods, such as Visium/10X, Slide-Seq ^10,11^, and XYZeq ^12^, are all low resolution SRT methods, meaning that they cannot achieve single cell resolution in the SRT expression data and their usefulness for CCC inference is still very limited. For example, one Visium sequencing spot can contain 10 or even more cells. The cells that are interacting with each other could be contained in one sequencing unit, thus making it difficult to infer such types of CCCs. Therefore, bioinformatics tools developed based on such low resolution SRT data still have not addressed the problem of accurately inferring CCCs. And many of the same caveats that we mentioned above for scRNA-seq-based methods are still outstanding.

Only very recently are single cell resolution SRT technologies developed and commercialized, such as Seq-scope ^13^, HDST ^14^, MERSCOPE/Vizgen, CosMx/Nanostring and Xenium/10X, *etc*. First, these newest SRTs allow researchers to pinpoint each single cell with its spatial location and identify the other cells within its neighborhood. It is thus feasible to detect interactions between each pair of single cells without having to aggregate to cell types. In addition, since SRTs enable examining the expression profiles of all possible pairs of single cells, yielding very rich data that reveal ongoing CCCs, we can apply a much more data-driven approach to infer CCCs, overcoming the constraint of relying on previously known pathways. Furthermore, the high-dimensional and multi-modal information contained in these cell pair data also enables the modeling of downstream target genes in the receiving cells in detail, removing the limitation to ligand-receptor pairs documented by prior databases. Therefore, single cell SRTs overcome all major caveats encountered in mapping of CCCs from scRNA-seq data and low resolution SRT data.

However, significant data analytical challenges came with the maturation of these state-of-the-art SRT technologies. In particular, during CCCs, multiple senders in a neighborhood could confer signals to and impact the same receiver. SRTs allow us to match these potential senders and receivers. But these pairing relationships result in complex multi-to-one data structures consisting of both expression and spatial modalities that are unsuitable for classical statistical learning approaches, and have rarely been considered sufficiently by existing CCC detection approaches so far (**Sup. Table 1**). To address this inadequacy, we innovate spacia, a Bayesian multi-instance learning (MIL) framework, to explicitly model these multi-to-one relationships and to detect the interaction between individual signal-sending and receiving cells from SRT data.

Spacia achieved single-cell resolution detection of CCCs, models the downstream targets of CCCs, rather than being limited to ligand-receptor relationships documented in prior databases, and significantly reduced artefacts of CCC inferences while increasing specificity. We tested spacia on data from all three commercialized single cell resolution SRTs so far: MERSCOPE, CosMx, and Xenium, proving the suitability of spacia and these SRT platforms for inferring CCCs. **Sup. Table 1** summarized these advances of spacia over prior works in terms of conceptual advantages and comprehensiveness of testing datasets. In real data applications, spacia out-performed benchmark tools by a large margin and revealed novel biological insights regarding CCCs from diverse biological contexts.

## RESULTS

### Multi-Instance Learning for Mapping CCCs from SRT Data

We present a Bayesian multiple-instance learning (MIL) model, spacia, to infer CCCs from SRT data. Under a MIL framework, the receiver cells are modeled as “bags” with labels (expression of receiving genes/pathways), and each bag is a collection of multiple instances (senders) characterized by the instance-level features (expression of genes/pathways and spatial closeness to receivers). In this way, the multiple-sender-to-one-receiver relationships during CCCs are explicitly modeled. The technical details of spacia are described in **Sup. File 1** and the statistical innovation of spacia is provided in the discussion section. At a high level, spacia has two tiers. The first tier identifies signal-sending cells (senders) that are truly impacting signal-receiving cells (receivers) based on spatial closeness, indicated by a *δ* variable (**Fig. 1ab**), out of all candidate sending cells included into each bag. As different CCCs could have varying effective spatial ranges, depending on the cell types and types of interactions (contact-based or secretion-based), the model estimates a variable, indicated by *b*, to allow flexibility in determining *δ* in each specific dataset. When *b* is negative, the senders and receivers are determined to have stronger interactions when they are physically closer. In the second tier, spacia discovers gene co-expression patterns between the senders and the receivers (**Fig. 1ab**), but only for senders that are determined to be impacting each receiver cell (*δ=1*). We infer a variable *β* that indicates how the expression of the genes/pathways in the senders impacts the genes/pathways in the receiver cells. This model is solved by Markov Chain Monte Carlo (MCMC), which is an iterative process that generates a distribution for the value of each variable of interest so that we can provide both point estimates and inferences of statistical significance.

**Fig. 1.**
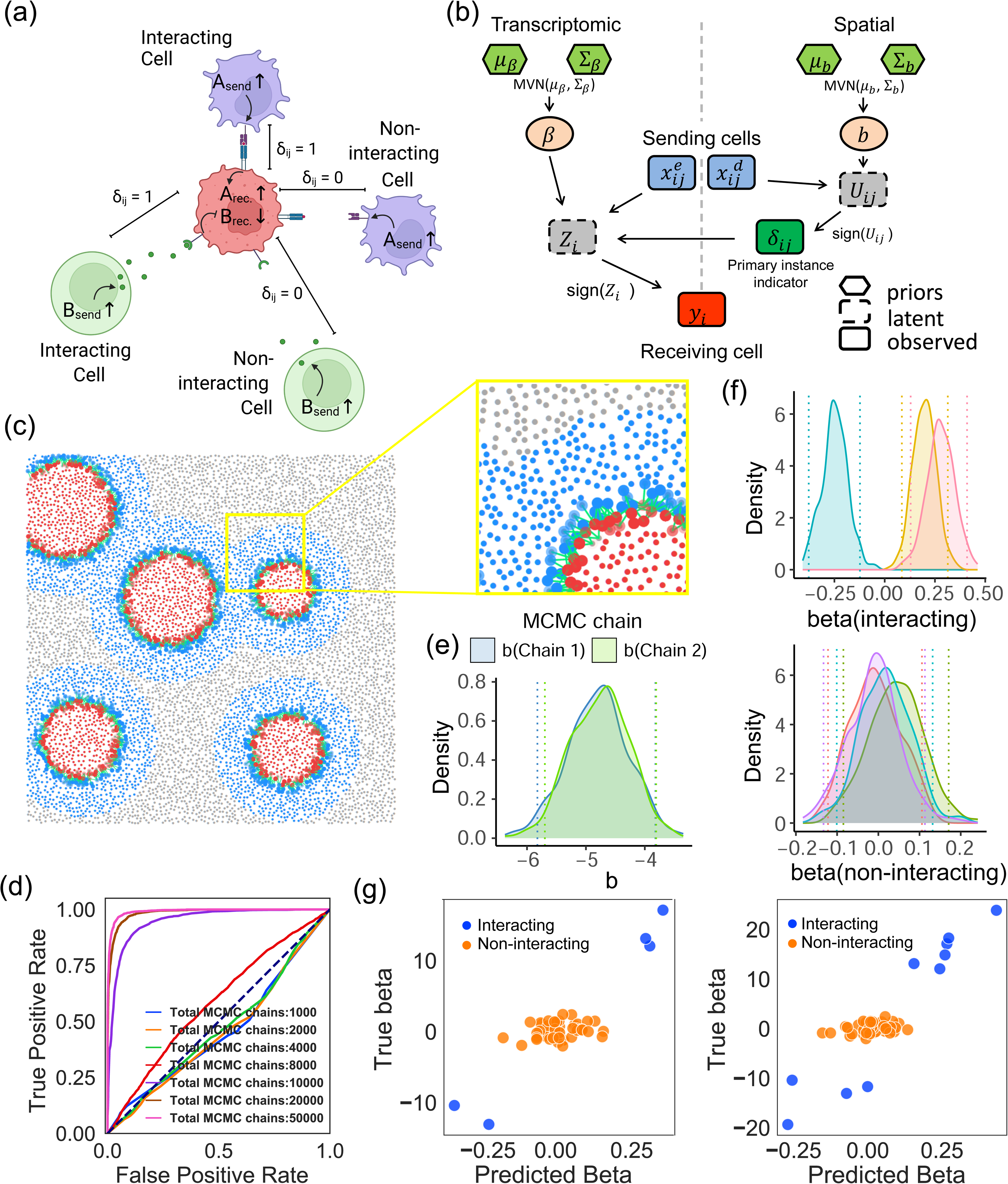
The spacia model. (a) Cartoons explaining the concept of “primary instances”, namely the sending cells that are truly interacting with the receiver cells. The purple senders refer to senders that interact with receivers through cell-to-cell contact. The green senders refer to senders that interact with receivers through secreted ligands. (b) Diagram showing the key elements of the model structure of spacia. (c) Inference results of spacia on a simulation dataset. Blue color refers to sender cells, red color refers to receiver cells and green arrows refer to CCC. (d) ROC curves measuring the accuracy of spacia finding the correct primary instances in the simulation data, with increasing numbers of total MCMC chains. (e) The distribution of the *b* variables across MCMC iterations after stabilization, in two MCMC chains. (f) The distribution of the *β* variables across MCMC iterations after stabilization, for genes in senders that are (top) or are not (bottom) truly interacting with receiving genes. (g) Point estimates of the *β* variables. Left: 5 truly interacting genes in the simulation data; right: 10 truly interacting genes.

Spacia infers, for each receiver cell, a set of sender cell(s) that truly interact with this receiver. This unique model design enables us to accurately infer single cell-to-single cell interactions for each cell captured by SRT. On the other hand, the sending and receiving genes that can be considered by our model do not have to be ligands and receptors, but rather, all genes captured by SRT can be considered for both the sending and receiving portions. This allows us to avoid the questionable rationale of examining the co-expression of ligands/receptors from pre-defined interaction databases for mapping CCC, while allowing us to model the upstream and downstream regulatory signals that occur during CCC.

Importantly, cell-cell communications happen in a variety of manners. As reviewed by Armingol *et al* ^2^, there are four major types of cell-cell communications: autocrine, paracrine, juxtacrine, and endocrine. The first three types of communication naturally require cells to be in close proximity. In contrast, endocrine interactions occur over long distances through systemic circulation, and it is not feasible to track such CCCs by SRT. Similarly, some CCCs, though happening within one organ, also still do not depend on spatial proximity of the interacting cells. It is important to note that spacia only considers the types of cell-cell communications that require the interacting cells to be closely localized, in addition to demonstrating co-expression patterns.

To validate its efficacy, we tested spacia on simulated data. As we demonstrated in **Fig. 1c**, we simulated two types of cells that are interacting. The blue cells are senders, while the red cells are receivers, but the red cells’ expression was simulated to be regulated by only nearby blue cells. As expected, spacia only infers red and blue cells that are close to each other to be interacting (**Fig. 1c**). We showed, in **Fig. 1d**, the rate of correct identification of the truly interacting cell pairs, measured by Area Under the ROC curve (AUROC). With more than 10,000 MCMC iterations, the AUROC achieves >0.95 (spacia’s default is 50,000 iterations), meaning an almost perfect detection of interacting cell pairs. We then showed in **Fig. 1e** that the *b* variables were estimated to be negative, consistent with the simulation assumption that only nearby cells are interacting. We also evaluated spacia’s estimations of the *β* variables, in **Fig. 1f** (distributions of *β*s across all MCMC iterations) and **Fig. 1g** (point estimates of *β*s). As is shown, the sender genes that were simulated to be truly interacting with the receiver genes had *β*s that were significantly different from 0, while the converse was true for non-interacting genes. Spacia allows us to perform statistical inference of the inferred *b* and *β* variables. For example, their confidence intervals are visualized in **Fig. 1ef**, according to the sampling from the MCMC iterations, from which we can also calculate P values. The P values are <1E-4 for both chains for *b* (**Fig. 1e**), range from 0.008 to 3.4E-5 for the truly interacting genes’ *β*s (top panel of **Fig. 1f**), and range from 0.09-0.77 for the non-interacting genes’ *β*s (bottom panel of **Fig. 1f**). Finally, the posterior samples of the *b* and *β* variables demonstrate stable convergence and minimum auto-correlations (**Sup. Fig. 1**). Additional analyses were provided in **Sup. File 2**. Taken as a whole, these results indicate that spacia exhibits excellent statistical properties.

### Spacia Validates in SRT Data and Overcomes Limitations of Existing Approaches

Next, we investigated a prostate cancer MERSCOPE dataset and used spacia to infer how the non-tumor cells from the tumor microenvironment impact prostate cancer cells (**Fig. 2a**). We compared spacia to several SRT-based CCC inference tools, including SpaTalk ^15^, COMMOT ^16^, and SpatialDM ^17^, that achieved high to top advantage points in **Sup. Table 1**, and also two popular non-SRT-based CCC inference software, CellPhoneDB ^7^ and CellChat ^5^. We visualized the interacting cell pairs in their spatial locations in **Fig. 2b** and **Sup. Fig. 2ab**. Consistent with our expectation, the inferred CCCs are all local for spacia, COMMOT, and SpaTalk. In contrast, the CCCs inferred from CellPhoneDB and CellChat indicate numerous CCCs across the entire span of the tissue, which is highly unlikely. In addition, some software such as SpatialDM and COMMOT cannot determine individual interaction strengths between single cells (or sequencing units). SpatialDM only outputs a score measuring the probability that each single cell participates in CCCs (as denoted in **Sup. Fig. 2a**), while not inferring the interaction partners. COMMOT only outputs a direction of interaction for each cell, without the direction pointing to any specific interacting cells.

**Fig. 2.**
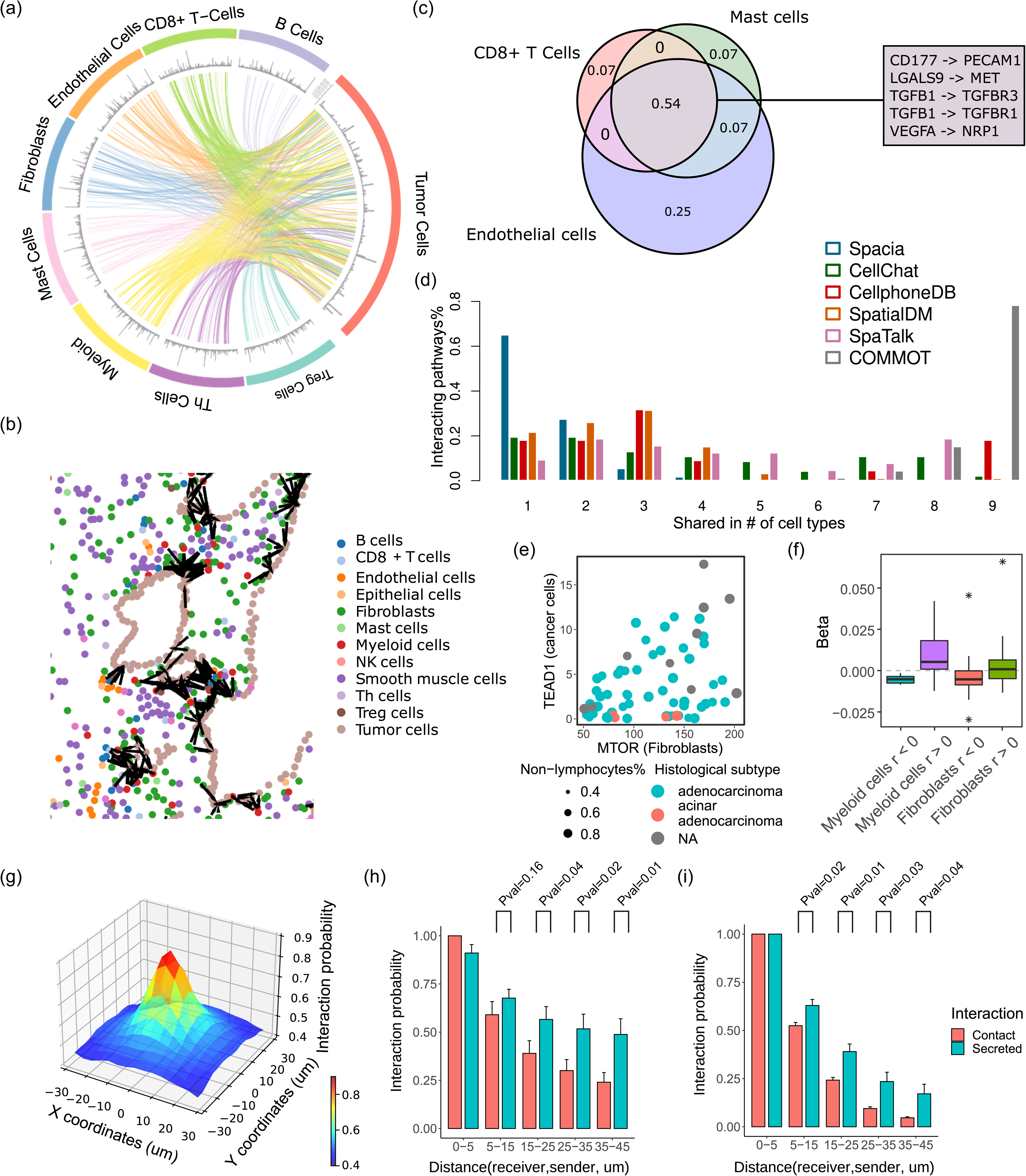
Validating spacia in real data. (a) Spacia inferred CCCs from select immune and stromal cells to tumor cells in the prostate cancer MERSCOPE dataset. The spacia results were filtered to only visualize those that satisfy *b* < 0, *b* Pval<1x10E-30, *β* Pval<0.001, and top 1% of z-scored *β* across all results. (b) Spatial representation of the spacia results. Cells are color labeled by cell types, and significantly interacting cell pairs inferred by spacia (*b*s) are indicated in black. (c) Overlap between the predicted CCCs by CellphoneDB on the same MERSCOPE dataset for three example sending cell types. The numbers indicate the proportions of total CCCs found in each overlap. Some example CCCs that are inferred to exist in all three cell types are listed. (d) Comparison of the degree of overlap in inferred CCCs between different sending cell types, by spacia, CellChat, CellphoneDB, SpatialDM, SpaTalk, and COMMOT. The Y axis refers to the proportions of interactions shared in *n* cell types *vs.* all predicted interactions. (e) An example scatterplot showing the correlations between the expression of sending and receiving genes in their respective cell types in the PDX RNA-seq data. (f) Spacia *β* values are more likely to be positive for sending genes and receiving genes that are positively correlated in the PDX data, and *vice versa*. (g-i) Probability of interaction as a function of distance from the sending cells to the receiving cell. (g) Gene pairs of both secretion-based and contact-based CCCs combined in the prostate cancer MERSCOPE dataset. The sending cells are shown in their actual X-Y coordinates relative to the receiving cells and the interaction probabilities are averaged at each location. The Z axis shows the primary instance probabilities of the sending cells, explained in **Sup. File 1**. (h,i) The probabilities for each type of CCCs shown separately, and the sending cells are ordered and binned along the distances between sending and receiving cells. (h) the prostate cancer and (i) the lung cancer MERSCOPE datasets. For sake of consistency, for each gene pair shown in panels (g-i), all interaction probabilities are normalized by the maximum interaction probabilities and shown in the plots. T test is used to test if the sending-receiving cell pairs have higher interaction probabilities in secretion-based CCCs than in contact-based CCCs.

More importantly, we noticed the benchmark tools inferred highly similar sets of interacting genes regardless of the types of cells. For example, in **Fig. 2c**, we showed CellPhoneDB’s inference results for three very dissimilar cell types (as sending cells), CD8^+^ T cells, mast cells, and endothelial cells. Among all unique sending-receiving gene pairs that were inferred to be active in at least one of these sending cell types, 54% exists in all three sending cell types, which suggests an alarming lack of specificity. We investigated this systematically by examining how many predicted interacting genes were shared between all sender cell types (**Fig. 2d**). Of all the CCCs inferred by spacia, 92.7% were unique to one or two sender cell types, with no interactions found to be shared by more than four different sender cell types. In striking contrast, 63.6% of the CCCs are shared among three or more cell types according to CellphoneDB, 60.9% according to CellChat, 52.2% for SpatialDM, 71.9% for SpaTalk, and 100% for COMMOT. In particular for COMMOT, 78.3% of interactions were shared among all cell types examined, which is impossible.

Furthermore, we note that, for all of COMMOT, SpatialDM and SpaTalk, if we ran these tools on the full SRT dataset, the memory usage exceeds 1-2TB, which is far more than the maximum memory limit of regular high performance computing servers and the programs all failed. We had to downsample the number of cells in the SRT data dramatically (maximum of 5,000 for COMMOT, 1,000 for SpatialDM, and 3,000 for SpaTalk, for each cell type) so that the execution would be successful.

We utilized other forms of data to validate spacia. The NCI Patient-Derived Models Repository (PDMR, pdmr.cancer.gov) provides RNA sequencing data of 70 prostate cancer patient-derived xenografts (PDXs). In PDX models, human immune/stromal cells die out quickly and mouse T/B/NK cells are generally non-existent due to the choice of NSG mouse models ^18^. We separated the bulk PDX gene expression into the human tumor cell *vs.* mouse stromal/immune cell components with Disambiguate ^19^. Next, we used CIBERSORTx ^20^ to further demultiplex the mouse stroma/immune cell component into cell type-specific expression. We then evaluated the correlation between human tumor cells’ expression and the expression of mouse fibroblasts and myeloid cells (one example gene pair shown in **Fig. 2e**), which are the two most abundant cell types in the murine components, according to CIBERSORTx. Then we cross-referenced these against the pairs of signal-sending genes in senders (fibroblasts and myeloid cells) and downstream target genes in receiver tumor cells that were detected by spacia. Overall, positively correlated gene pairs from the PDX data are indeed more likely to have positive βs in the corresponding interactions inferred by spacia and *vice versa* (**Fig. 2f**).

Finally, we categorized the interacting gene pairs into “contact-based” and “secretion-based” interactions according to whether the sending gene is known to participate in contact-based or secretion-based interactions ^5^, and investigated cellular interaction probabilities as a function of the distances between sending and receiving cells (**Fig. 2g**). As is shown in **Fig. 2h**, the probabilities of interactions drop off along with increasing distances in both types of interactions, but the interaction probabilities drop off much faster for contact-based interactions than for secretion-based interactions, which is consistent with the fact that secretion-based interactions can happen over longer distances by moving of secreted molecules from one cell to another. We also investigated another MERSCOPE dataset from a lung cancer patient and observed the same phenomenon (**Fig. 2i**).

### Induction of Prostate Cancer EMT and Lineage Plasticity by the Tumor Microenvironment

TGFBs secreted by fibroblasts induce EMT in tumor cells ^21,22^. We investigated whether the sender genes are enriched in signaling pathways that are associated with the induction and regulation of EMT in tumor cells. We performed Gene Ontology analysis using GOrilla ^23,24^ on the sending genes, for each sender cell type and for each of several receiving genes in the tumor cells that are classical markers or drivers of EMT ^25–33^ (**Fig. 3a**). We found that fibroblasts, B cells, endothelial cells and T_h_ cells possess the highest numbers of enriched pathways (FDR<0.05) directly or closely related to the regulation of EMT (the full GO results for *JAK1* as an example receiver gene are shown in **Sup. Table 2**). In other words, spacia correctly inferred the upstream signaling pathways in several cell types of the tumor microenvironment that regulate the downstream induction of EMT in the tumor cells.

**Fig. 3.**
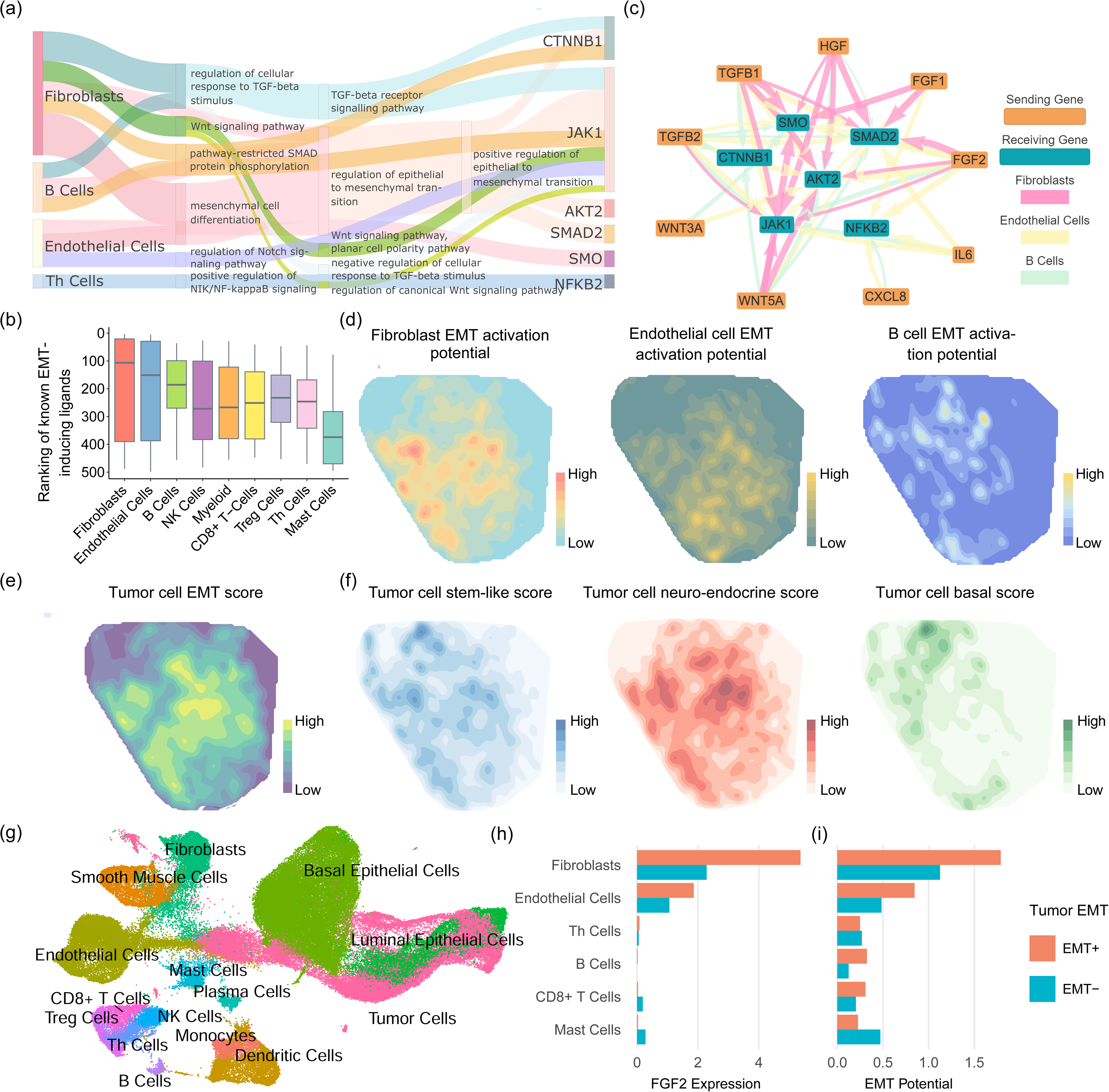
Applying spacia to reveal EMT and lineage plasticity induction signals from the prostate cancer tumor microenvironment. (a) Sankey diagram describing GO term enrichment in the sending genes, of each sender cell type, that are inferred by spacia to be impacting well known EMT marker genes in the tumor receiver cells. The width of the flow is scaled with the P values, and the parent term is connected with the child term if both terms were identified between the same sender cell-receiving gene pair. (b) The rankings of *β*s for known cytokine ligands that could induce EMT, among all the sending genes input into spacia, for each cell type. A smaller rank refers to a stronger interaction strength (larger *β*). (c) The top sending-receiving gene pairs inferred by spacia, for fibroblasts, endothelial cells and B cells as sending cell types. (d) The kernel density estimation of the EMT activation potentials in fibroblasts, endothelial cells and B cells. (e) The kernel density estimation of the EMT levels in prostate cancer cells. (f) The kernel density estimations of the stem-like, neuro-endocrine, and basal lineage plasticity scores in the prostate cancer cells. (g) UMAP plot showing the cell types of single cells from our prostate cancer patients. (h) The *FGF* expression of different sending cell types in the EMT+ and EMT-patients. (i) The EMT activation potentials of different sending cell types in the EMT+ and EMT-patients.

We focused on a number of secretory ligands that are known to induce EMT activation in tumor cells within the sending cell types above; these ligands include *WNT5A*, *WNT3A*, *TGFB1*, *TGFB2*, *FGF2*, *IL6*, *CXCL8*, *HGF*, and *FGF1* ^21,22,25,26,28–30,32^. **Fig. 3b** shows the ranked *β*s of these ligands in each sender cell type, among all 500 genes captured by MERSCOPE, where a smaller rank refers to stronger up-regulation of the EMT marker/driver genes. Fibroblasts (Pval=0.03, one-sided T-test against the mean rank of 250), endothelial cells (Pval=0.04), and B cells (Pval=0.005) demonstrate the strongest activation of tumor EMT *via* these ligands. While the roles of fibroblasts and endothelial cells in inducing EMT are more well established, it is surprising to see that B cells are also inferred by spacia to induce EMT in prostate cancers. We also showed more details of the inferred interactions between these ligands and the EMT genes in a network plot (**Fig. 3c**). It is apparent that fibroblasts, endothelial cells and B cells each employ more than one secretory factor to induce tumor cell EMT. We also uncovered EMT-inducing interactions that have not been described before in prostate cancer. For example, while it has been reported that endothelial cells secrete IL-6 and induce EMT in head and neck tumors ^34^ and esophageal carcinoma ^35^, we showed that such a mechanism also exists in prostate cancers (**Fig. 3c**). We next visualized the spatial co-expression patterns of these ligands in sender cells (**Fig. 3d**) and the EMT levels of the tumor cells (**Fig. 3e**, definition of EMT level in **method section**). In particular, we aggregated over these secretary ligands to form an “EMT activation potential” in each sender, weighted by the βs inferred by spacia, and further aggregated all senders of each receiver cell through a weighted average with weights being the probability of the senders being “primary” (*δ=1*). **Fig. 3d** confirms that the EMT activation patterns have higher correlation with the EMT levels of the tumor for fibroblasts and endothelial cells (Spearman correlation: ρ=0.822, 0.838), but much lower for B cells (ρ=0.415) (**Fig. 3e**, **Sup. Table 3**). It is interesting, however, that fibroblasts and endothelial cells have largely overlapping spatial patterns of EMT activation, while B cells show a distinct pattern with its EMT activation. It appears that the sum of the EMT activation patterns of fibroblasts/endothelial cells and B cells better corresponds to the EMT level of the tumor cells (**Fig. 3e**).

Our prior work showed that EMT is usually part of a lineage plasticity dys-regulation program in prostate cancer cells ^36^. To determine if the observed increase in EMT activity was also associated with gene expression programs involved in lineage plasticity, we studied the spatial distribution of the stem-like, neuro-endocrine, and basal lineage plasticity levels in the tumor cells. **Fig. 3f** shows that the stem-like and neuro-endocrine lineages are largely correlated with the prostate cancer cell EMT levels (ρ=0.666, 0.854) and the EMT activation potential of fibroblasts and endothelial cells (**Sup. Table 3**). In particular, the correlation between fibroblast EMT activation potential and the tumor cell neuro-endocrine lineage is the highest, achieving a Spearman correlation of 0.916.

To validate the interactions inferred by spacia at the patient level, we generated 21 scRNA-seq datasets from a cohort of prostate cancer patients (**Fig. 3g**). We divided the patients into two subsets of EMT high (N=10) and EMT low (N=11), according to the EMT expression in their tumor cells. We first examined the expression of *FGF2* in fibroblasts, endothelial cells, B cells, and several other immune cell types as controls (**Fig. 3h**). **Fig. 3c** indicates that fibroblasts and endothelial cells are the major cell types that secrete FGF2 and induce prostate cancer cell EMT. Consistent with this observation from MERSCOPE, **Fig. 3h** shows that fibroblasts and endothelial cells are the only two cell types that abundantly express *FGF2*, and more importantly, the expression of *FGF2* is higher in fibroblasts (Pval<0.001) and endothelial cells (Pval<0.001) from patients with EMT-high tumors, compared with EMT-low tumors. We again examined all the secretory factors together by calculating the EMT activation potential as in **Fig. 3d**. As shown in **Fig. 3i**, fibroblasts cells overall possess the strongest EMT activation potential. As expected, fibroblasts and endothelial cells show stronger EMT activation potential in the EMT-high patients, compared with the EMT-low patients (Pval<0.001 for both). B cells also demonstrate the same trend, though not achieving statistical significance. Next, we assessed the association between lineage plasticities of tumor cells and the EMT activation potentials of the stromal/immune cells (**Sup. Fig. 3**), as is done for the MERSCOPE dataset. Consistent with **Sup. Table 3**, the most pronounced observation is that fibroblast EMT activation potential was significantly higher in the group associated with enhanced neuro-endocrine lineage (Pval<0.001). The endothelial EMT activation potential was also higher in the stem-like positive group than in the negative group (Pval<0.001), similarly for the basal lineage (Pval<0.001). Overall, the main discoveries regarding the intra-tumor heterogeneity of EMT and lineage plasticity induction in tumor cells identified by spacia can be extrapolated to inter-tumor differences.

Finally, as a comparison, we also performed the analyses of **Fig. 3b** for the five benchmark software applications. However, we note that none of these software applications outputs a direction of regulation. This is in sharp contrast to spacia, which outputs a *β* for each sending-receiving gene pair, whose direction clearly indicates an up-regulation or down-regulation effect. This is a significant advance of spacia over these five benchmark software applications and potentially many other existing CCC inference tools listed in **Sup Table 1**. Nevertheless, we ignored the direction of regulation for this particular analysis, and still found that none of these benchmark software applications can identify fibroblasts, endothelial cells, and B cells as the main drivers of tumor cell EMT (**Sup. File 2**).

### Spacia Infers the Impact of PDL1 Signaling on B cells in the Tumor Microenvironment

In another application, we deployed spacia to characterize the impact of *PDL1* signaling on the tumor microenvironment. While it is well known that tumor cells expressing *PDL1* inhibit T cells expressing *PD1*, whether other *PDL1*- and *PD1*-expressing cells interact with each other is not clear yet. We studied a breast cancer MERSCOPE dataset and applied spacia to investigate how the *PDL1* expression of tumor cells, endothelial cells, and macrophages leads to downstream transcriptional regulation in CD8^+^ T cells, T_h_ cells, T_reg_ cells, B cells, and macrophages (**Fig. 4a**). We visualized the pathways enriched in the downstream target genes that spacia inferred for different cell type pairs in **Sup. Fig. 4**. We show that the PDL1-PD1 axis is active in almost all possible combinations of sender cell type-receiver cell type pairs, although the strength of interactions is still the strongest for PDL1 signaling from tumor cells. It is also worth noting that, in addition to the enrichment of genes in pathways typically related to immune responses, there is also an enrichment related to apoptosis.

**Fig. 4.**
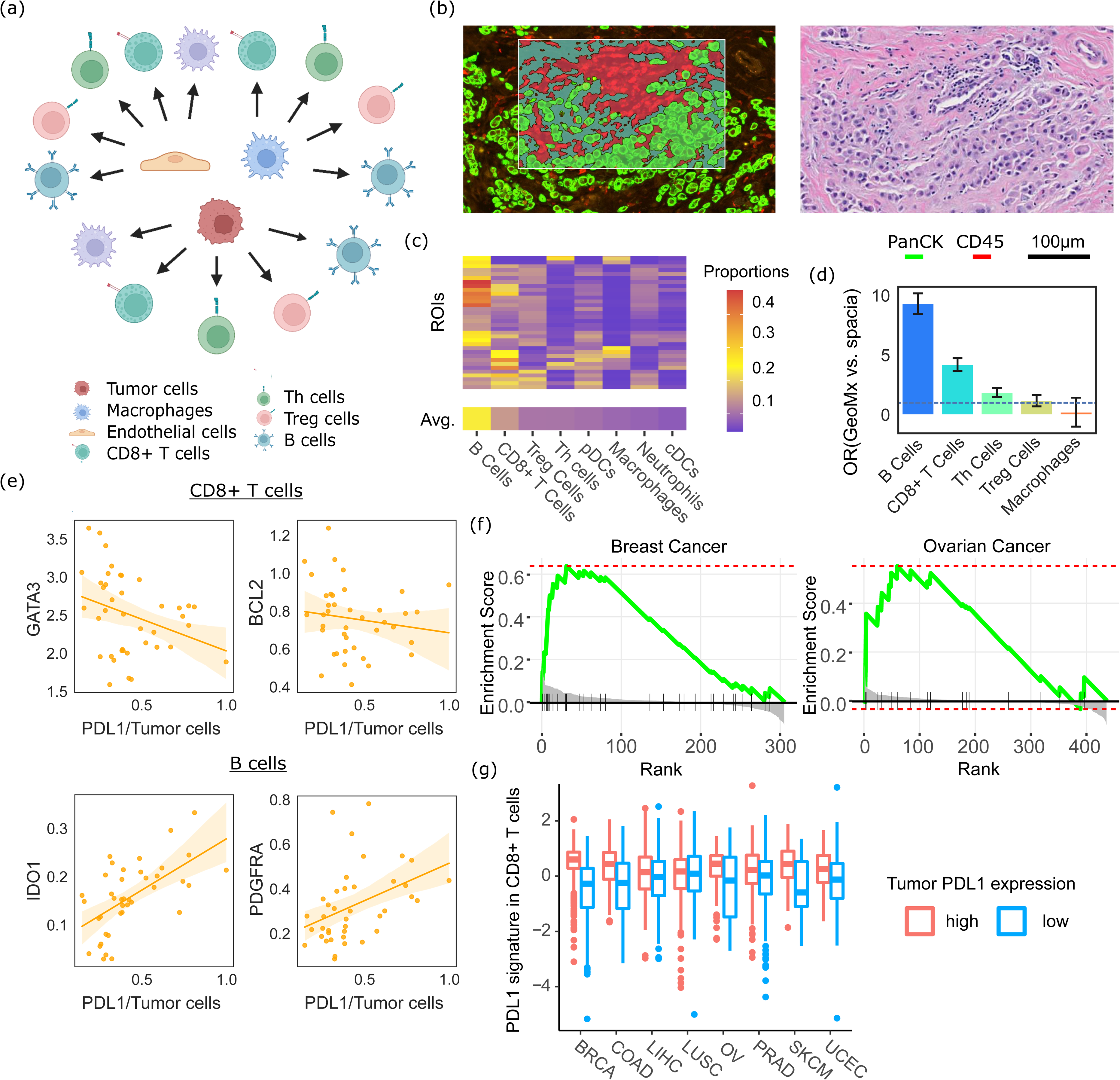
Spacia reveals *PDL1* downstream target genes. (a) Sending and receiving cell type pairs that were analyzed by spacia for the breast cancer MERSCOPE dataset. (b) Left: One example ROI of the GeoMX data; right: the region of the H&E slide corresponding to the same ROI. (c) Abundances of each type of immune cells in the GeoMx data, as predicted by BayesPrism. Cell types are ordered by their average abundance across the 40 ROIs. (d) Odds ratios showing the overlap between spacia-inferred *PDL1* downstream target genes and genes that are differentially expressed in each type of immune cells in the GeoMx data, comparing ROIs that are PDL1+ and PDL1-in the tumor cells. (e) Expression of several representative receiving genes that are in the overlap of (d) for B cells and CD8^+^ T cells, as a function of tumor cell *PDL1* expression, in the GeoMx data. The fitted curves between the X axis and the Y axis are shown as solid lines, with the shading denoting 95% CI. (f) Gene set enrichment analysis (GSEA) in the TCGA breast and ovarian cancer datasets to evaluate if the corresponding CD8-PDL1 signature genes are indeed among the top genes that are differentially expressed between PDL1+ and PDL1-patients. (g) Expression levels of the CD8-PDL1 signatures in CD8^+^ T cells in the TCGA samples, dichotomized by tumor cell *PDL1* expression.

To validate these observations, we generated a set of GeoMx data from eight treatment-naive breast cancer patients. Each patient was sampled for an average of five Regions of Interest (ROIs), resulting in a total of 40 ROIs with a mixture of tumor and immune cells. We defined the tumor and immune masks in each ROI (**Fig. 4b**, full images in **Sup. File 2**), and extracted mask-specific gene expression for the tumor cell and immune cell components. We deployed BayesianPrism ^37^ to dissect the immune cell gene expression into the gene expression of each immune cell type for each ROI. As **Fig. 4c** shows, B cells (19%) and CD8^+^ T cells (10%) are most abundant in these ROIs, followed by T_reg_ cells (6%) and T_h_ cells (6%), with macrophages accounting for the least proportions (4.5%). Next, we investigated genes in these immune cell types whose expression is positively or negatively correlated with the expression of *PDL1* in tumor cells. We then calculated the overlap between these genes and the *PDL1* downstream target genes that spacia inferred from MERSCOPE data, in terms of direction of expression regulation. Importantly, as BayesPrism’s authors suggested, BayesPrism is more accurate for cell types that are abundant in the tissue mixture (only B cells and CD8^+^ T cells achieve optimal accuracy according to their guidelines). Therefore, we focused this validation on the PDL1 target genes in B cells and CD8^+^ T cells. As **Fig. 4d** shows, there are statistically significant overlaps between GeoMx and spacia/MERSCOPE, for both B cells (Odds Ratio (OR)=9.3, Pval=0.01) and CD8^+^ T cells (OR=4.2, Pval=0.011). Note here that overlap is defined by a gene having the same inferred direction of regulation by PDL1 (both up-regulation or both down-regulation), in both the MERSCOPE and GeoMx data. As expected, the overlap is less pronounced for T_h_ cells (OR=1.87, Pval=0.14) and T_reg_ cells (OR=1.18, Pval=0.92), and there is no enrichment for macrophages (OR=0.21, Pval=0.37). Even though these two sets of PDL1-downstream genes were derived from two different technologies in two different cohorts of patients, the existence of a significant overlap speaks to the validity of spacia’s findings. Unfortunately, as the direction of regulation is critical for this analysis, we cannot perform comparison with the other five benchmark software applications, as none of them outputs the direction of regulation.

We showcase the top genes that are in these overlaps. In CD8^+^ T cells, both the spacia and the GeoMx analyses indicate that *BCL2* is inhibited by *PDL1* from tumor cells (**Fig. 4e**). *BCL2* is a key anti-apoptosis molecule^38^ and therefore our results suggest that *PDL1* promotes apoptosis in CD8^+^ T cells. We also found that *PDL1* down-regulates *GATA3* in CD8^+^ T cells (**Fig. 4e**), which supports the maintenance and proliferation of T cells downstream of TCR and cytokine signaling^39^. Both of these observations are in alignment with the well-known immuno-suppressive functions of *PDL1* for CD8^+^ T cells. For B cells, we found that *PDL1* of tumor cells up-regulates *IDO1* in B cells, in both the spacia and GeoMx analyses (**Fig. 4e**). The role of *IDO1* for B cells is less clear so far, but recent reports have linked *IDO1* with immuno-suppressive roles in B cells^40^. We also found that *PDL1* up-regulates *PDGFRA* in B cells (**Fig. 4e**). To our knowledge, there has been no literature on how the expression of *PDGFRA* in B cells is correlated with B cell functions. But we examined the breast cancer scRNA-seq data from Bassez *et al* ^41^, where data on various immune cell types pre- and post-anti-PD1 treatment were generated. These data show that *PDGFRA* expression in B cells decreased after anti-PD1 treatment (**Sup. Fig. 5a**, Pval=0.025), which suggests an activating role of *PDL1* for *PDGFRA*.

We also analyzed a breast cancer Xenium dataset (**Sup. Fig. 5b**) by spacia, which has *PDGFRA* in its gene panel. **Sup. Fig. 5c** showcases the spatial distributions of the cellular interactions that spacia inferred, which are local as expected. Reassuringly, spacia also inferred tumor cell *PDL1* to up-regulate *PDGFRA* in B cells (**Sup. Fig. 5d**), confirming our observations above. We systematically compared the genes that are shared between the MERSCOPE panel and Xenium panel (**Sup. Fig. 5e**), and observed that there is an overall concordance between the directions of *β*s from the spacia analyses on both the MERSCOPE and Xenium breast cancer datasets, for both B cells (OR=3.42) and CD8^+^ T cells (OR=1.9). Interestingly, we again observed that the concordance is higher for B cells, than for CD8^+^ T cells, reflecting our results in **Fig. 4d**.

### The Spacia-derived *PDL1* Signature in CD8^+^ T Cells Is Prognostic and Predictive

Next, we applied spacia to a set of pan-cancer MERSCOPE datasets, which include breast cancer, colon cancer, melanoma, lung cancer, liver cancer, ovarian cancer, prostate cancer, and uterine cancer (**Sup. File 2**). Here, we still focused on tumor cell-to-CD8^+^ T cell interactions. Spacia yields a set of downstream target genes for tumor-to-CD8^+^ T cell interactions in each cancer type, which we term the CD8-PDL1 signatures (**Sup. Table 4**). We first validated these gene signatures, by leveraging the RNA-seq data from The Cancer Genome Atlas Program (TCGA), for the same eight cancer types. We used BayesPrism ^37^ to dissect the TCGA RNA-seq data into cell type-specific gene expression in each patient sample. We investigated the differential gene expression in the CD8^+^ T cell components in the TCGA patients, between patients with high and low *PDL1* expression in their tumor cell components. Gene Set Enrichment Analysis (GSEA) ^42^ confirmed that the CD8-PDL1 gene signatures identified by spacia are indeed enriched in the top differentially expressed genes. **Fig. 4f** showcases the GSEA results for breast cancer (Pval=0.0007) and ovarian cancer (Pval=0.025). We also calculated a composite score as the weighted average of the genes in each CD8-PDL1 signature, with weights being the inferred *β*s by spacia, to reflect the overall impact of *PDL1* signaling on CD8^+^ T cells. For almost all cancer types, patients with higher *PDL1* expression in the tumor cells tend to have higher expression of the CD8-PDL1 signatures (**Fig. 4g**, Pval(liver cancer)=0.012, Pval(lung cancer)=0.69, and P values for all other cancer types <0.002). Overall, these results validate the CD8-PDL1 signatures inferred by spacia.

We evaluated the prognostic and predictive powers of the CD8-PDL1 signatures to determine whether they possess any translational value. These signatures can potentially reflect the actual impact of tumor cells’ *PDL1* signaling on T cells, which is a more direct measurement of *PDL1* signaling effectiveness and could reveal stronger biological signals compared with testing tumor *PDL1* expression alone. First, we tested the association between bulk tumor *PDL1* expression and patient overall survival for the eight cancer types of interest in the TCGA patients (**Sup. Fig. 6**). Somewhat counterintuitively, we observed that higher *PDL1* expression tends to predict better survival in these patients (Pval=0.016, all patients combined). One might expect higher *PDL1* expression leads to worse survival as it is known to suppress anti-tumor immune functions. However, inflammatory T-cell responses could incite tumor metastasis ^43^, thus PDL1 can alleviate the risk for metastasis through the inhibition of inflammation. While the underlying biological mechanisms of this effect are not within the scope of this work, we assessed whether the CD8-PDL1 signatures in CD8^+^ T cells better capture this phenomenon.

We performed Kaplan-Meier analyses for the CD8-PDL1 signatures in CD8^+^ T cells in all cancer types combined (**Fig. 5a**), and also for *PDL1* expression in the tumor samples as control (**Fig. 5b**). Here, we also subset the patients into patients with high and low CD8^+^ T cell infiltrations, based on the CD8^+^ T cell proportions from BayesPrism. Inspired by our prior observations ^44,45^, we posit that the correlation between CD8-PDL1/PDL1 and survival should be stronger when the tumors have more infiltrating CD8^+^ T cells (otherwise, *PDL1* signaling is irrelevant as there are few T cells to inhibit). For all cancer types combined, we found that high CD8-PDL1 signature expression indeed predicts better survival in the CD8^+^ T cell-high patients (Pval=0.004, OR=0.714, 95% CI=0.564-0.902), whereas there is no significant difference for the CD8^+^ T cell-low patients (Pval=0.34, OR=0.911, 95% CI=0.750-1.11) (**Fig. 5a**). On the other hand, the expression level of *PDL1* in tumor cells is less prognostic in both patient subsets (CD8^+^ T cell-high: Pval=0.26, OR=0.862, 95% CI=0.667-1.11; CD8^+^ T cell-low: Pval=0.16, OR=0.866, 95% CI=0.710-1.05) (**Fig. 5b**). We next limited our analyses to breast cancer only, and again confirmed that the CD8-PDL1 signature is a more robust biomarker than tumor *PDL1* expression for depicting the effect of *PDL1* signaling on patient survival (**Fig. 5c**: CD8-PDL1 signature, CD8+ T cell-high: Pval=0.0002, OR=0.337, 95% CI=0.190-0.597, CD8+ T cell-low: Pval=0.08, OR=0.627, 95% CI=0.371-1.06; **Fig. 5d**: tumor *PDL1* expression, CD8+ T cell-high: Pval=0.46, OR=1.35, 95% CI=0.616-2.94, CD8+ T cell-low: Pval=0.06, OR=0.593, 95% CI=0.347-1.01).

**Fig. 5.**
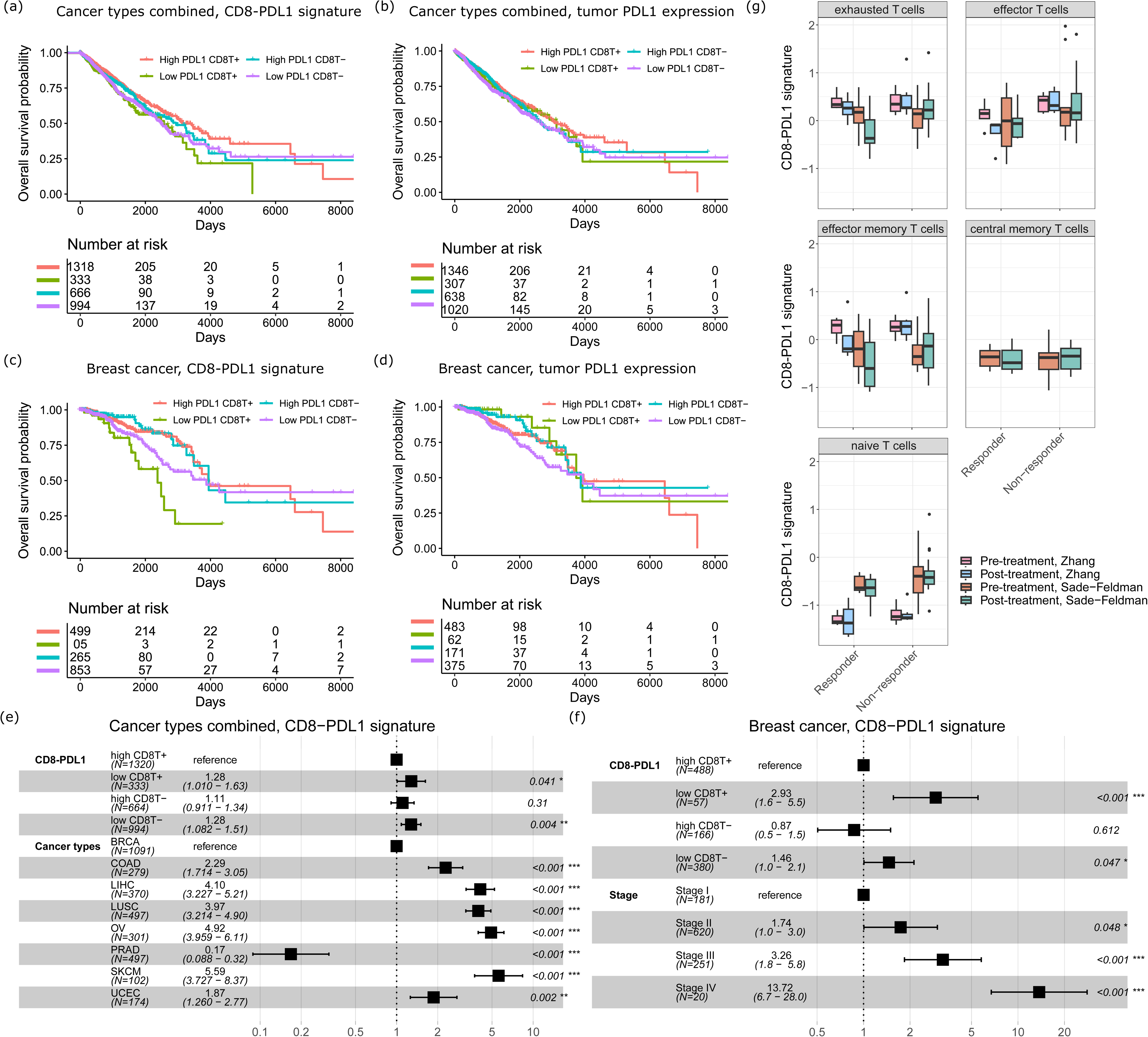
The PDL1-CD8 signature is prognostic and predictive. (a) Prognostic value of the PDL1-CD8 signature for overall survival of TCGA patients of all eight cancer types. (b) Prognostic value of tumor *PDL1* expression for overall survival of TCGA patients of all eight cancer types. (c) Prognostic value of the PDL1-CD8 signature for overall survival of TCGA breast cancer patients. (d) Prognostic value of tumor *PDL1* expression for overall survival of TCGA breast cancer patients. (e) Forest plot of hazard ratios from a Cox proportional hazard (CoxPH) model considering cancer types as confounding variables for patients of all eight cancer types. 95% CI is indicated by the whiskers. (f) Forest plot of hazard ratios from a CoxPH model considering cancer stages as confounding variables for breast cancer patients. 95% CI is indicated by the whiskers. (g) The CD8-PDL1 signature expression levels of CD8^+^ T cells in responders/non-responders and in pre-/post-treatment samples from the Zhang *et al* study and Sade−Feldman *et al* study.

We performed multivariate CoxPH analyses to adjust for the effect of confounding clinical covariates. In the pan-cancer cohort, the CD8-PDL1 signature is still significantly predictive of survival in the high CD8^+^ T cell patients after adjusting for the covariate of different cancer types (**Fig. 5e**, Pval = 0.041, Hazard Ratio=1.28, 95% CI=1.01-1.63). In breast cancer, the CD8-PDL1 signature is also still significantly prognostic in the high CD8^+^ T cell patients after adjusting for cancer stage (**Fig. 5f**, Pval < 0.0001, Hazard Ratio=2.93, 95% CI=1.6-5.5).

Next, we also investigated patients who were treated with anti-PD1/PDL1 therapies. We studied two anti-PD1/PDL1-treated cohorts, Sade-Feldman *et al* ^46^, which consists of 32 scRNA-seq datasets generated from peripheral blood of melanoma patients on anti-PD1 or anti-CTLA4+PD1 treatment, and Zhang *et al* ^47^, which consists of 11 scRNA-seq datasets generated from various tissue biopsies of breast cancer patients on anti-PDL1 treatment. Both cohorts contained responders and non-responders (as defined in the original works), and the biopsies were collected before and after treatment for scRNA-seq. We performed cell typing for the scRNA-seq data. For the CD8^+^ T cells in particular, we further classified them into exhausted, effector, effector memory, central memory, and naive T cells, according to marker genes from Sun *et al* ^48^ in order to conduct more fine-grained analyses. We observed that exhausted T cells, effector memory T cells, and effector T cells have higher overall expression levels of the CD8-PDL1 signature, with exhausted T cells having the highest expression (**Fig. 5g**). This is expected as the CD8-PDL1 signature measures the immuno-supppressive effect of PDL1 signaling in T cells. In responders, we observed that the CD8-PDL1 signature significantly decreased after treatment in exhausted (Pval=0.005, testing both cohort together) and effector memory T cells (Pval=0.048), while these changes were not observed in non-responders (Pval=0.23 and 0.61 respectively). This contrast indicates that the CD8-PDL1 signature is a candidate biomarker for measuring the effectiveness of immunotherapies that block the PD1/PDL1 axis.

Taken together, the CD8-PDL1 signatures that we defined from SRT data using spacia possess both prognostic and predictive values, which demonstrates the power of these new technologies for yielding novel insights of translational value.

### Differential Roles of γδ T Cells in Healthy Livers and Liver Cancers

We next applied spacia on two CosMX datasets, one generated from a healthy liver biospecimen and another one from a liver cancer biospecimen (**Fig. 6a** and **Sup. Fig. 7a**). We focused on a special class of T cells, γδ T cells, and characterized how these T cells impacted the transcriptomics programs of normal hepatocytes and malignant liver cancer cells. While γδ T cells are a minor class of T cells compared with αβ T cells, their abundance is much higher in the liver^49,50^. However, the exact roles of γδ T cells in the liver are not clear yet. In **Sup. Fig. 7b**, we first showed that γδ T cells surrounding hepatocytes have a generally much lower probabilities of truly interacting with the hepatocytes, while γδ T cells surrounding tumor cells have a much higher probabilities of interactions (T test Pval<1E-10). We also visualized the spatial distances between γδ T cells and their nearest healthy/cancerous hepatocytes (**Sup. Fig. 7c**, T test Pval=1.32E-5) and again observed that the distances are much smaller in the liver cancer specimen. These results suggested that γδ T cells likely have a different mode of action in the tumor context, as opposed to healthy livers.

**Fig. 6.**
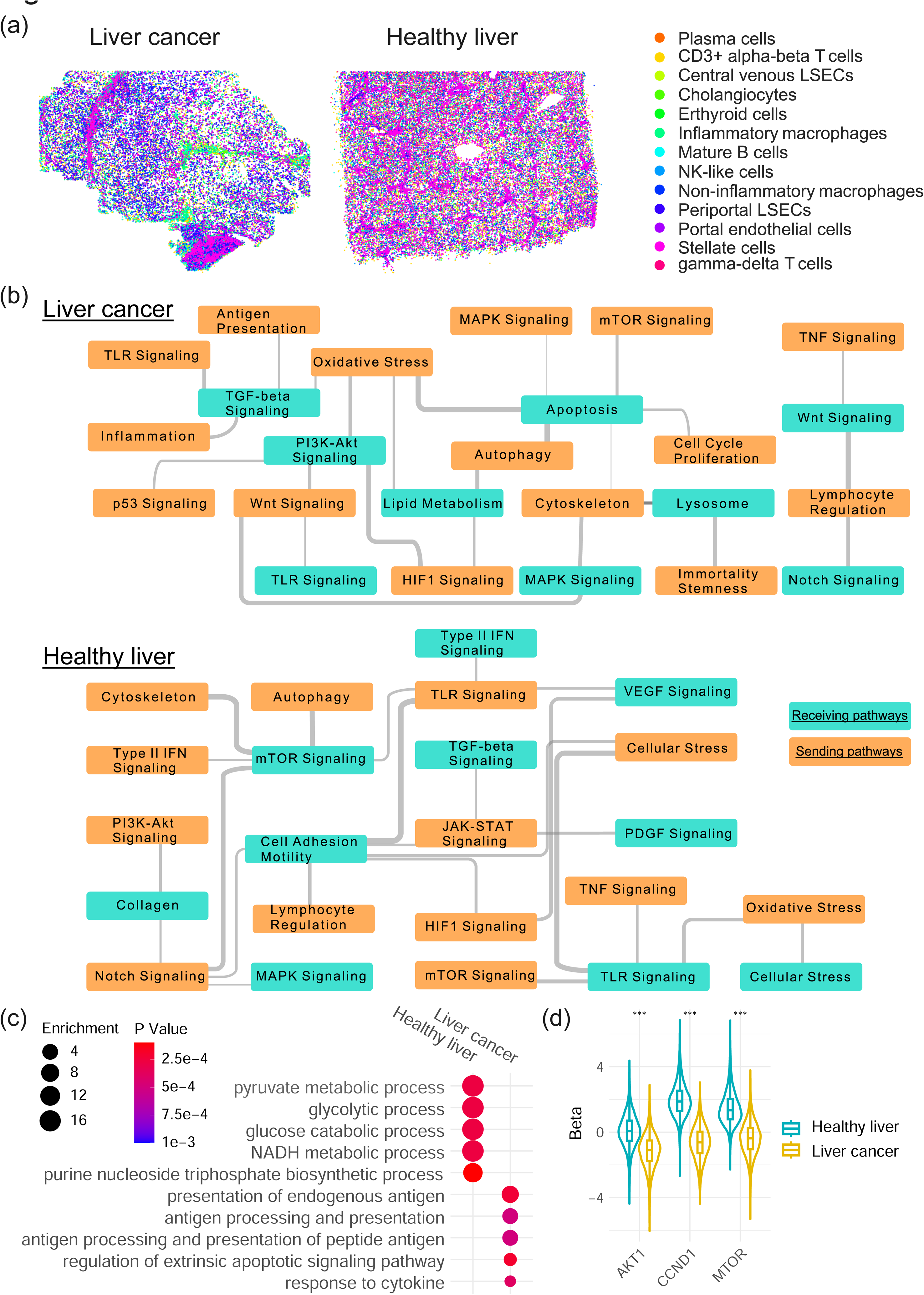
Spacia infers differential roles of γδ T cells in healthy livers and liver cancers. (a) The spatial distribution of stromal/immune cells in the liver cancer and healthy liver CosMx datasets. (b) The significant interactions between sending pathways in γδ T cells and receiving pathways in healthy/cancerous hepatocytes. Wider gray lines denote the more significant interactions. (c) Significant pathways from GO analyses of top receiving genes in healthy and cancerous hepatocytes, with *SQSTM1* in γδ T cells as the sending gene. (d) Heatmap showing the *β*s between the *SQSTM1* sending gene in γδ T and the *MTOR*/*AKT1*/*CCND1* receiving genes in healthy and cancerous hepatocytes. A positive *β* denotes a positive regulation relationship.

In **Fig. 6b**, we showcased the interactions that are active in each of the two samples at the pathway level, based on the recommended pathways by CosMX for these datasets (nanostring.com/resources/cosmx-smi-universal-cell-characterization-panel-gene-list), and found that the molecular pathways that are involved in the interactions between γδ T cells and healthy/cancerous hepatocytes are overall very different in the two contexts. For example, on the receiving side, apoptosis pathway shows up in the liver cancer cells, which reflects the cytotoxic role of γδ T cells. We also observed that the regulation of lipid metabolism pathway being impacted in the liver cancer cells. This might reflect an active modulation of the liver cancer cells, in response to the cytotoxic effect of γδ T cells, which recognize and are activated by lipid antigens ^51,52^. For the healthy liver, we observed the regulation of mTOR pathway in the hepatocytes, which has been linked with hepatocyte homeostasis and metabolism ^53^. This observation also resonates with the regulation of TGFβ/VEGF/PDGF pathways as shown in **Fig. 6b**, which are all important for hepatocyte homeostasis ^54^. On the sending side, we observed that the autophagy of γδ T cells impacts both the apoptosis of liver cancer cells and the mTOR pathway of healthy hepatocytes. As autophagy (especially selective autophagy) has emerged as a crucial regulator of T cell homeostasis, activation and differentiation^55^, our observations here seem to indicate that the homeostasis of γδ T cells is in a dynamic balance with the homeostasis of liver hepatocytes and impacts liver hepatocytes differently in different biological contexts.

We then examined individual genes and focused on *SQSTM1*, which is captured by the CosMx platform and is an important marker for selective autophagy ^56^. We examined the hepatocyte genes regulated by *SQSTM1* of γδ T cells and rank-ordered by the magnitude of regulation (*β*s). Through Gene Ontology (GO) analyses, we confirmed again that the top pathways enriched in healthy livers are almost all related to metabolism (**Fig. 6c**), while the top pathways enriched in liver cancer cells are enriched with apoptosis and antigen presentation, echoing our spacia analysis at the pathway level. We further examined *MTOR* and *AKT1*, in the mTOR pathway, as well as *CCND1*, which is a key gene that regulates hepatocyte homeostasis downstream of the mTOR pathway ^53^. (*PI3K* is not captured in the CosMx gene panel.) While *SQSTM1* in γδ T cells positively up-regulates *MTOR*, *ATK1*, and especially *CCND1* in healthy liver, we observed an opposite trend in liver cancer cells, as shown in **Fig. 6d**. The *β*s are statistically significantly different between the healthy liver and liver cancer conditions, for each of the three genes (K.S. Test, Pval<1E-10). Taken together, spacia analysis revealed that γδ T cell homeostasis participates in very different biological processes through their interactions with hepatocytes in the healthy and cancerous contexts.

## DISCUSSION

We developed spacia to fulfill the unmet gap in detection of cell-to-cell and gene-to-gene interactions from SRT data. Our work has five major conceptual advantages: (1) spacia achieves single cell resolution detection of CCCs, (2) spacia is not limited to prior interaction datasets and (3) not limited to ligand-receptor relationships, (4) spacia addressed the high false positive rate and low specificity issues commonly seen in prior works, and (5) spacia explicitly infers the multiple-sender-to-one-receiver relationships in SRT data through a novel Bayesian multi-instance learning framework. We proved that the quality of modern commercialized single cell resolution SRT technologies already enables the detection of CCCs with appropriate statistical models like spacia, ensuring the broad applicability of our work. We comprehensively tested spacia on simulation data and real data from diverse biological conditions, and very importantly on data from all three single cell resolution SRT technologies: MERSCOPE, CosMX, and Xenium. Practically, spacia can be executable on such SRT data that yield >50,000 cells, while we have to aggressively down-sample the number of cells to execute the other SRT-based CCC inference tools that we benchmarked to avoid memory overflow.

Whereas most existing CCC inference tools work by examining data against a known database, spacia focuses on the more general concept of CCC through accurate *in situ* modeling. Spacia achieves this through organically integrating the rich spatial and transcriptional information, which allows it to minimize the number of assumptions and arbitrary parameters. While this approach does not rely on existing knowledge, it offers an unbiased, independent interrogation based on first principles. Furthermore, it enables discoveries that complete known CCC pathways and uncover new pathways in a cellular context specific manner. It is important to note that the target genes identified by spacia, through modeling of transcriptional regulation as a function of sending gene expression, could be either direct targets that are within the core of the interacting pathways or represent more downstream and secondary effects. This could be an advantage or disadvantage depending on one’s vantage point. At this time, cross-referencing to existing databases could be helpful for the interpretation of the identified CCCs and distinguishing between “direct target” *vs.* “indirect target”, and allows for further interrogation of the regulatory relationships between genes in existing pathways. Therefore, we propose a paradigm of unbiased search of CCCs followed by filtering through prior knowledge in spacia, in contrast to those of prior works, such as CellPhoneDB, whose performance is limited to prior knowledge from the very beginning.

The core of spacia is a fully integrated Bayesian MIL model. Due to the complexity of learning the multiple-to-one functions, MIL problems are much more difficult than typical machine learning problems. Spacia’s MIL model allows solving this difficult mathematical problem in a graceful manner. Whereas existing MIL methods mainly focus on predictive performance ^57–59^, our two-tier MIL approach enables concurrent identification of primary instances (the sender cells responsible for the reaction in the receiver cell) and elucidation of the relationships between bags and instances (βs). This technical innovation is important for both the fields of MIL in general and the specific application of finding CCC in SRT data. Critically, in the era of data explosion in biomedical research, MIL can offer elegant solutions to disentangle complex interactions and large, diverse data from various fields. For example, in biochemistry, one might be interested in how the many conformations of a chemical compound are related to its bioactivity ^60^. We envision that our work will propel the wide adoption of MIL in biomedical research, by providing a success case and also by providing a model that can be further improved upon.

Admittedly, one major limitation associated with Bayesian MIL model is that spacia’s runtime quickly increases with increasing sample sizes, namely numbers of bags, instances and gene features, to attain convergence. Our current spacia already leverages C++ programming to enhance the computational efficiency, implemented by the Rcpp package in the R language. In our future iterations, we will deploy mean-field variational inference to approximate the joint posterior and to further enhance computational efficiency, where our Bayesian formulation will allow a close-form solution in each step of the mean-field algorithm.

Despite the exciting results presented in our study, mapping CCC is still challenging. Perhaps one of the biggest challenges is associated with the low signal-to-noise ratios (SNRs) of the SRT data. The inaccuracy in cell segmentation is a major contributor to this low SNR. Most, if not all, high-resolution SRT technologies generate the raw expression counts at subcellular or even pixel levels. To aggregate such data to single cell levels, cell boundaries are usually created from the matching H&E staining or fluorescence images *via* segmentation techniques. However, improper cell segmentation can lead to serious errors. For example, two cells that are close by (potentially two cells that are interacting) can be segmented as the same cell. In this scenario, two genes that are interacting with each other from two different cells can be mistaken as a pair of genes that are transcriptionally linked within the same cell. However, the field has seen continuous improvement in the cell segmentation procedure and other pre-processing procedures of SRT data, either in academically generated tools ^61–64^ or commercially available software. We expect such caveats to become less pronounced with future iterations of SRT technologies and the corresponding bioinformatics analysis software.

As discussed above, we recommend SRT users to visually assess the image segmentation quality to determine if the cell types of interest have been accurately segmented out. In addition, we suggest users to deploy de-noising software, such as Sprod ^62^, to increase the SNRs of the expression data from SRTs, and use dimension-reduction/visualization tools ^65–67^ to assess the overall clustering quality of the cells.

In summary, we built a general and principled framework for the analysis of CCCs from SRT data that can be applied to a vast number of biological systems. More broadly, when coupled with remarkable experimental and analytical advances in single cell and spatial approaches ^68–77^, spacia will enable us to understand how complex cell states arise from communications in the local cellular community and to move towards holistic models and theories of entire organisms.

## METHODS

### Simulation data creation

Simulated datasets were generated with the following steps. First, 8,000 cells were generated in a two-dimensional space of 2,000 x 2,000 units. These cells were classified into three types, with the receivers forming the core of several blobs, senders lining the perimeters, and non-interacting cells filling the space between blobs. Senders were divided into two categories: primary senders and non-primary senders based on their distances to a given receiver (distance cutoff = 50). Next, we simulated expression data (50 genes) associated with sender and receiver cells. Expression of each gene of the sender cells was generated with normal distribution (mean = 0, s.d. = 1). The first 5 or 10 genes (in two simulation settings) were designated as truly interacting genes, in two different settings, while the remaining genes were designated as non-interacting genes.

Expression for receivers was generated as a weighted sum of gene expression from their primary senders, with weights (βs) generated from uniform distributions. The uniform distribution ranges from 10 to 20 for the truly interacting genes and 0 to 1 for the other genes. The signs of these weights were randomly assigned to be positive or negative to model upregulation and downregulation of receiving genes by sending genes. Lastly, we added noise to the receiver genes, with noises sampled from a normal distribution (mean = 0, s.d. = 0.5).

### MERSCOPE data pre-processing and annotation

The pan-cancer MERSCOPE datasets consisting of 8 different cancer types were downloaded from the publically available MERFISH FFPE Human Immuno-oncology Data Set. The R Seurat package was used to load the “cell_by_gene.csv” file for each MERSCOPE experiment to create a Seurat object for downstream processing and analysis. To ensure we only retain high-quality cells, each experiment was subset so that cells with more than 100 total counts were kept. After the initial clustering and annotation, we subset the T cell population and the epithelial population and performed a second round of the standard workflow for each population to identify the fine-grained clusters for the T cell subpopulations and to segregate epithelial cells into tumor epithelial cells and normal epithelial cells. The final version of the annotated clusters in the UMAP space and the average gene expression of cell type markers in each cluster can be found in **Sup. File 2**.

### Benchmarking with existing CCC software

CellPhoneDB: Normalized counts and cell type labels were converted into CellPhoneDB compatible TXT files. A microenvironments file was generated to limit the predictions to stromal-tumor interactions only. CellphoneDB 3.1.0 was run using method statistical_analysis, options --counts-data hgnc_symbol, --pvalue 1, and --threads 24 options. Since CellPhoneDB does not designate sending and receiving cells, they were defined by which of the interacting genes was labeled as receptor. For each interaction, the cell type with the gene labeled as receptor was designated as the receiving cell. Interactions where both or neither genes were labeled as receptor were discarded due to ambiguity of the direction of interaction. For consistency with spacia results, results with smooth muscle and normal epithelial cells as sending cells were removed, and only interactions with tumor cells as the receiving cell were kept. Interactions with Pval > 0.05 were removed.

CellChat: CellChat 1.5.0 was used with the default CellChatDB.human ligand-receptor interaction database. Functions *identifyOverExpressedGenes* and *identifyOverExpressedInteractions* were run with default options; *computeCommunProb* was run with type = “truncatedMean” and trim = 0.05; *filterCommunication* used min.cells = 10, and *computeCommunProbPathway*, *aggregateNet*, and *subsetCommunication* were all run with default options. For the outputs, “source” was considered equivalent to spacia’s sending cell, “target” equal to receiving cell, “ligand” as sending gene, and “receptor” as receiving gene. For consistency, results with smooth muscle and normal epithelial cells as sending cells were removed, and only interactions with tumor cells as the receiving cell were kept. Interactions with Pval > 0.05 were removed.

COMMOT: COMMOT version 0.0.3 was used in the analysis. The function commot.pp.ligand_receptor_database was run with options database=CellPhoneDB_v4.0, and species=human to generate the ligand-receptor dataframe. Function commot.tl.spatial_communication was run with option database_name=cpdb, dis_thr=500, heteromeric=True, and pathway_sum=True to infer spatial communication. Results were filtered by sending and receiving cell types to match that of spacia, and cell type level results were generated using the funtion commot.tl.cluster_communication with option database_name=cpdb. For visualization, commot.pl.plot_cell_communication was run with options: database_name=cpdb, plot_method=cell, scale=0.03, ndsize=10, background=cluster, background_legend=True, cmap=Alphabet, and clustering=cell type labels.

SpatialDM: SpatialDM version 0.1.0 was used. Weight matrix was calculated using sdm.weight_matrix with options l=50, cutoff=0.2, and single_cell=True. The value for l was found according to the method described in the authors’ tutorial. Following the tutorial, sdm.extract_lr was run with species=human, and min_cell=3, then sdm.spatialdm_global was run using method=both, and sdm.sig_pairs with method=permutation. Local results were calculated by running functions sdm.spatialdm_local with option method=both, and sdm.sig_spots with options method=permutation and fdr=False.

SpaTalk: SpaTalk version 1.0 was used. The function createSpaTalk was run with options species=Human, if_st_is_sc=T, and spot_max_cell=1 to generate the SpaTalk object. For the analysis, first the function find_lr_path was run using the default ligand-receptor pairs and pathways, then function dec_cci was run using the same sending and receiving cell types as those in the spacia analysis.

### Validation with PDMR data

Bulk prostate cancer PDX RNA-sequencing data were downloaded from The NCI Patient-Derived Models Repository (PDMR), NCI-Frederick, Frederick National Laboratory for Cancer Research, Frederick, MD (pdmr.cancer.gov). Disambiguate ^19^ was used to segregate the PDX RNA-seq data into the human and mouse components. CIBERSORTx was used to further deconvolute the mouse stromal/immune component into cell type specific expression values. The CIBERSORTx web portal (cibersortx.stanford.edu) was used and the signature matrix for CIBERSORTx was created on the web portal using the expression matrix and cell typing results from the prostate cancer MERSCOPE data. To comply with the computational limits of the web server, for cell types with large numbers of cells, 5,000 cells were randomly sampled to create the reference sample file. CIBERSORTx was then run in high resolution mode for the PDMR data using default options.

### EMT process calculation in the prostate MERSCOPE dataset

We define the “EMT activation potential” scores as the sum of the strengths of EMT-induction signals from nearby sender cells for each tumor cell. This is calculated with the following steps. The β values from the sender-to-tumor cells spacia results are filtered to keep those with sending genes included in a list of secreted factors that have been reported to impact EMT (*HGF*, *WNT3A, WNT5A*, *FGF2*, *IL6*, *CXCL8*, *FGF1*, *TGFB1*, and *TGFB2*), and receiving genes being EMT upstream regulators that have some prior evidence of being impacted by these factors (*JAK1*, *AKT2*, *SMO*, *CTNNB1*, *SMAD2*, and *NFKB2*) ^25–33^. For each sender cell and each sending gene, a set of scores are calculated by multiplying the β of each sending gene-receiving gene pair with the corresponding expression of the sending gene. These scores, one for each receiving gene, are averaged. The averaged scores for all sending genes are further averaged to arrive at a composite score to be assigned as the EMT activation potential of this sender cell. Finally, an EMT activation potential score for each tumor cell is computed by the weighted sum of the EMT activation potentials of the sender cells in each tumor cell bag with the weights being the primary instance probabilities of the senders.

The “EMT score” of a tumor cell is defined to be the mean expression of the EMT marker genes from Gorgola *et al* ^78^: *FN1, TWIST1, SNAI1, SNAI2, ZEB1, TGFB1, TGFB2,* and *CTNNB1*, which represents the activity level of EMT in that tumor cell. We performed additional analyses to validate that this EMT score is valid in our scRNA-seq datasets (**Sup. File 2**).

### Lineage score calculation in the prostate cancer MERSCOPE dataset

We collected gene markers from Deng *et al* ^36^ to calculate the lineage plasticity scores for the basal, neuro-endocrine and stem-like lineages. We were not able to investigate the luminal lineage, as the most important canonical markers such as *AR* and *KLK3* were missing. For the basal lineage, *TP63*, *CAV1*, and *LAMB3* were used. For the neuro-endocrine lineage, *EZH2* and *NCAM1* were used. For the stem-like lineage, *KIT*, *LGR5*, and *LGR6* were used.

### Prostate patients for scRNA-seq

The study was performed following protocols approved by the Institutional Review Board of the University of Texas Southwestern Medical Center. There are two studies from which we obtained patient biopsies. STU 072010-098: Tissue Procurement and Outcome Collection for Radiotherapy Treated Patients & Healthy Participants, and STU062014-027: Phase I Clinical Trial of Stereotactic Ablative Radiotherapy (SABR) of Pelvis and Prostate Targets for Patients with High Risk Prostate Cancer. We obtained a total of 21 androgen deprivation (ADT)-treated patients, 4 untreated prostate cancer patients, and 4 healthy donors. The scRNA-seq experiments were done in Dr. Douglas Strand’s lab following the protocol in Henry *et al* ^79^. Briefly, we performed a 1 hour digestion with 5mg/ml collagenase type I, 10mM ROCK inhibitor, and 1mg DNase. 3’ GEX barcoding was performed on a 10X controller and sequencing was performed on an Illumina NextSeq 500 sequencer. These 29 scRNA-seq datasets were processed through QC, integration, and cluster annotation prior to the analysis. We utilized the R DoubletFinder package and followed the recommended workflow for filtering the doublets or multiplets in the data. For the joint analysis of multiple patient scRNA-seq datasets, we followed the Seurat integration vignette. Each dataset was processed through the *ScaleData* and *RunPCA* functions. The anchors for integration were found with the “rpca” mode of the *FindIntegrationAnchors* function.

Utilizing these anchors, we proceeded with the *IntegrateData* function for the integration of the 29 scRNA datasets. Initially, the clusters were automatically annotated with the R SingleR package and the annotated dataset from Song *et al* ^80^.

(1) The patients were dichotomized into EMT+ and EMT-groups with the following steps. First, the EMT level of each tumor cell is defined as above in the prostate cancer MERSCOPE dataset. The global median EMT level is calculated from all the tumor cells. Then, for each patient, the percentage of tumor cells with higher EMT levels than the global median cutoff is calculated. The patients were divided into a EMT+ group and a EMT-group based on the median of these percentages. (2) The calculation of the EMT activation potential in the scRNA-seq data follows the same method as in the MERSCOPE dataset, except for that we calculated the activation potential for each sender cell, and did not aggregate to receiver cells through spatial averaging (as these are not SRT data). (3) Finally, the lineage scores for basal, neuro-endocrine, and stem-like were calculated as in the MERSCOPE dataset, utilizing the complete marker set from Deng *et al* ^36^.

### Breast cancer patients for GeoMx

The study was performed following protocols approved by the Institutional Review Board of the University of Texas Southwestern Medical Center. The IRB Protocol Number is STU2018-0015: Pre-surgical trial of letrozole in post-menopausal patients with operable hormone-sensitive breast cancer. Formalin-fixed, paraffin-embedded (FFPE) tumor tissues were obtained from diagnostic core needle biopsies from postmenopausal patients with stage I to stage III operable ER+/HER2-breast cancer enrolled in a clinical trial (UT Southwestern SCCC-11118).

To perform the GeoMx® assays, 5 µm FFPE sections were mounted on charged slides, baked, and prepared on the Leica Biosystem, following the manufacturer’s automated slide preparation user manual. After hybridization with RNA probes that are conjugated to barcoded oligonucleotide tags with an ultraviolet (UV) photocleavable linker and staining with fluorescent morphology markers consisting of pan-cytokeratin (epithelial and tumoral regions), CD45 (immune cells), SYTO13 (nuclear) and Ki67 (proliferation marker), the slides were loaded onto a GeoMx® digital spatial profiler (DSP, NanoString Technologies) and scanned. After labeling, the tissues were imaged, and we selected specific regions of interest (ROIs) sized 222 x 354.6 µm after consultation with a pathologist. Each ROI was further segmented into Areas of Illumination (AOI) based on morphological features. Subsequently, the selected areas were exposed to UV light, and the barcoded oligos were released, aspirated, and dispensed into a collection plate for library construction for next-generation sequencing (NGS).

GeoMx NGS libraries were prepared according to the manufacturer’s instructions. Briefly, after the collection of the probes was completed, aspirates in the collection plate were dried at 65°C for 1 hour in a thermal cycler with an open lid and resuspended in 10 µL of nuclease-free water. 4 μL of rehydrated aspirates were mixed with 2 μL of 5×PCR Master Mix and 4 μL of SeqCode primers. PCR amplification was then performed with 18 cycles. The indexed libraries were pooled equally and purified twice with 1.2×AMPure XP beads (Beckman Coulter). The final libraries were evaluated and quantified using Agilent’s High Sensitivity DNA Kit and Invitrogen’s Qubit dsDNA HS assay, respectively. Total sequencing reads per DSP collection plate were calculated using the NanoString DSP Worksheet. The libraries were sequenced using 38 bp paired-end sequencing (PE 38) on an Illumina NovaSeq 6000 system with a 100-cycle S1 kit (v1.5). FASTQ files were processed into digital count conversion digital files using Nanostring’s GeoMx NGS Pipeline software. Quality control, data filtering and normalization (Q3) were performed using the GeoMx DSP Data Analysis suite.

### BayesPrism bulk expression data deconvolution

The BayesPrism R package was installed according to its authors’ instructions at github.com/Danko-Lab/BayesPrism. The histology-matching MERSCOPE dataset was used as the single cell reference in order to maximize consistency with spacia’s results. BayesPrism was run using the *run.prism* function with default options. The cell type assignments we produced for the MERSCOPE datasets were used as the “cell.type.labels”. Cell type fractions were extracted using the *get.fraction* function. Predicted cell type-specific expression was extracted by running the *get.exp* function on each cell type.

### The Cancer Genome Atlas Program (TCGA) RNA-seq data analysis

TCGA data of eight cancer types that are of the same histologies as the MERSCOPE data were downloaded from Broad GDAC Firehose. These include: BRCA, COAD, LIHC, LUSC, OV, PRAD, SKCM, and UCEC. We only considered primary tumor samples of the TCGA cohort since all MERSCOPE datasets are from primary tumor samples. As we expected, inclusion of the TCGA metastatic samples (data not shown) in all downstream analyses yields similar results but with less statistical strength. BayesPrism was then run with default options, using the corresponding MERSCOPE data as the reference.

To define the CD8-PDL1 signature for each cancer type, we extracted, from spacia’s results, all receiving genes with *b* < 0, *b* Pval < 0.01, and *β* Pval <0.1. The genes of each cancer type were further filtered to keep those that have appeared in the gene list from breast cancer and at least two other cancer types. This was done due to our promising analysis results in breast cancer in **Fig. 4a-e**. The genes that passed all filters were designated as the CD8-PDL1 signature for each cancer type. The R Fast Gene Set Enrichment Analysis (fgsea) package was used to perform GSEA using the CD8-PDL1 signatures for the TCGA BayesPrism-dissected expression data. GSEA was run using the *fgsea* function with options eps=0, minsize=10, and scoreType=“pos”. The results were plotted using the *plotEnrichmentData* function from the fgsea package and the R ggplot2 package.

### TCGA patient survival analysis

The same TCGA datasets and CD8-PDL1 signature genes as above were used. For each patient sample in the TCGA datasets, the CD8-PDL1 signature expression level was calculated as the dot product of the normalized and log-transformed gene expression values of the BayesPrism-dissected CD8^+^ T cells and the *β* values of the genes in the corresponding CD8-PDL1 signature gene set. Survival analyses were performed using the *survfit* function from the R survival package with the CD8-PDL1 signature or tumor *PDL1* as the covariate. The Kaplan-Meier curves were plotted using the *ggsurvplot* function from the R *survminer* package. For the forest plots, the *coxph* function from the R *survival* package was used to fit Cox proportional hazards regression models with CD8-PDL1 signature and cancer type or stage as covariates. The forest plots were then generated using the *ggforest* function from the *survminer* package.

### Anti-PD1/PDL1 scRNA-seq data analyses

The Zhang *et al* ^47^ study originally identified naive (CD8Tn), effector (CD8Teff), and effector memory (CD8Tem) CD8^+^ T cell subsets in the patient peripheral blood, while the Sade-Feldman *et al* ^46^ study originally classified CD8Tem, central memory (CD8Tcm), exhausted (CD8Tex) subtypes. To homogenize classification and nomenclature, we loaded each dataset as a Seurat object to recluster and tried to identify all CD8^+^ T subtypes in both studies. Apart from the 20 dimensions used in FindNeighbors and FindClusters, all default parameters were used. Utilizing the gene marker sets from Sun *et al* ^48^, we annotated 28 clusters in the melanoma dataset and 13 clusters in the breast cancer dataset, compared to 4 and 6 clusters originally. All CD8^+^ T subtypes were identified, except for CD8Tcm in the Zhang *et al* dataset. The CD8-PDL1 signature score was calculated as in the TCGA data analysis, except that we have the actual CD8^+^ T cell gene expression in these scRNA-seq data, as opposed to BayesPrism-dissected expression for the TCGA samples.

### Statistics & reproducibility

Computations were performed in the R (3.6.3 and 4.1.3) and Python (3.7 and 3.8) programming languages. All statistical tests were two-sided unless otherwise described. Spacia was run in the PCA mode (see our documentation) for all our analyses in this work. Unless otherwise stated, the spacia results were filtered to keep those interactions satisfying *b* < 0, *b* Pval<1x10E-30, and *β* Pval<0.01. For pre-processing of all SRT and scRNA-seq data, we loaded the original data into Seurat objects and analyzed them through a standard workflow, using *NormalizeData*, *FindVariableFeatures*, *ScaleData*, *RunPCA*, *FindNeighbors*, *FindClusters*, and *RunUMAP* functions, before other custom operations. We executed the pipeline with default parameters except for dims = 1:20 for *FindNeighbors* and *RunUMAP* in the prostate cancer scRNA-seq datasets to capture sufficient variability in the principal components. GOrilla ^23,24^ pathway enrichment analyses were performed by inputting the genes on the MERSCOPE gene panel into GOrilla, where the genes were filtered to keep only those that satisfy *b* < 0 and sorted by *β* values for each sending cell type (tumor cells are the receiving cells). The Kernel Density Estimation for the visualization of spatial EMT patterns was calculated from the *gaussian_kde* function from the SciPy package with bandwidth less than the silverman factor, to demonstrate robustness from the spatial noise.

### Data and code availability statement

The MERSCOPE datasets were downloaded from vizgen.com/data-release-program. The breast cancer Xenium dataset was downloaded from www.10xgenomics.com/resources/datasets. The PDX RNA-seq datasets were downloaded from pdmdb.cancer.gov/web/apex/f?p=101:41:0:. The TCGA data were downloaded from gdac.broadinstitute.org. The scRNA-Seq datasets by Zhang *et al* ^47^ and Sade-Feldman *et al* ^46^ were accessed *via* Gene Expression Omnibus with accession numbers GSE169246 and GSE120575, respectively. The scRNA-Seq datasets by Bassez *et al*^41^ were accessed from biokey.lambrechtslab.org. The CosMx data are available from nanostring.com/products/cosmx-spatial-molecular-imager/ffpe-dataset/human-liver-rna-ffpe-data set. The prostate cancer scRNA-seq data that we generated were archived at doi.org/10.5281/zenodo.8270765. The breast cancer GeoMx data that we generated were archived at github.com/yunguan-wang/Spacia/data. The spacia software is available at the Database for Actionable Immunology ^76,81,82^ (dbai.biohpc.swmed.edu, Tools page).

## ACKNOWLEDGMENTS

We acknowledge Dr. Shijia Zhu for creating **Fig. 1c**.

## AUTHOR CONTRIBUTION STATEMENT

J.Z., Y.W., W.C. contributed to all bioinformatics analyses and implemented the software. A.M., J.G., R.H., P.M., D.S., N.D. generated the prostate cancer scRNA-seq data and/or provided critical insights in analyses. M.Z., F.N., L.G., N.U., A.H., C.A, generated the breast cancer GeoMX datasets. M.Z., S.Z., Z.Z., D.C. performed *in situ* ISS analyses on breast cancer mouse models. F.W. created the readthedocs website. G.X., W.X., Y.X., T.W. developed the initial concept and provided resources for the study. All authors wrote the manuscript.

## FIGURE AND TABLE LEGENDS

**Sup. Fig. 1.**
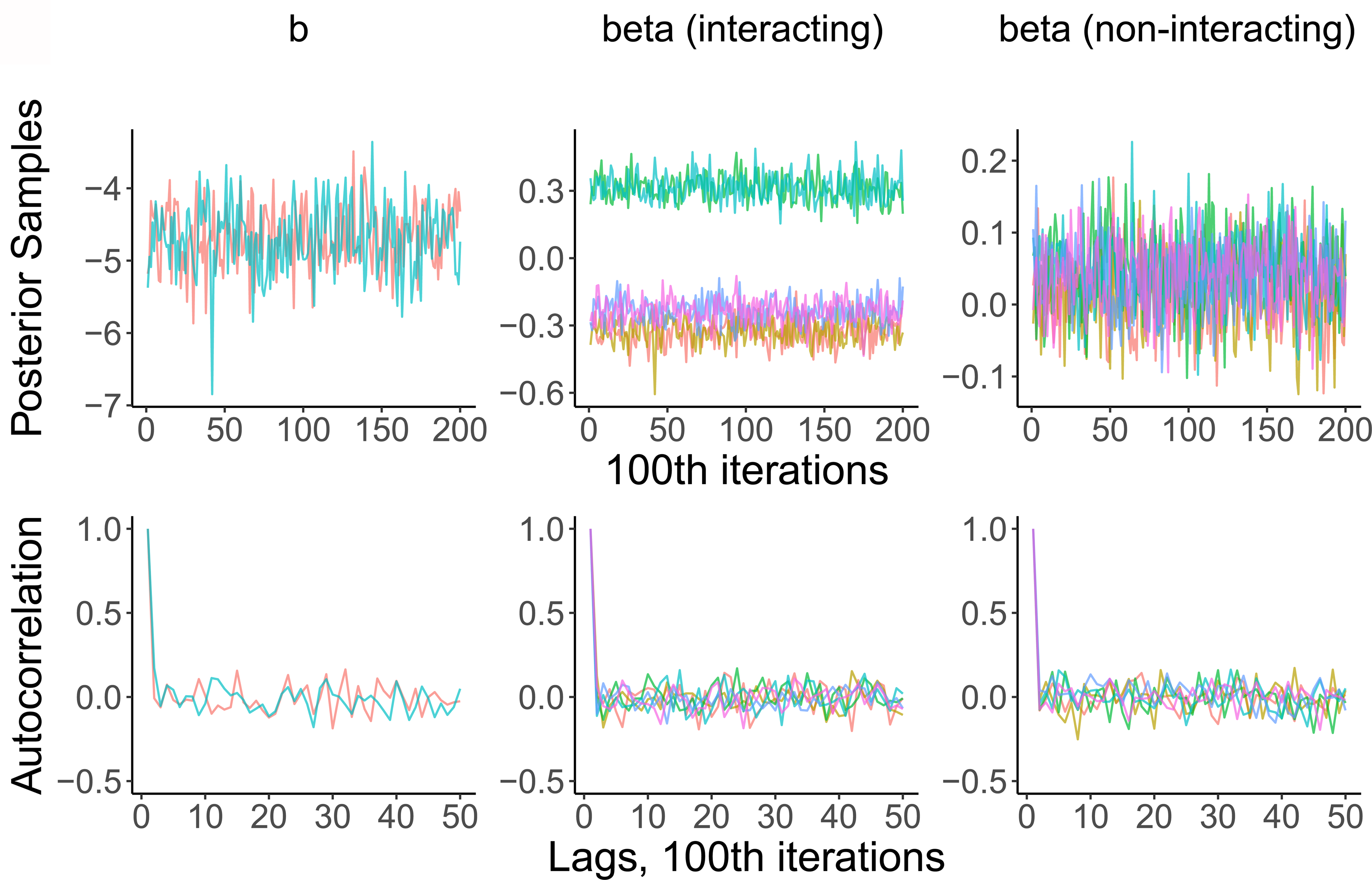
Traceplots and autocorrelation plots prove convergence and stability of the MCMC estimation process in spacia. Only MCMC iterations after the burn-in period are shown.

**Sup. Fig. 2.**
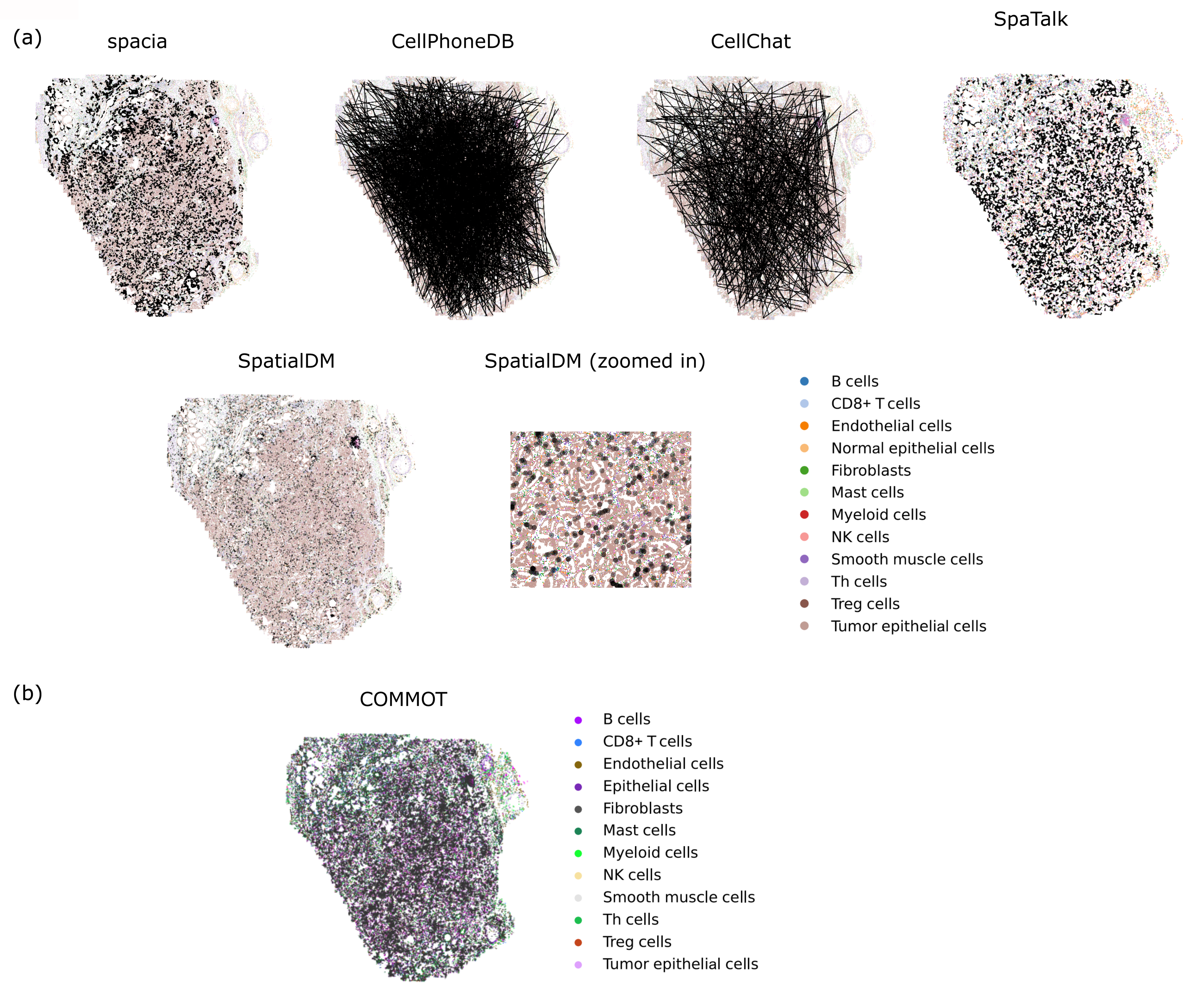
Visualizing the CCCs predicted by (a) spacia, CellPhoneDB, CellChat, SpatialDM, SpaTalk, and (b) COMMOT in their spatial context and at the single cell level. To reduce cluttering, for each sender-receiver cell type pair, 10 connections were selected at random and visualized for CellPhoneDB’s results, and 500 connections were selected at random and visualized for CellChat’s results. The interactions refer to the overall potential for a given pair of cells to interact, taken from the output statistics of each CCC inference software. COMMOT was separately listed in panel (b) since the internal data used by COMMOT for the spatial plot were not readily accessible, and the plot was generated by COMMOT’s own plotting function instead. For SpatialDM, the software did not output specific interactions between individual cells, but rather, only gave a score for each cell that participated in the CCCs without knowing the interaction partners. The black dots refer to this score.

**Sup. Fig. 3.**
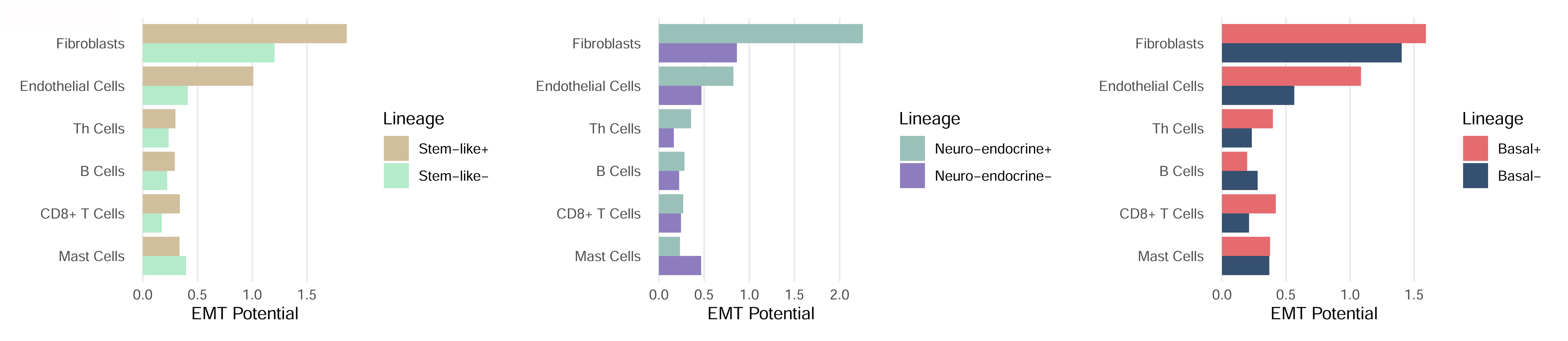
EMT activation potentials of each sending cell type by patient groups, dichotomized according to their lineage plasticity levels, for each lineage, in the prostate cancer cells.

**Sup. Fig. 4.**
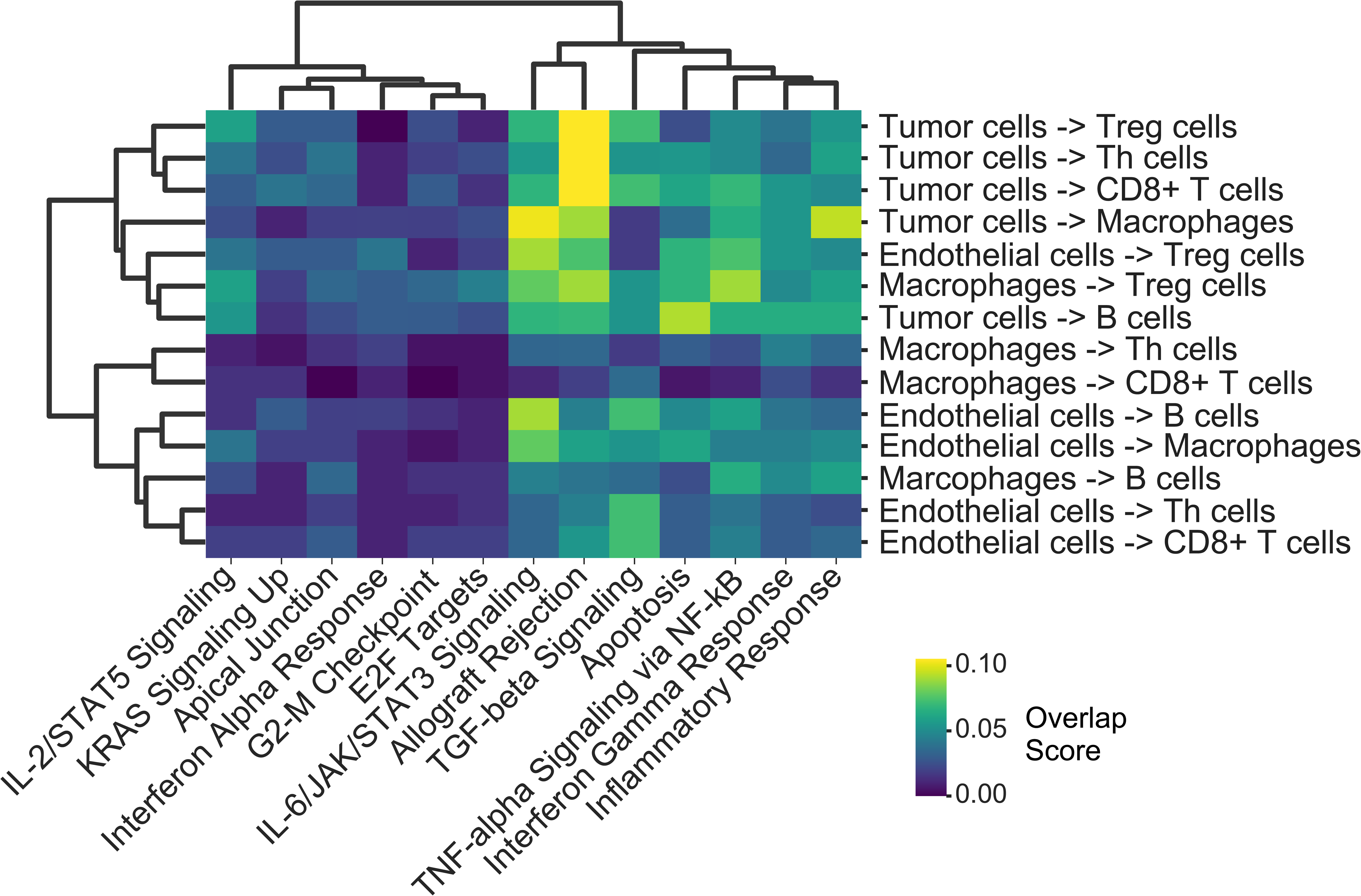
Overlap of *PDL1* regulated genes with the MsigDB Hallmark pathways. Genes in receiver cells that were significantly regulated by *PDL1* from sender cells were intersected with the genes of each MsigDB Hallmark pathway. The ratio values represent the proportions of all genes of each pathway that are intersecting.

**Sup. Fig. 5.**
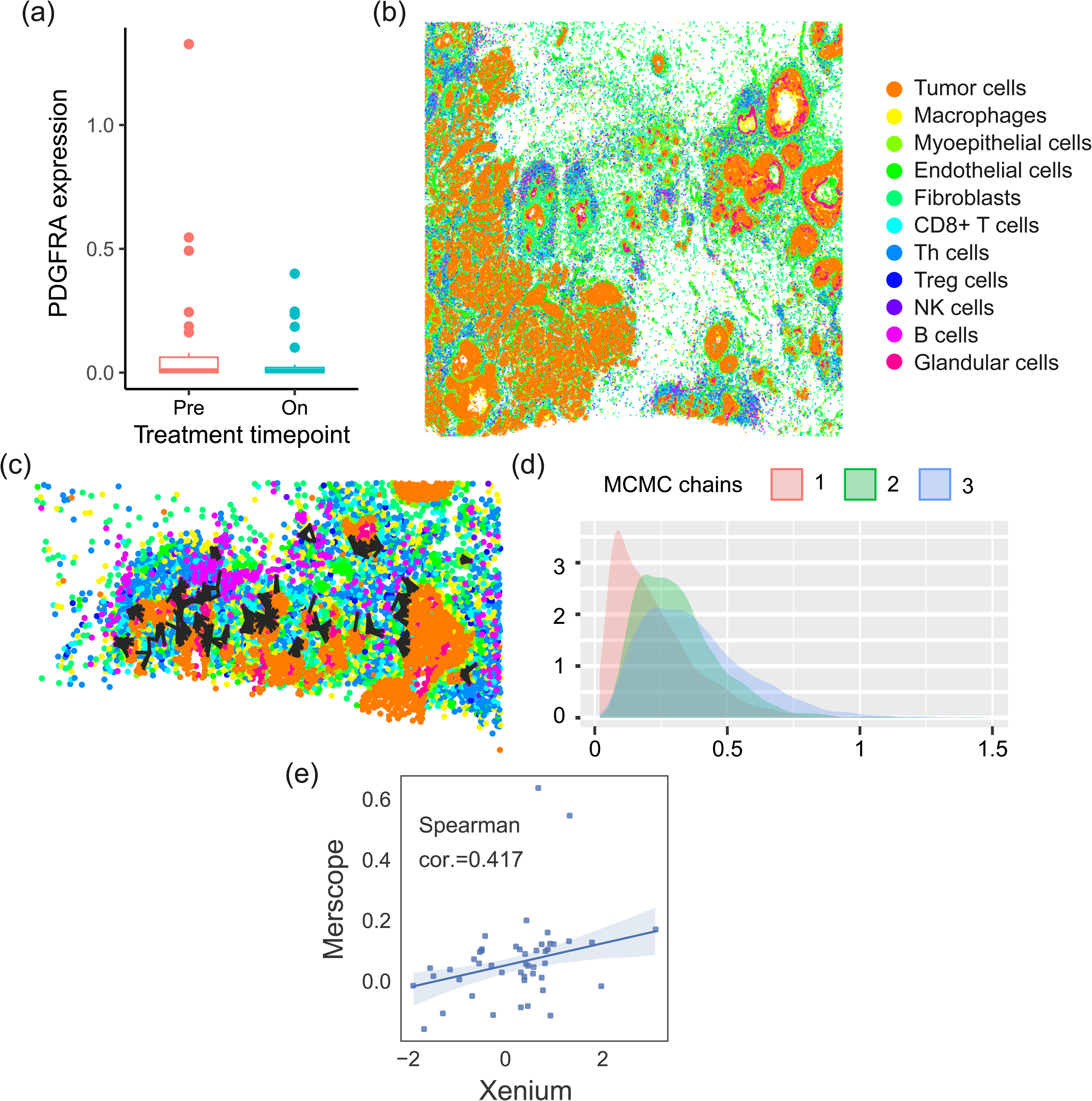
Tumor cell *PDL1* up-regulates *PDGFRA* expression in B cells. (a) The expression of *PDGFRA* in B cells before and after anti-PD1 treatment, in the Bassez cohort. (b) The spatial distribution of the different types of cells in the breast cancer Xenium dataset. (c) The spatial distribution of the CCCs that spacia inferred in this dataset. We zoomed into one area to more clearly show the interactions. (d) The distributions of the inferred βs by spacia, across MCMC iterations, for the interactions between tumor *PDL1* and B cell *PDGFRA*, indicating the direction and the strength of the interaction between these two genes. (e) Scatterplot showing the *β*s from the spacia analyses on both the MERSCOPE and Xenium breast cancer datasets for B cells.

**Sup. Fig. 6.**
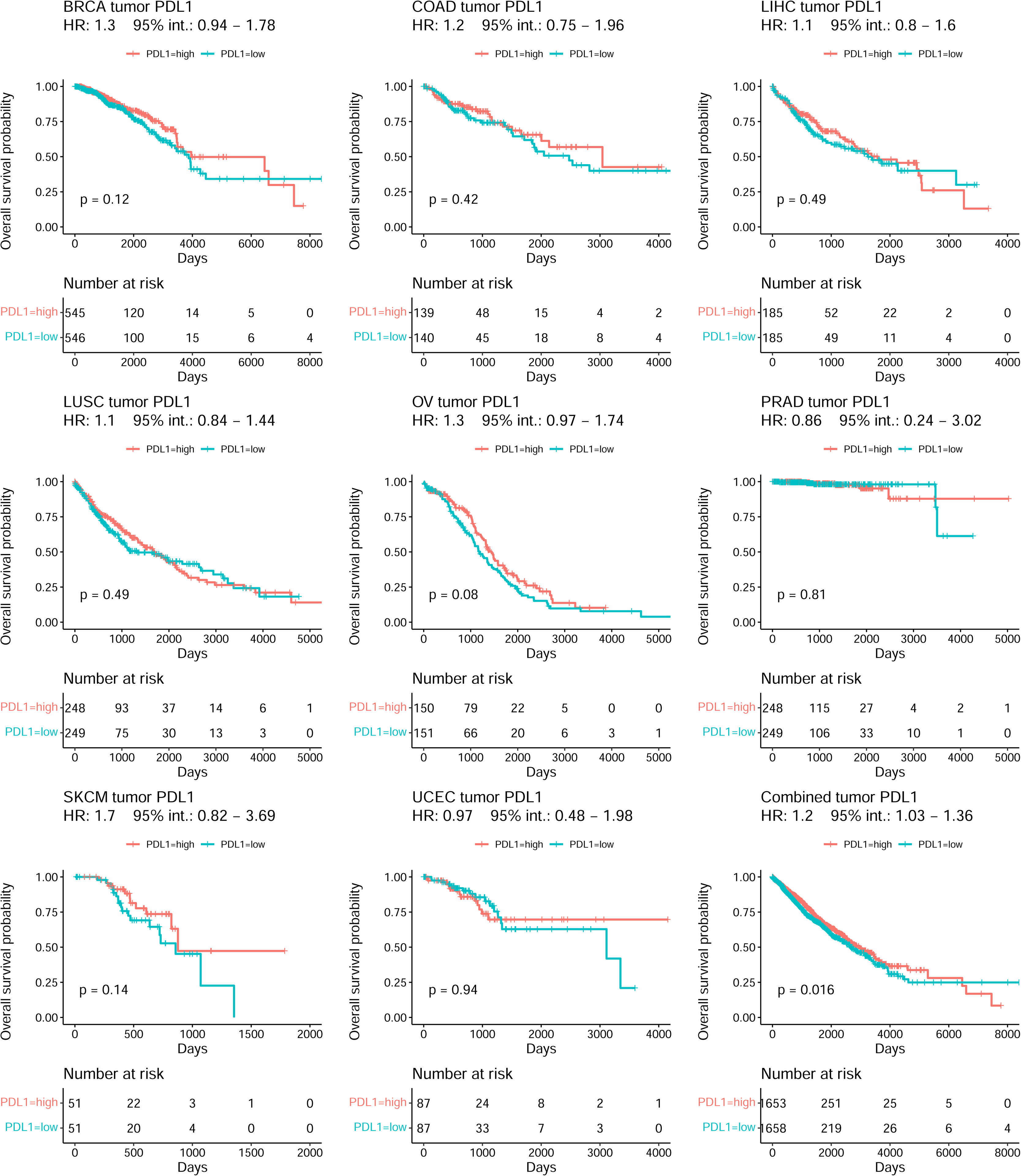
Higher tumor *PDL1* expression is associated with better overall survival in TCGA patients of all eight cancer types, separately or combined. Patients were dichotomized by bulk tumor *PDL1* expression

**Sup. Fig. 7.**
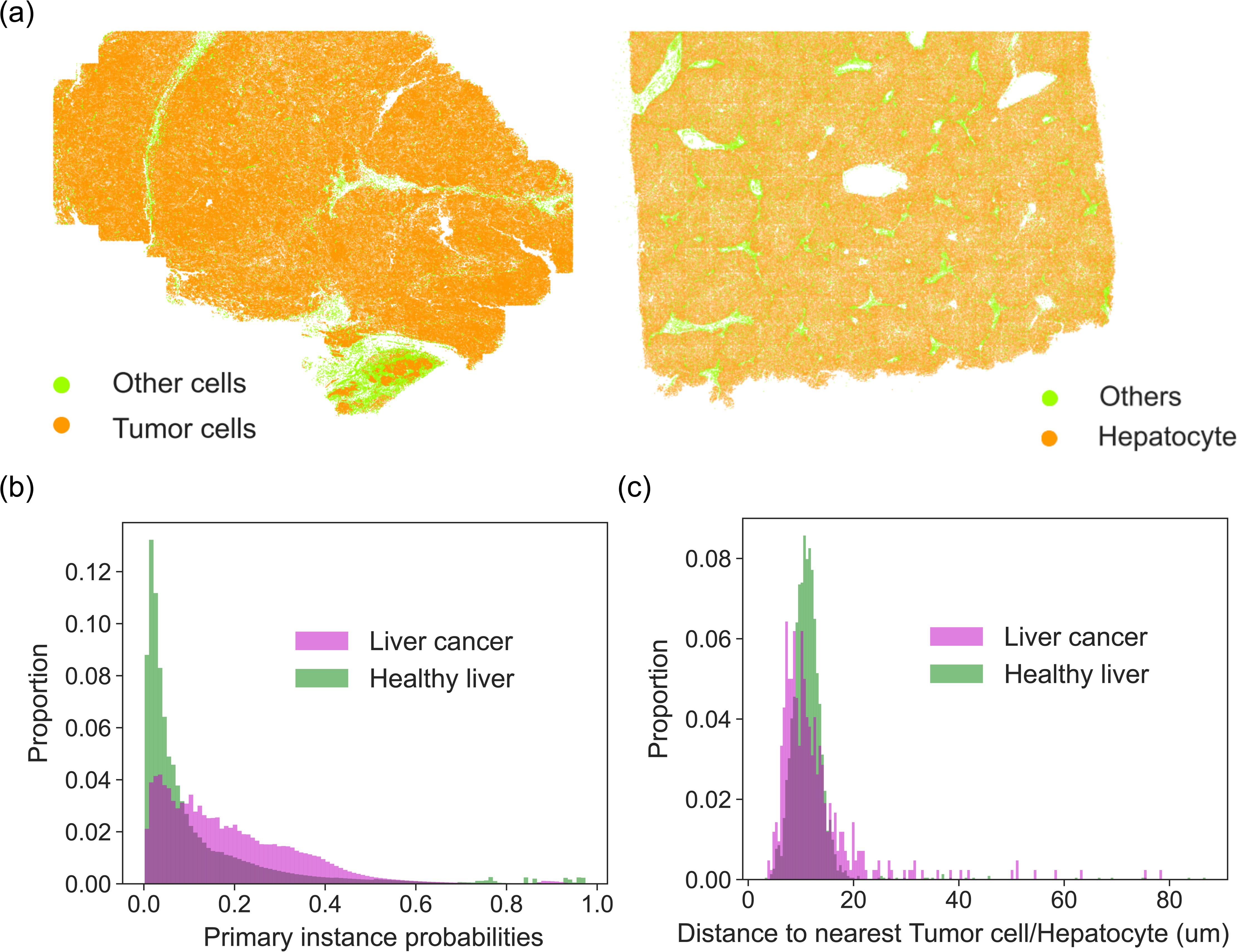
Analyses of the liver cancer and healthy liver CosMx datasets. (a) The spatial distributions of healthy or cancerous hepatocytes *vs.* all other immune/stromal cell types. For clarity, we showed only immune/stromal cell types in Fig. 6a, while we showed hepatocytes in this figure. (b) Density plot showing the primary instance probabilities of the sender cells (γδ T cells) for interacting with the receiver cells (healthy or cancerous hepatocytes). (c) Density plot showing the spatial distances between γδ T cells and their nearest healthy or cancerous hepatocytes.

**Sup. Table 1** Comparing spacia against other related tools previously published.

**Sup. Table 2** GO ontology analyses to identify the top enriched pathways in the sending genes of each sending cell type with the most significant interactions with *JAK1* as receiver gene in the tumor cells.

**Sup. Table 3** Correlations between EMT activation potentials of fibroblasts, endothelial cells and B cells and the EMT levels and lineage plasticity levels of the prostate cancer cells.

**Sup. Table 4** CD8-PDL1 signature genes in all eight cancer types.

**Sup. File 1** Mathematical and implementation details of spacia.

**Sup. File 2** Additional bioinformatics analyses associated with this study.

