## Supplemental File 1 for "Mapping Cellular Interactions from Spatially Resolved Transcriptomics Data"

### Sup. File 1: Spatially Resolved Transcriptomics-based Cell-to-Cell Interaction Detection Using A Multiple Instance Learning Model

August 25, 2023

#### 1 Overview

Spacia investigates the interactions between the signal-receiving cells and signal-sending cells from a spatially resolved transcriptomics dataset (such as Seq-scope, Slide-seq, HDST, Visium, SEPFISH, MERSCOPE, etc). Spacia models the expression of the one receiving cell (as the response variable) as the function of the expression signature of multiple sending cells in the vicinity of this receiving cell. This task motivates a multi-instance learning (MIL) approach, which is often useful in describing multiple-to-one relationships. In MIL terminology, the input dataset consists of "bags" and "instances". The "bags" are individual receiving cells that are labeled by the expression of the receptor genes/pathways (we model one gene or pathway (see below) at a time), and each bag is a collection of "instances", which are the sending cells in the vicinity characterized by the instance-level features. The features include the expression of the ligands and their up-stream regulators, and also the distances of the sending cells to the receiving cells.

#### 2 Compatibility

Spacia is capable of analyzing spatially resolved transcriptomics data for cell-to-cell interactions in the following cases:

a Lower resolution data (e.g. Visium)
Spacia will work at the sequencing spot level. The users can either define the cell type of the spots according to their gene expression or their pathological imaging appearances. It is understood that a spot will have multiple cells, and spacia will not be able to investigate single cell-to-cell interaction, but rather one group of cells of one or even multiple cell types against another group of cells. We suggest removing spots with mixed expression patterns and then removing genes not belonging to the cell type of the spot (such as immune cell-enriched genes in tumor cell spots) from the remaining spots. The algorithm will assume it has received clean data.

b High-resolution data without matched images. (e.g. Slide-seq)
Spacia will not work. There are no matching images to define cell masks.

c High-resolution data with matched images. (e.g. MERSCOPE, HDST, and SeqScope)
Spacia is mainly designed for such types of technologies/data. The users will define cell masks according to the imaging data via segmentation techniques. Then the expression of the sequencing spots belonging to the same cell should usually be aggregated by sum. Cell type can be determined from the aggregated expression or from the imaging appearances. Unlike (a), spacia will be able to investigate single cell-to-single cell interactions in this case.

#### 3 Input data preparation

When using spacia, users are responsible for cell segmentation, cell type annotation, and expression normalization as this software does not process image data. Before using spacia, users need to prepare their input data, which includes normalized counts of the spatial transcriptomics experiment, the positional information of the cells, the types of cells, and optionally, the list of sending/receiving genes. We advise log-transforming the normalized gene expression data and we also recommend removing noises from the spatially resolved transcriptomic data using tools such as Sprod[1].

One challenge of spatially resolved transcriptomics data is the sparsity of the data, and our solution is to aggregate single genes (receiving or sending) into pathways, composed of highly correlated genes, to boost the signal. This operation could also help reduce the dimensionality of the input data and reduce the computational burden. We have several options:

a No aggregation
Users have the option to provide either one or multiple single genes for the sending/receiving pathways. Expressions of all the genes in the provided list will be treated individually in the downstream analyses.

b Knowledge-driven aggregation
Users supply a file with information about which genes to group into pathways for the sending/receiving pathways. This data can be compiled from resources such as CellChat and CellphoneDB. It's important to include only genes that are expected to have a positive correlation in the same pathway, for the same reasons as above.

c Correlation-driven aggregation
In this mode, users will provide a list of one or more genes. Spacia will then extend each of these genes based on highly correlated genes. There are two ways the genes will be aggregated: 'simple' and 'weighted'. In the 'simple' method, the expressions of the top positively correlated genes are averaged with equal weights. With 'weighted', the expressions of the top correlated genes by absolute correlation are summed into the seed gene expression using weighted averages. Here, weights are the Pearson correlation coefficients of the genes. Both positive and negative correlation values will be used as weights. If not specified, the 'simple' method is used by default.

d Clustering-driven aggregation
This mode is run when no genes or pathway information is given. Spacia applies hierarchical clustering, a type of unsupervised machine learning algorithm, to construct gene modules based on positive correlation. The identified clusters of genes become the aggregated pathways.

e PCA-driven aggregation
Spacia will run Principal Component Analysis (PCA) on the sending gene expression matrix and will keep only the top Principal Components (PCs) as the "sending pathways". The number of top PCs to keep is the choice of the users. This approach significantly reduces the number of genes/pathways on the sending side from hundreds to several dozen or even fewer PCs. After Markov Chain Monte Carlo (MCMC) has estimated the PC-level betas, the calculated results are mapped back into the original space so that the results can be interpreted at the gene level. Our investigations revealed that this mode boosts the signal significantly, and all our analyses were done in this aggregation mode.

Regardless of the chosen mode, spacia requires a quantile cutoff value to transform the receiving gene's or pathway's expression value into a binary response variable. By default, this value is set to the median, but this cutoff can either be user-defined, or spacia can automatically determine an appropriate cutoff. In the automatic mode, if the data follows a bimodal distribution, the cutoff will be set at the midpoint between the two means of the two fitted Gaussian distributions. If the two fitted Gaussian distributions' 1 standard deviations overlap, i.e. not a bimodal distribution, the mean + 1 standard deviation of the distribution with a smaller mean will be used as a cutoff instead. The cutoff value is applied to the aggregated expression of the pathways if correlation aggregation is enabled.

### 4 Model specification

#### 4.1 Notation

Let $\mathrm{B}_i$ denote bag $i$ (receiving cell) containing $m_i$ instances (sending cells in the neighborhood), and $y_i$ denote the observed binary bag label(activation of a receiving gene or pathway) for $i = 1, ..., n$, where $n$

is the total number of bags observed. For the expression data part, suppose there are $e$ features(genes or pathways) that characterize each instance $j$. We use $X_i^e = (x_{ij}^e)_{j=1}^{m_i}$ to denote the $m_i \times (e+1)$ design matrix of $\mathrm{B}_i$, where $x_{ij}^e = (1, x_{ij1}, ..., x_{ije})$ is a row vector of length $e+1$. For the cell-to-cell distance data part, we use $X_i^d = (x_{ij}^d)_{j=1}^{m_i}$ to denote the $m_i \times 2$ design matrix of $\mathrm{B}_i$, where $x_{ij}^d = (1, d_{ij})$ is a row vector of length 2. $d_{ij}$ is the distance between two cells and the column of 1s is the intercept term.

In our application, it's important to note that not every instance, or "sending cell", is necessarily significant. The spatial range within which two cells interact is likely to differ, depending on the specific pathways and cell types interacting. We implement a relatively flexible spatial cutoff when forming the 'bags,' which denote the pairs of sending-receiving cells. We include sending cells within this distance limit of a receiving cell into the bags, due to the uncertainty surrounding the precise range of interaction. This is especially the case given that the interacting distance between cells can differ amongst different interaction pairs. In our algorithm, the model will learn what this range should be. In each round of estimation, we term sending cells that are truly influencing the receiving cells as the primary instances, and vice versa. Notation-wise, for each bag $i$ (receiving cell), we refer to those primary instances collectively as $\mathrm{B}_i^*$. Let $\delta_{ij}$ be a latent indicator variable, with $\delta_{ij} = 1$ indicating that instance $j$ is a primary instance of $\mathrm{B}_i$ and 0 otherwise.

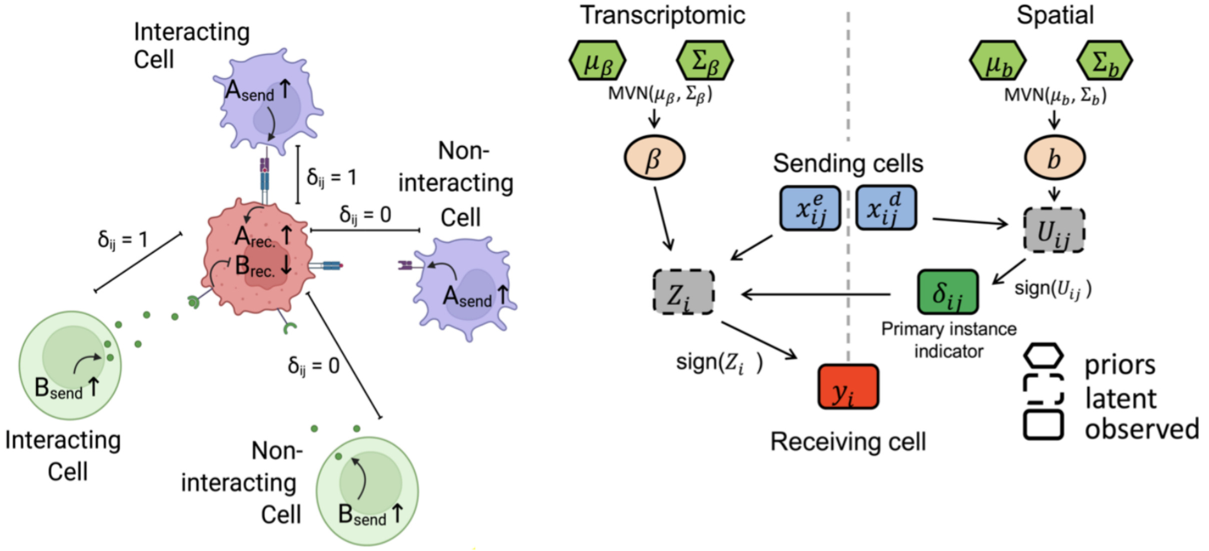


Figure 1: Structure of the Spacia Model

#### 4.2 Prior specification

We model the relationship between the covariate matrix $X_i^e$ (expression of genes/pathways in the sending cells) and the binary outcome $y_i$ (high or low expression of one target gene/pathway in the receiving cells) by a probit regression, assuming the primary instances of all bags are known.

$$y_i = \mathrm{sign}(Z_i),$$
$$Z_i = \sum_{j=1}^{m_i} \delta_{ij} (x_{ij}^{\mathrm{e}})^T \beta / C_i + \epsilon_i,$$

Here, $\beta = (\beta_r)_{r=0}^e$, where $\epsilon_i \overset{ind}{\sim} N(0,1)$ for $i = 1, ..., n$, is a vector of the regression coefficients and also the intercept term that explains the effect of the gene expression of the sending cells on the outcome variable. The normalizing factor, $C_i$, is used to consider the varying number of primary instances in each bag. In our application, $C_i = 1$, which corresponds to the sum contribution of all sending cells. In case where $\mathrm{B}_i^* = \emptyset$, we let $Z_i = \beta_0 + \epsilon_i$. Namely, $\Pr(y_i = 1|X_i^e, \beta, \delta_{i1}, ..., \delta_{im_i}) = \Phi\left(\sum_{j=1}^{m_i} \delta_{ij}(x_{ij}^{\mathrm{e}})^T \beta\right)$, where $\Phi(\cdot)$ is the cumulative distribution function of the standard normal distribution.

Similarly, another probit regression is used to model the latent primary indicator of instance $j$ in bag $i$ (i.e., $\delta_{ij}$).

$$\begin{aligned} \delta_{ij} =& \mathrm{sign}(U_{ij}), \\ U_{ij} =& (x_{ij}^{\mathrm{d}})^T b + e_{ij}, \end{aligned}$$

where $e_{ij} \overset{ind}{\sim} N(0,1)$ for $i = 1, ..., n$ and $j = 1, ..., m_i$. The column vector of intercept and coefficients, $b = (b_r)_{r=0}^{1}$, describes the relationship between the instance status (primary or non-primary) and the distances between sending cells and receiving cells. We have $\Pr(\delta_{ij} = 1|x_{ij}^d, b) = \Phi\left((x_{ij}^{\mathrm{d}})^T b\right)$, similar to the probit regression of observed outcomes. $Z_i$'s and $U_{ij}$'s are the latent variables in each probit model, and we would only observe these variables when all primary instances are known. The addition of these latent variables in a model is also known as data augmentation, and it allows posterior sampling by MCMC method. We fix the variance of $\epsilon_i$ and $e_{ij}$ at 1 for model identifiability. We propose a Bayesian hierarchical model and employ conjugate priors, an approach that is frequently used in Bayesian literature for convenient posterior sampling. We specify the priors for the regression coefficients, $\beta$, as $\beta|\mu_\beta, \Sigma_\beta \sim N_{e+1}(\mu_\beta, \Sigma_\beta)$. Following the standard practice, we set $\mu_\beta = (0, 0, ..., 0)^{\mathrm{T}}$. Also, we set $b|\mu_b, \Sigma_b \sim N_2(\mu_b, \Sigma_b)$ and assign $\mu_b = (0, 0)^{\mathrm{T}}$. For $\Sigma_\beta$ and $\Sigma_b$, we employ a diagonal matrix with $(1, 1, ..., 1)$ on the diagonal entries.

#### 4.3 Posterior Computation

We define $X = \{(X_i^e)_{i=1}^n, (X_i^d)_{i=1}^n\}$ as the set of covariate matrices from all bags, $y = (y_i)_{i=1}^n$ as a column vector of length $n$. Suppose $\Delta$ and $U$ are $(\Sigma_{i=1}^n m_i) \times 1$ column vectors of binary indicators $\delta_{ij}$ and corresponding latent variables $U_{ij}$ for $i = 1, \ldots, n$ and $j = 1, \ldots, m_i$, respectively. Similarly, $Z = (Z_i)_{i=1}^n$ represents a column vector of latent variables $Z_i$ associated with the bag labels $y_i$ for $i = 1, \ldots, n$. All model parameters and latent variables are represented by $\Theta = (\beta, b, \Delta, Z, U)$. With the hyper-parameters $\mu_\beta$, $\Sigma_\beta$, $\mu_b$, and $\Sigma_b$ defined, the complete probability model is represented as

$$\begin{aligned} p(y, \Theta|X) =& p(y|Z) \times p(Z|X, \Delta, \beta) \times p(\Delta|U) \times p(U|X, b) \\ & \times p(\beta|\mu_\beta, \Sigma_\beta) \times p(b|\mu_b, \Sigma_b) \\ =& \prod_{i=1}^{n} \left\{ p(y_i \ Z_i) \cdot p(Z_i|x_\delta, \beta) \cdot \left[ \prod_{j=1}^{m_i} p(\delta_{ij}|U_{ij}) \cdot p(U_{ij}|x_{ij}, b) \right] \right\} \\ & \times p(\beta|\mu_\beta, \Sigma_\beta) \times p(b|\mu_b, \Sigma_b) \end{aligned}$$

We employ MCMC to extract random samples from the joint posterior distribution $p(\Theta|X, y)$, which is proportional to $p(y, \Theta|X)$. The conditional posterior distribution of each parameter (or latent variable), when given all the other parameters, becomes a tractable, known family of distributions, so that the samplings are all amenable to Gibbs samplers.

#### 4.4 Posterior Inference

Let $T$ be the number of iterations after the burn-in period for the Gibbs sampler. The point estimation of the variables can be made based on posterior means, such as $\hat{\beta} = \frac{1}{T} \sum_{t=1}^{T} \beta^{(t)}$. Uncertainty estimation is quantified using equal-tailed intervals.

We calculate the posterior inclusion probability to identify the primary instances in bag $i$:

$$\widehat{\pi}_{ij} = \frac{1}{T} \sum_{t=1}^{T} \delta_{ij}^{(t)}, \ j = 1, ..., m_i$$

We define the instance $j$ in bag $i$ as primary if $\widehat{\pi}_{ij} > \theta$. $\theta$ is a cutoff established by controlling the Bayesian false discovery rate (FDR) [2]on all instances of the entire dataset. We estimate the FDR for a given $\theta$,

$$\widehat{\mathrm{FDR}}(\theta) = \frac{\sum_{i=1}^{n}\sum_{j=1}^{m_i}(1-\widehat{\pi}_{ij})\cdot \mathrm{I}(\widehat{\pi}_{ij} > \theta)}{\sum_{i=1}^{n}\sum_{j=1}^{m_i}\mathrm{I}(\widehat{\pi}_{ij} > \theta)},$$

where the indicator function, $\mathrm{I}(\cdot)$, is equal to 1 if the function is satisfied, and 0 otherwise. We also define the FDR to be 0 if the denominator is zero. Let $\kappa \in (0,1)$ be an FDR we aim to control, then we choose the $\theta$ so that $\widehat{\mathrm{FDR}}(\theta) \leq \kappa$.

### 5 Outputs

The software outputs several estimated variables that describe the association between the signal receiving and sending cells. Our algorithm estimates $\widehat{\pi}_{ij}$, which are the probabilities of the sender cells being truly responsible for the receiving cells' phenotypes (primary instances). There are also two coefficients calculated from the nested probit regression model, $\hat{b}$ and $\hat{\beta}$. $\hat{b}$ explains how the distances between sender and receiver cells affect the probabilities of the sender cells being primary, and $\hat{\beta}$ shows the effect of the expression of the genes or pathways in the primary sender cells on the receiving cells. Finally, the cutoffs to define primary instances, $\hat{\theta}$, are calculated from $\kappa$s, the user-specified FDRs.

Spacia investigates the converged distributions of the $\beta$ and $b$ variables to perform statistical inference. For both $\beta$ and $b$, spacia samples their values from 50 uniformly spaced MCMC iterations after burn-in. This is done so that we have the same sample size (in terms of sampled values) regardless of how many MCMC iterations have been performed in each spacia run. Then we use the T-test to test whether the $\beta$ variables are significantly different from 0 and the $b$ variables are significantly smaller than 0.
