## Supplemental File 2 for "Mapping Cellular Interactions from Spatially Resolved Transcriptomics Data"

**Sup. File 2 Additional Results Related to spacia**

**Numbers of senders in each bag in the simulated datasets**

The histogram below shows the numbers of all possible senders (all instances in the bags of receiver cells), and the number of primary instances (truly interacting sender cells), in our simulation analysis. The numbers of primary and all senders were simulated to fit our best guess about the possible range of these two numbers in real data, according to our own experiences in exploring real SRT data.

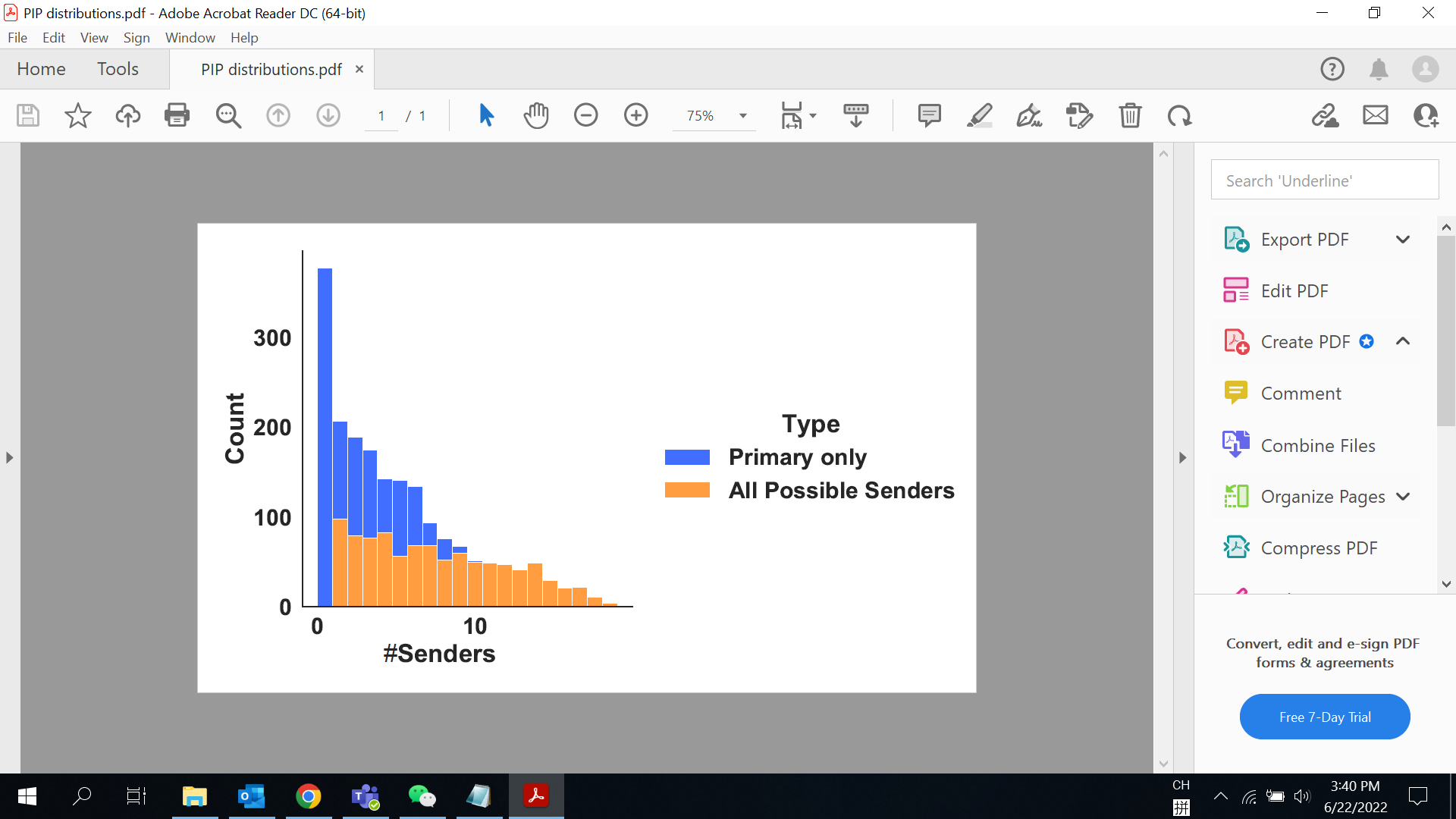

***Sup. File 2 Fig. 1*** *The numbers of all possible senders and the number of primary instances in our simulation analysis.*

**Robustness of spacia with respect to choices of hyper-paramaters**

The only hyper-parameter in the spacia model is the variances we specified for the prior distributions of *b* and *beta*. This hyper-prior structure was employed to stabilize model estimates. Our default choice of this parameter is *1* for both *b* and *beta*. We varied this parameter from *0.1* to *5*, and assessed its impact on the simulation dataset in **Fig. 2**. We employed the AUROC for identifying the true senders as the criterion, as we found that all the performance criteria employed in **Fig. 2** are generally highly correlated with each other and AUROC has a straightforward interpretation. As can be seen from the figure below, when the hyper-prior is chosen to be between *0.1* and *2*, spacia has very good performance, with the choice of *1* leading to the best performance. When the prior is larger than *5*, the model failed to operate properly and did not converge. Therefore, we specified *b*’s and *beta*’s prior distributions to have a variance of *1* in the spacia model.

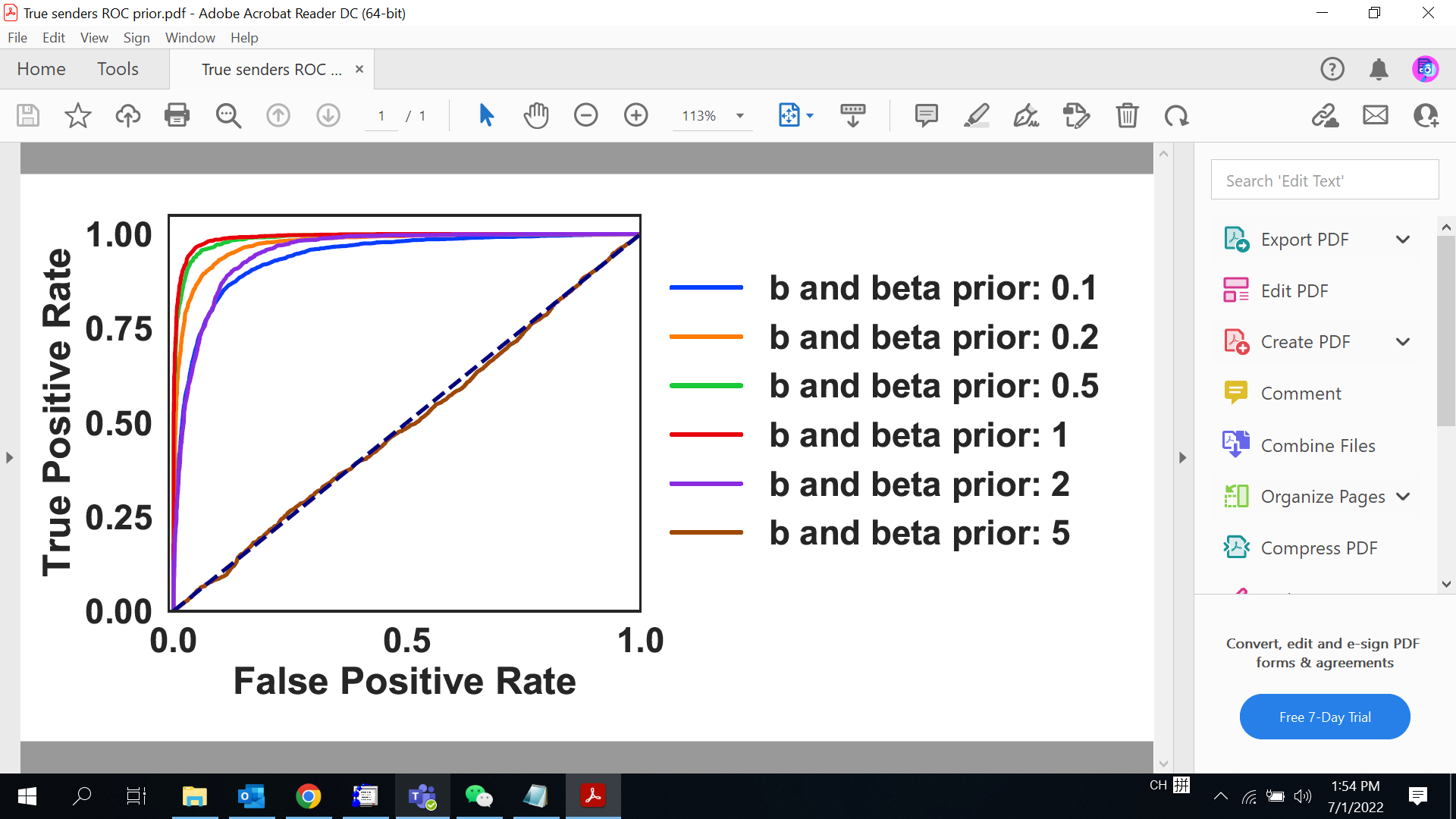

***Sup. File 2 Fig. 2*** *AUROC for identifying true senders in the simulated dataset, for different choices of the hyper-parmaeter.*

**Cell typing for the MERSCOPE datasets**

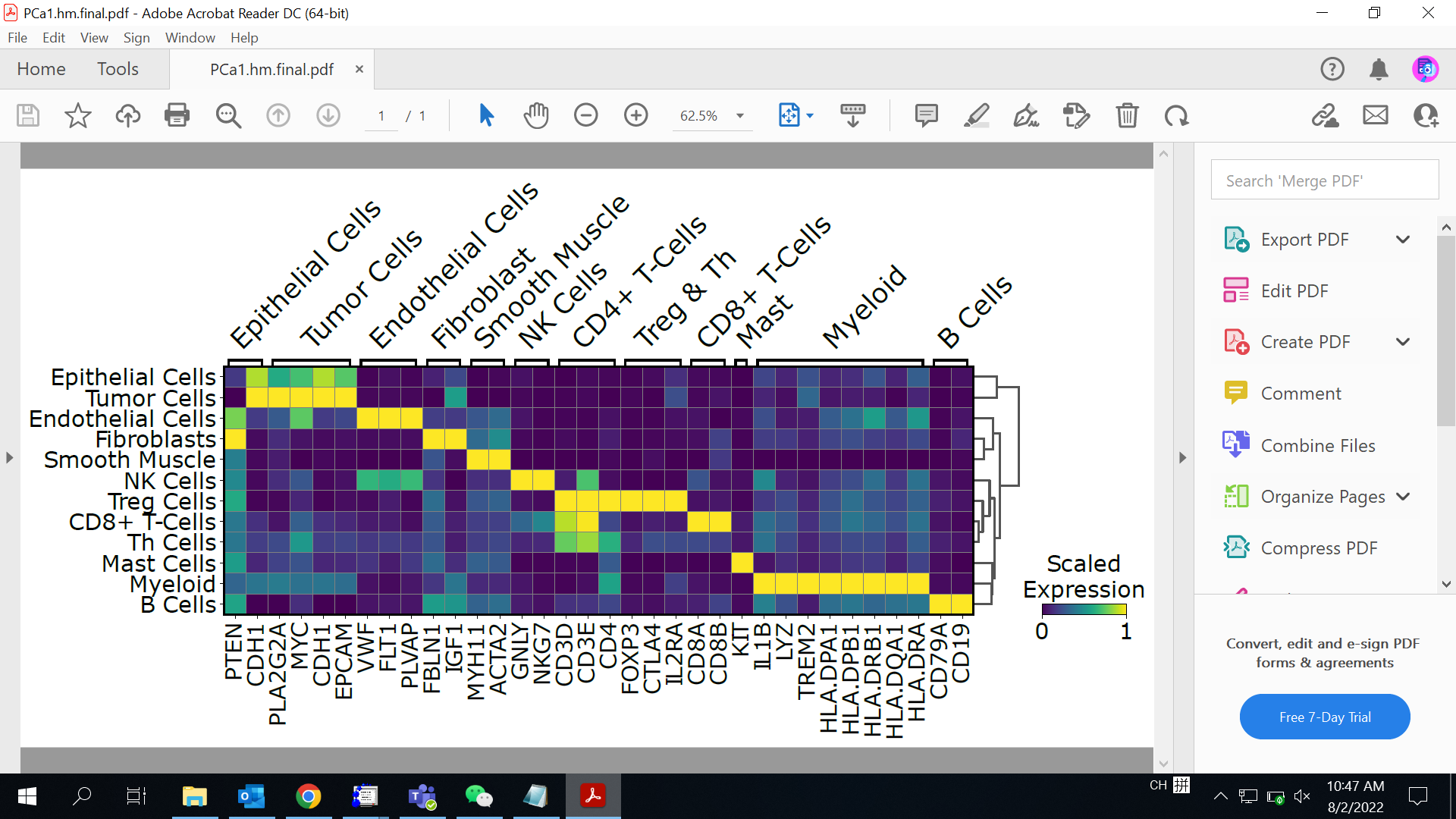

***Sup. File 2 Fig. 3*** *Heatmap showing the expression of the key marker genes for all cell types detected in the Prostate Cancer dataset*

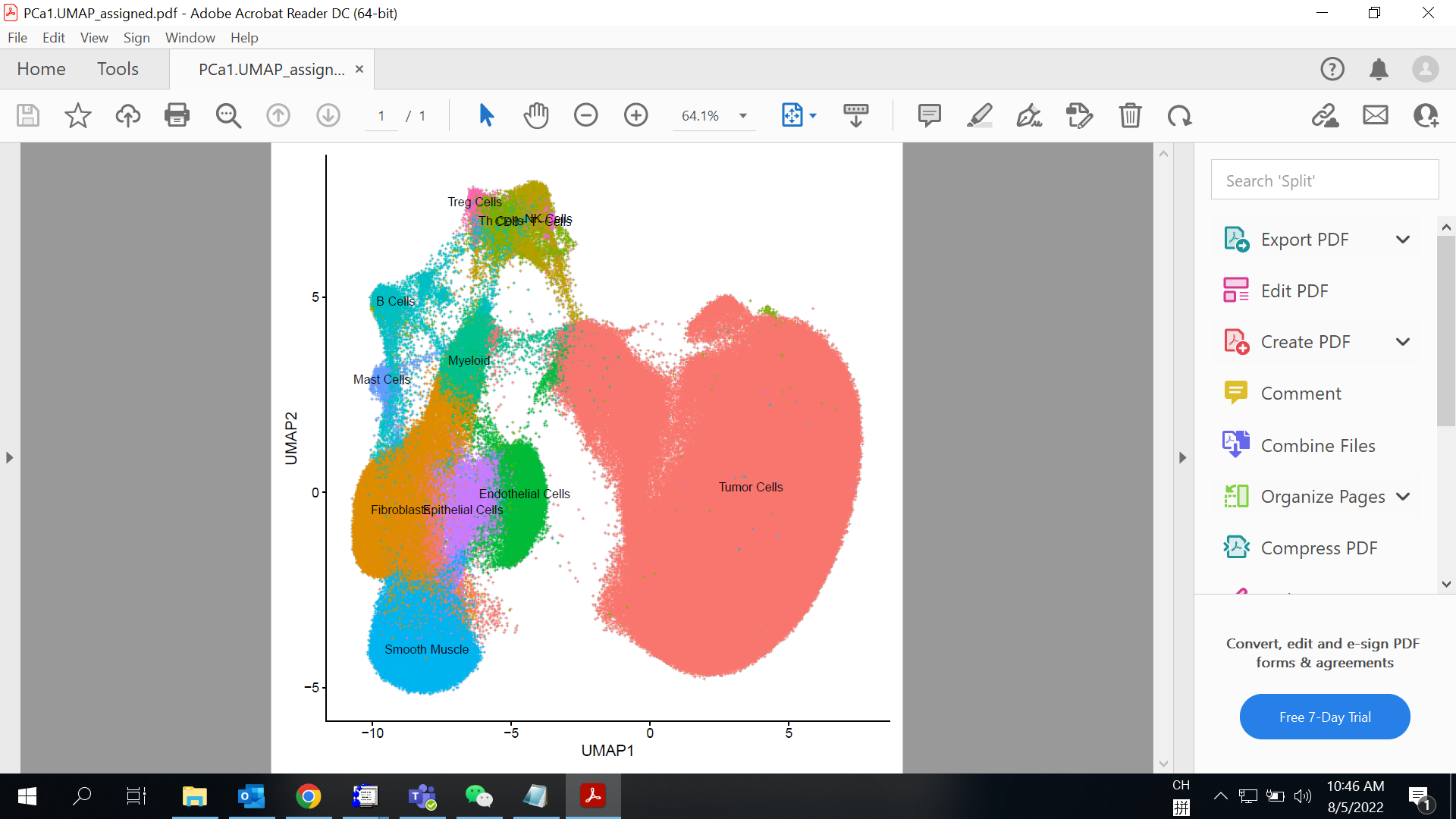

***Sup. File 2 Fig. 4*** *Umap plot showing the cells and their cell types of the Prostate Cancer dataset.*

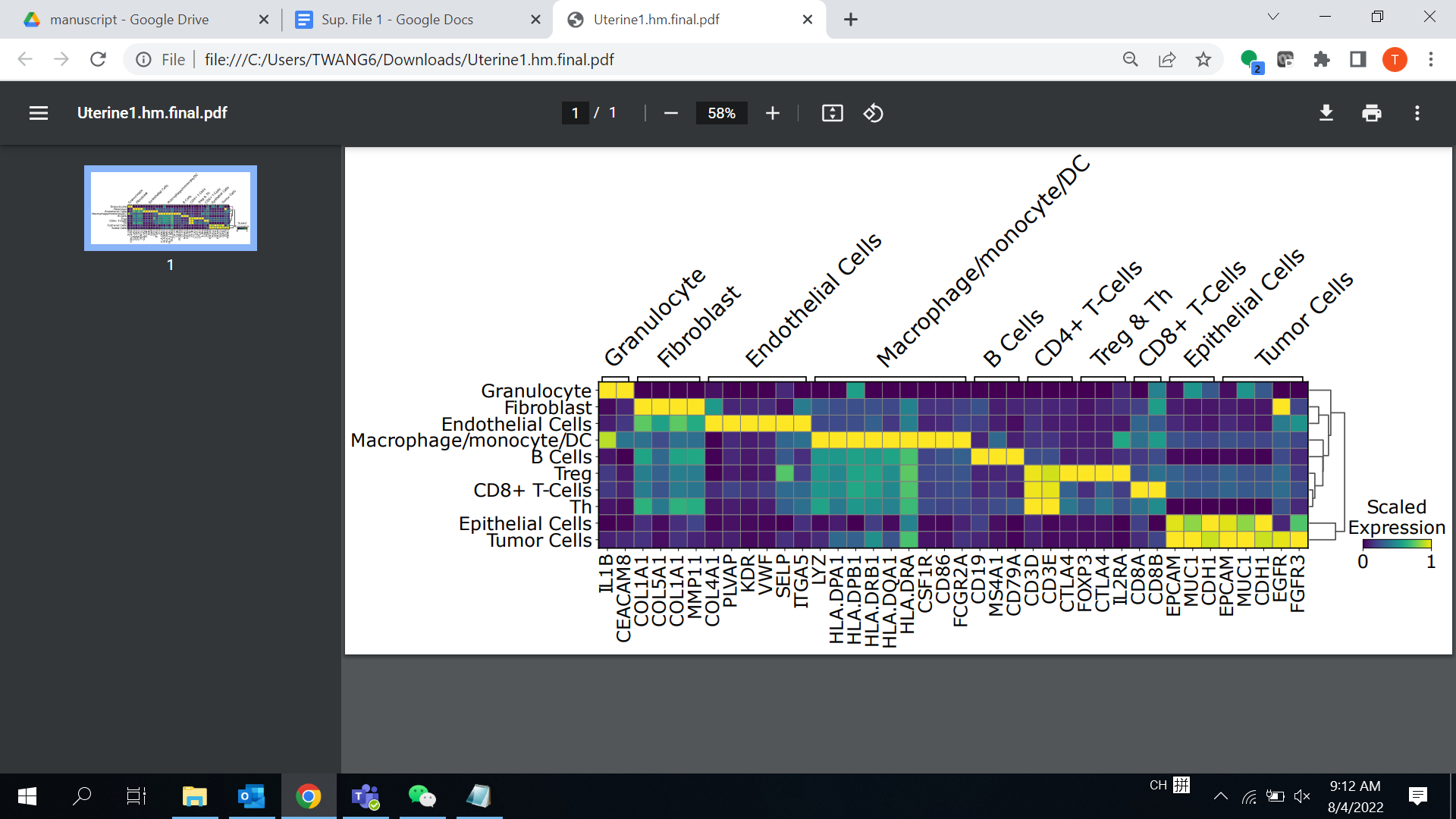

***Sup. File 2 Fig. 5*** *Heatmap showing the expression of the key marker genes for all cell types detected in the Uterine Cancer dataset*

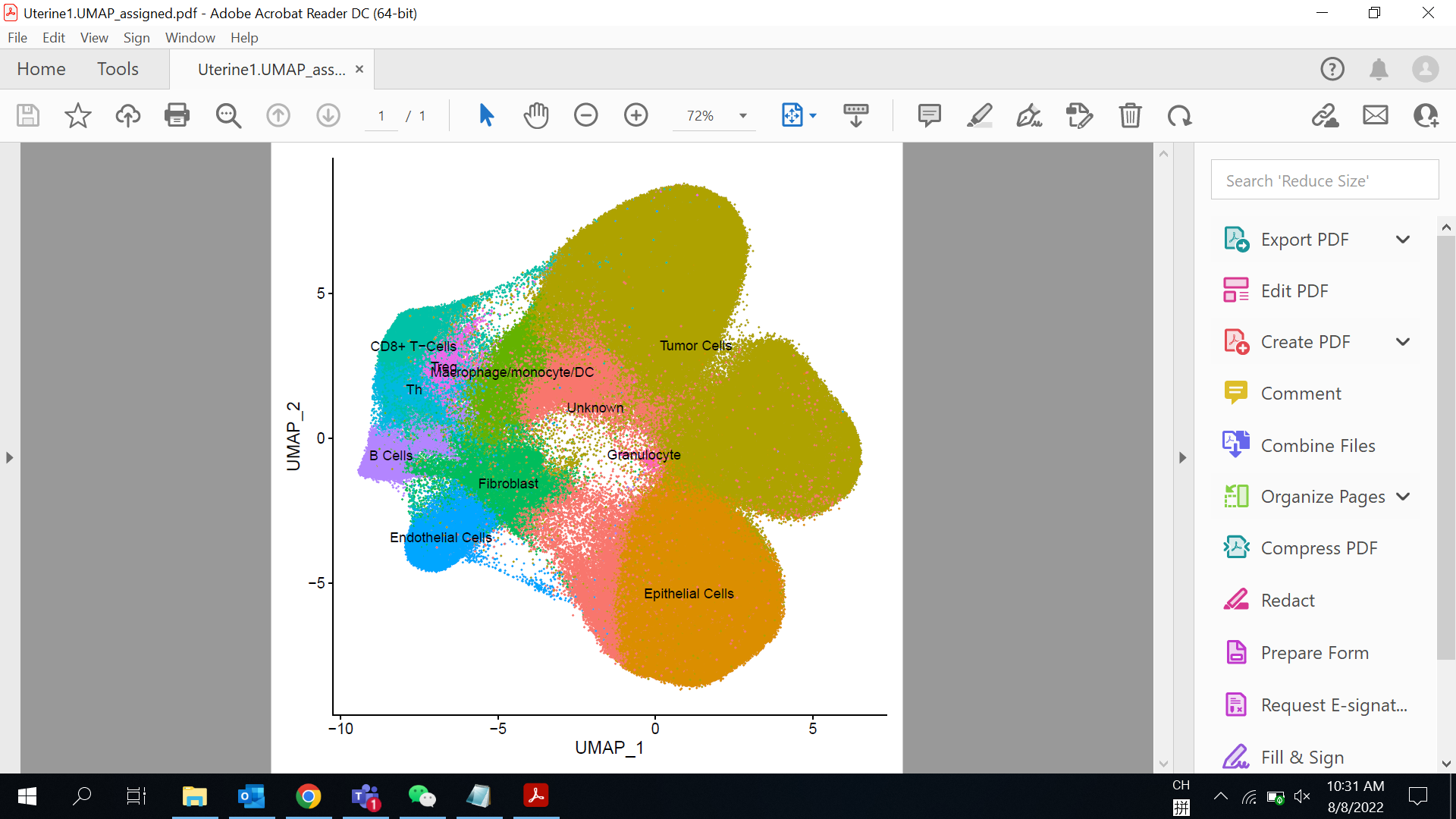

***Sup. File 2 Fig. 6*** *Umap plot showing the cells and their cell types of the Uterine Cancer dataset.*

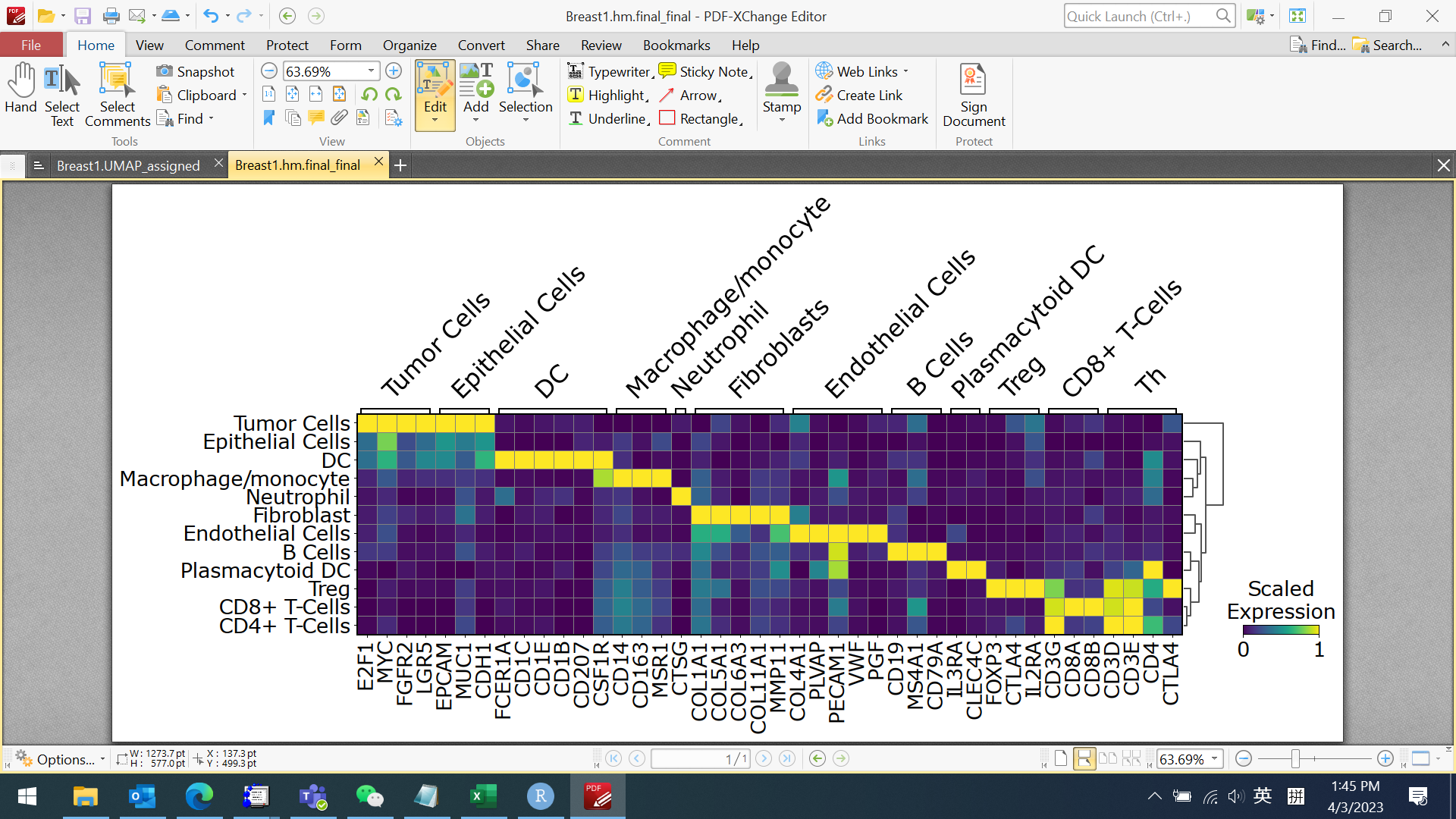

***Sup. File 2 Fig. 7*** *Heatmap showing the expression of the key marker genes for all cell types detected in the Breast Cancer dataset*

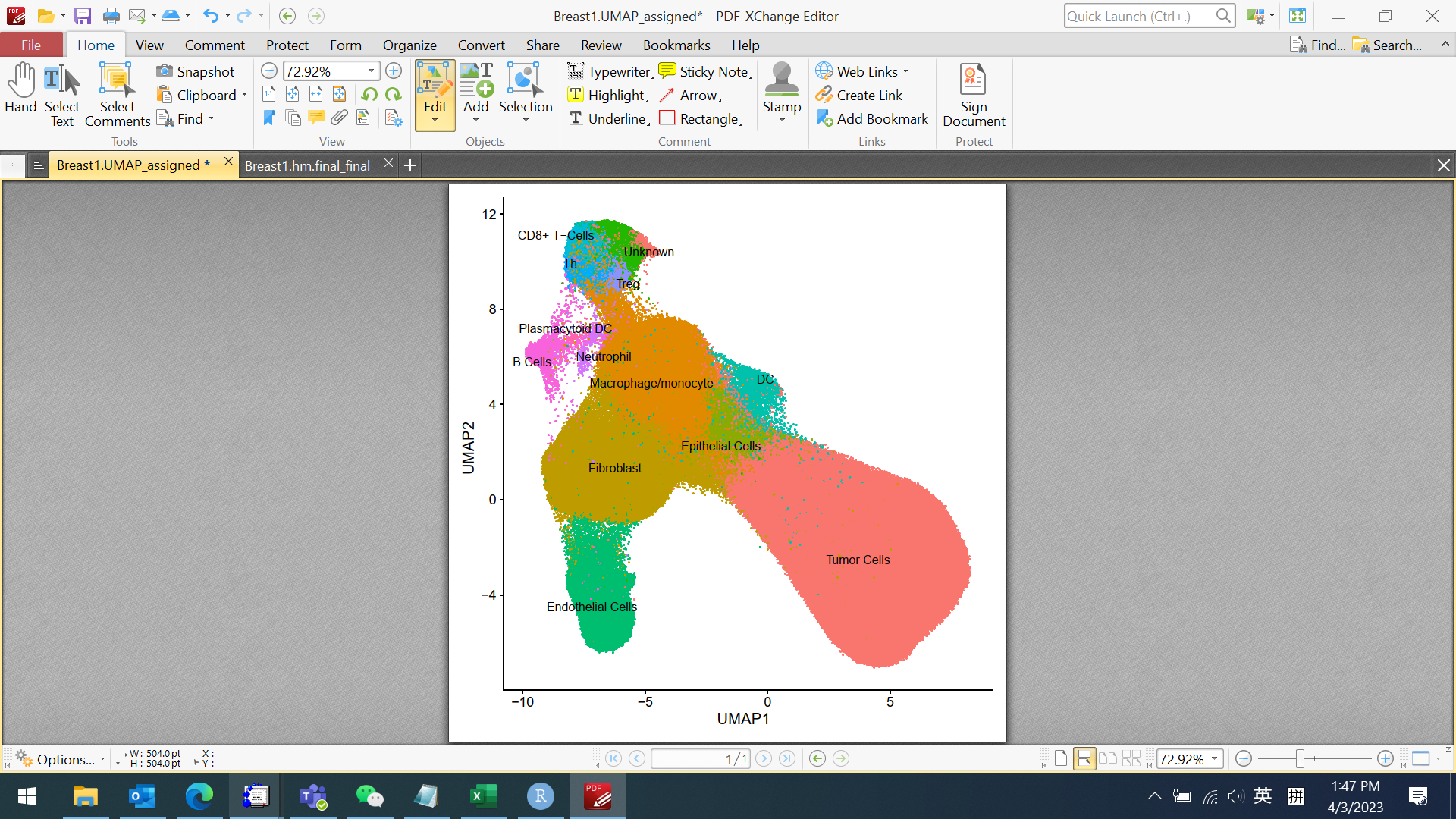

***Sup. File 2 Fig. 8*** *Umap plot showing the cells and their cell types of the Breast Cancer dataset.*

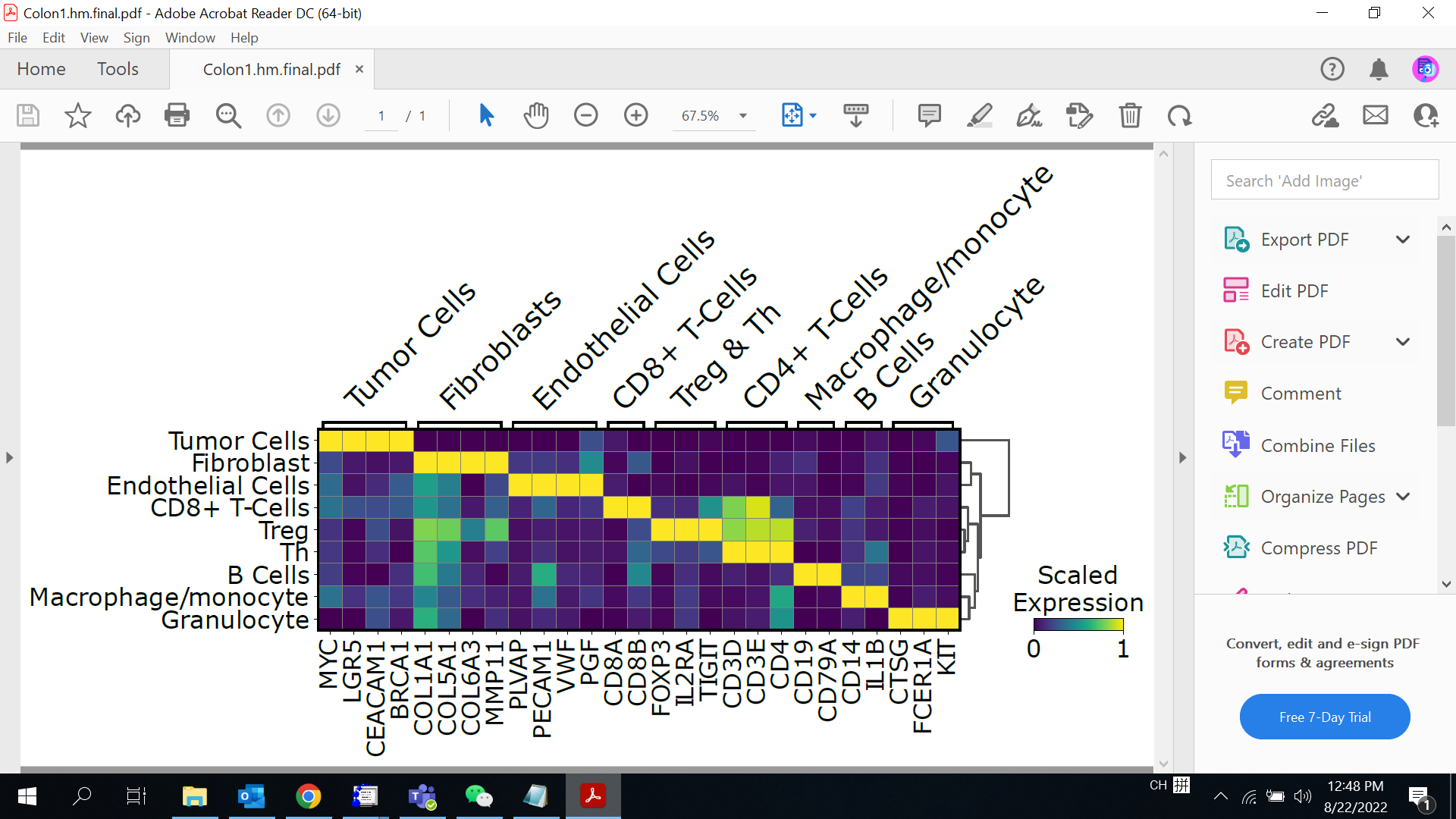

***Sup. File 2 Fig. 9*** *Heatmap showing the expression of the key marker genes for all cell types detected in the Colon Cancer dataset*

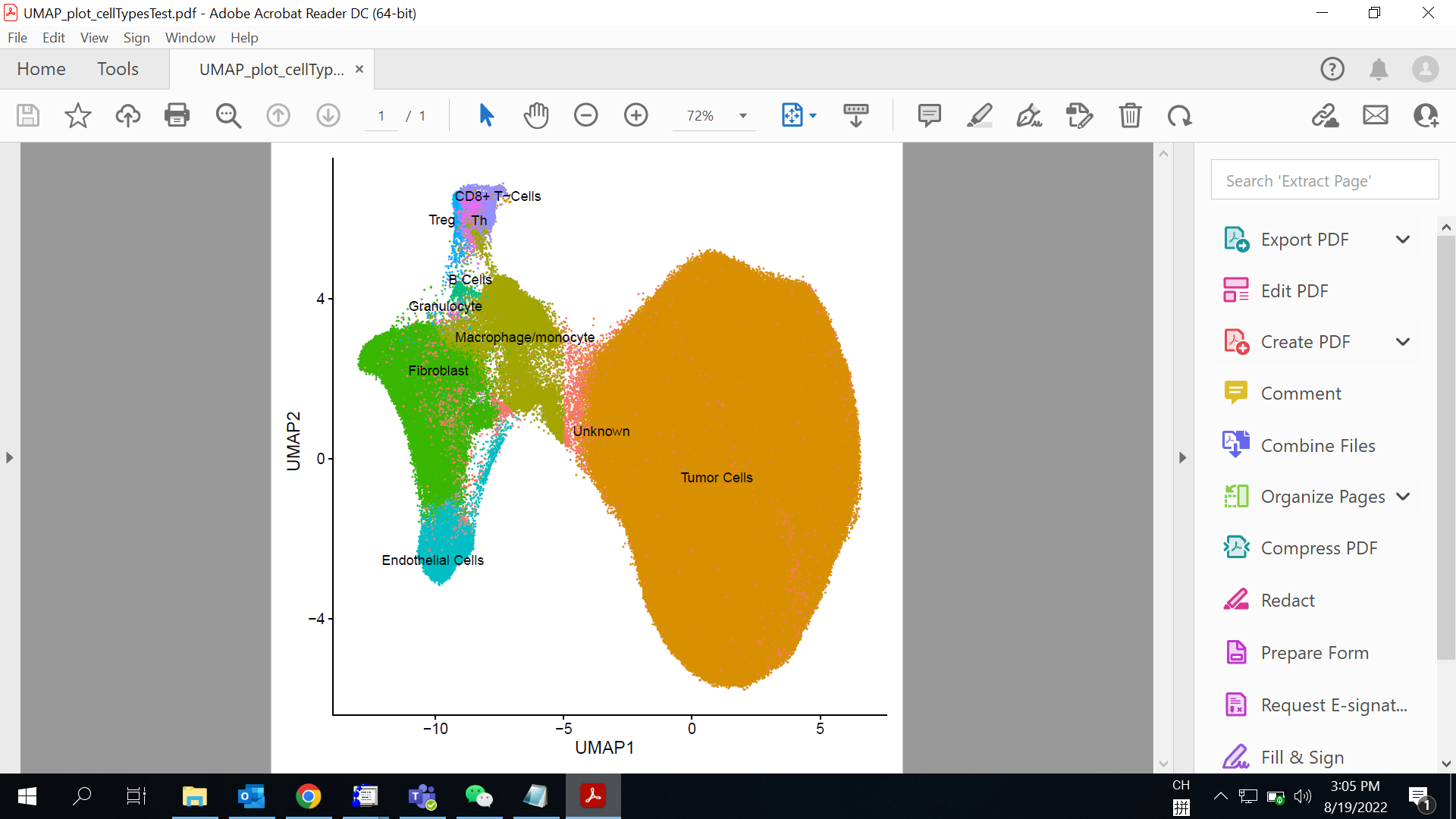

***Sup. File 2 Fig. 10*** *Umap plot showing the cells and their cell types of the Colon Cancer dataset.*

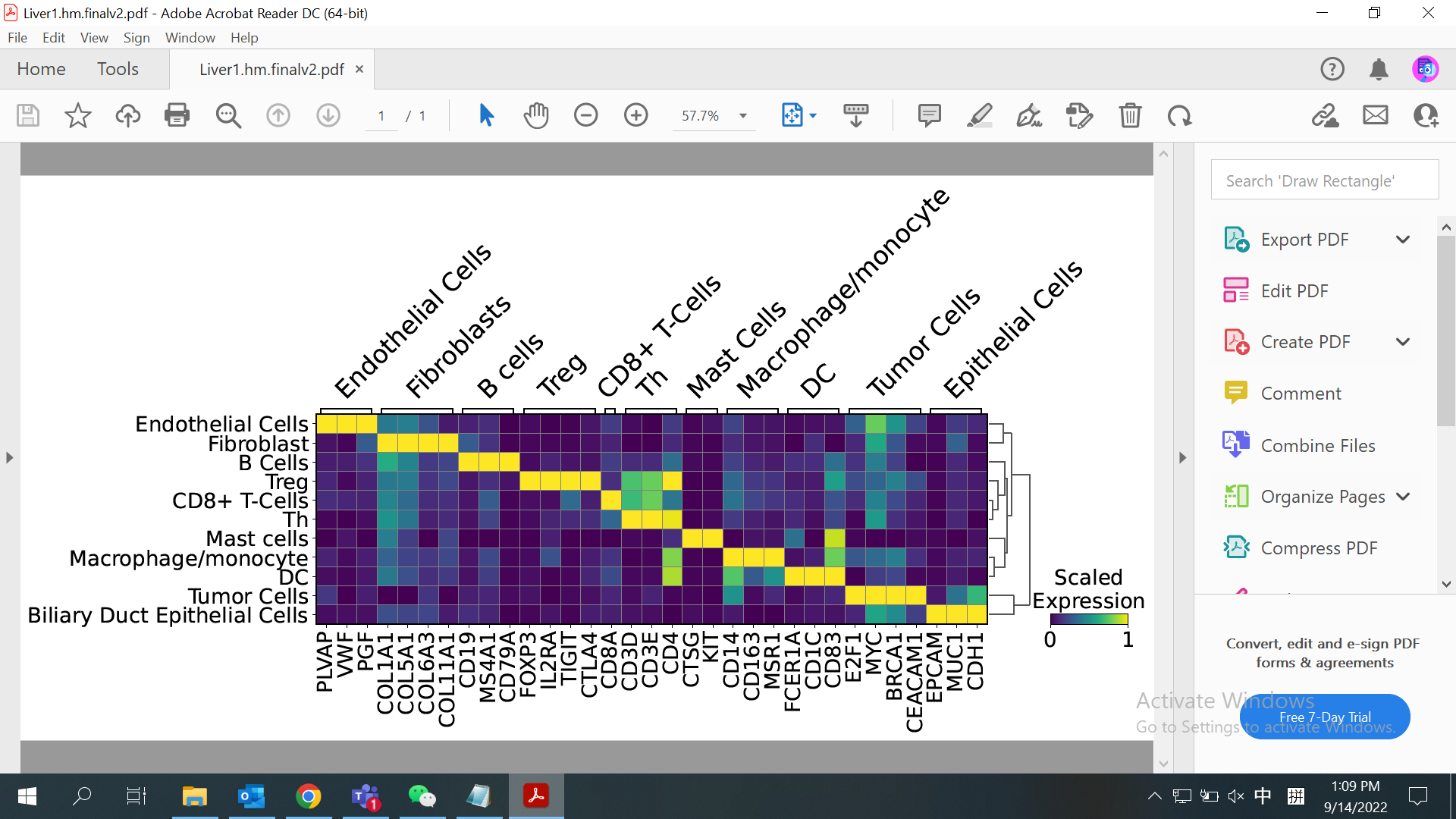

***Sup. File 2 Fig. 11*** *Heatmap showing the expression of the key marker genes for all cell types detected in the Liver Cancer dataset*

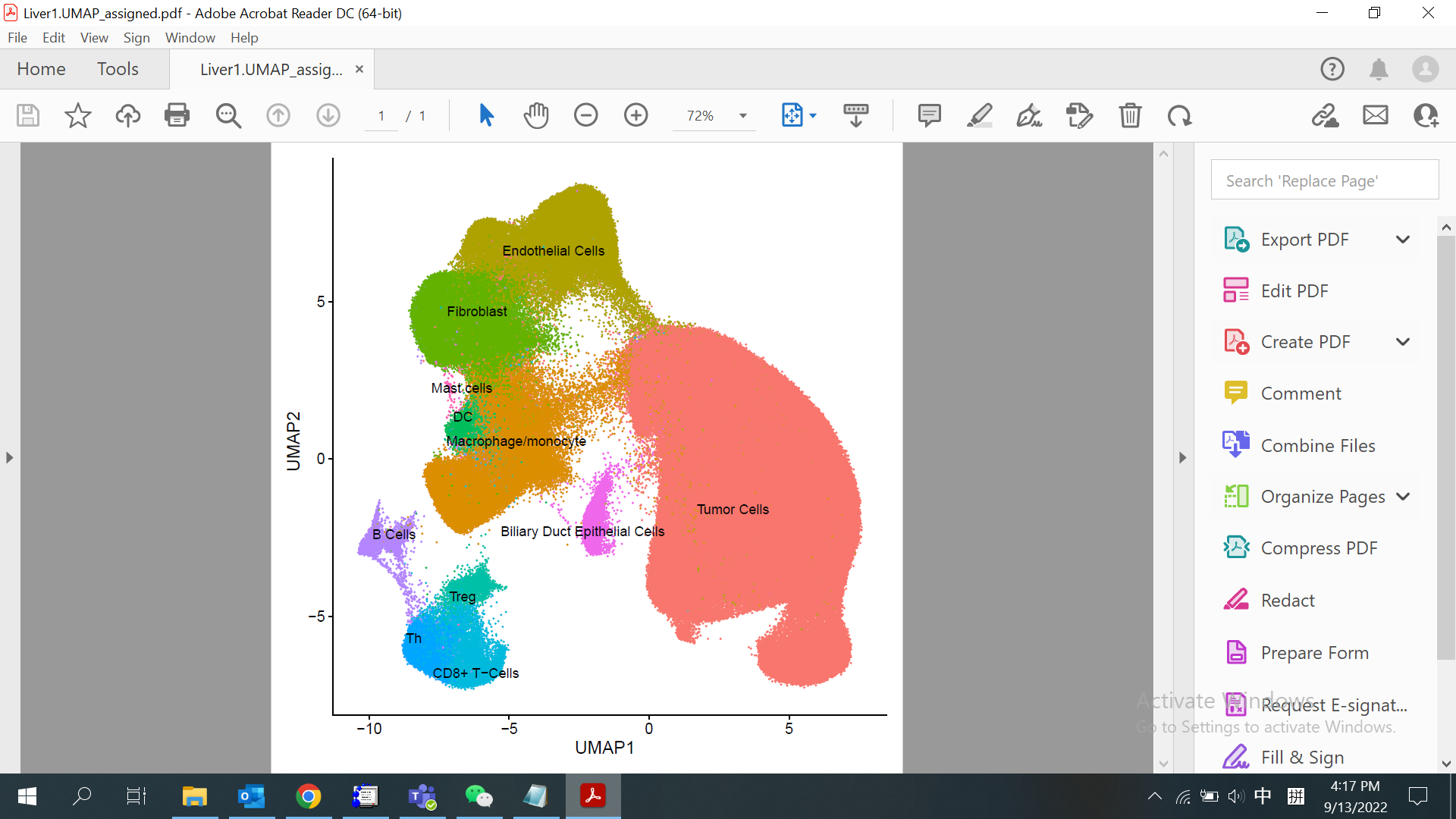

***Sup. File 2 Fig. 12*** *Umap plot showing the cells and their cell types of the Liver Cancer dataset.*

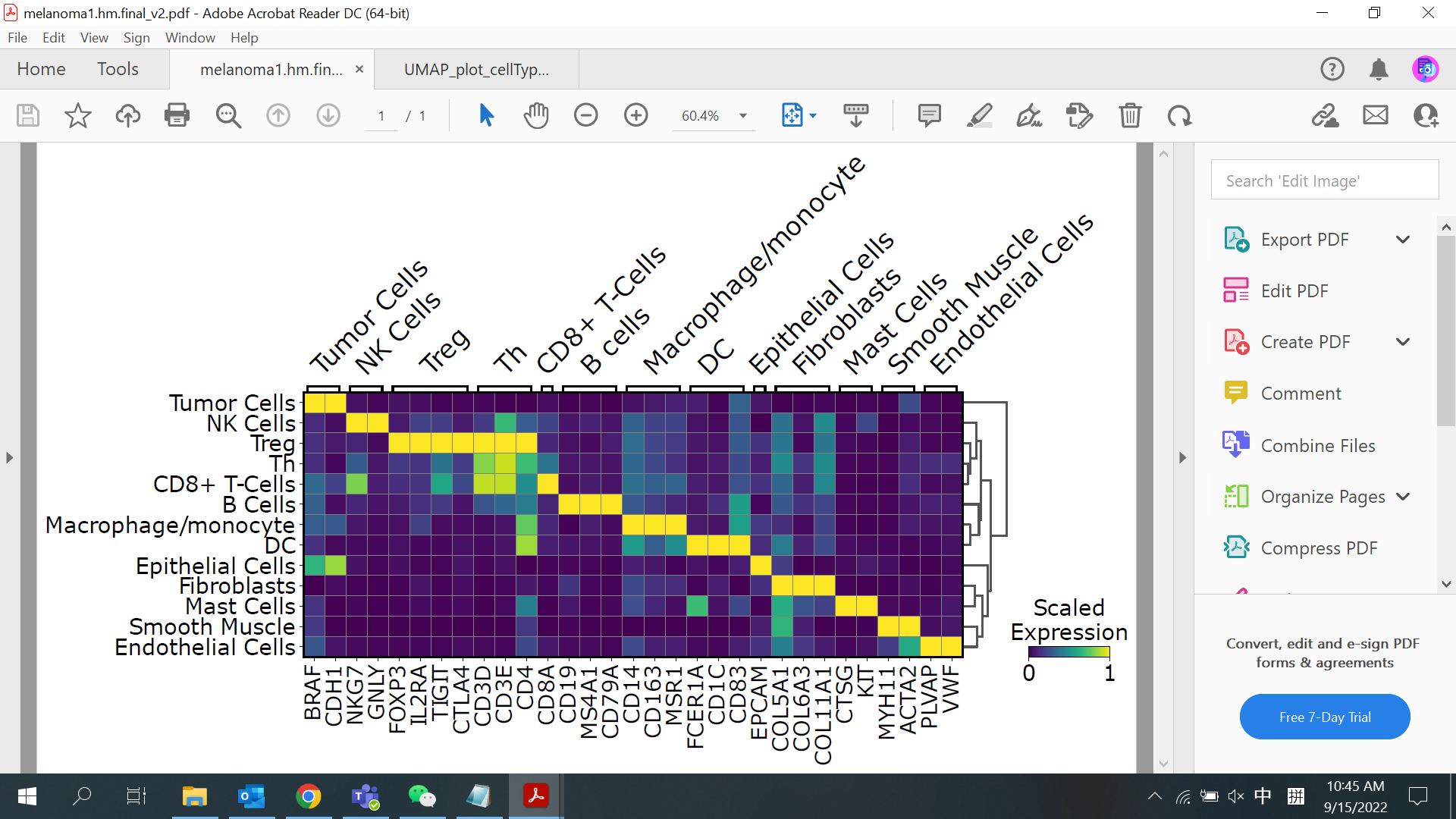

***Sup. File 2 Fig. 13*** *Heatmap showing the expression of the key marker genes for all cell types detected in the Melanoma dataset*

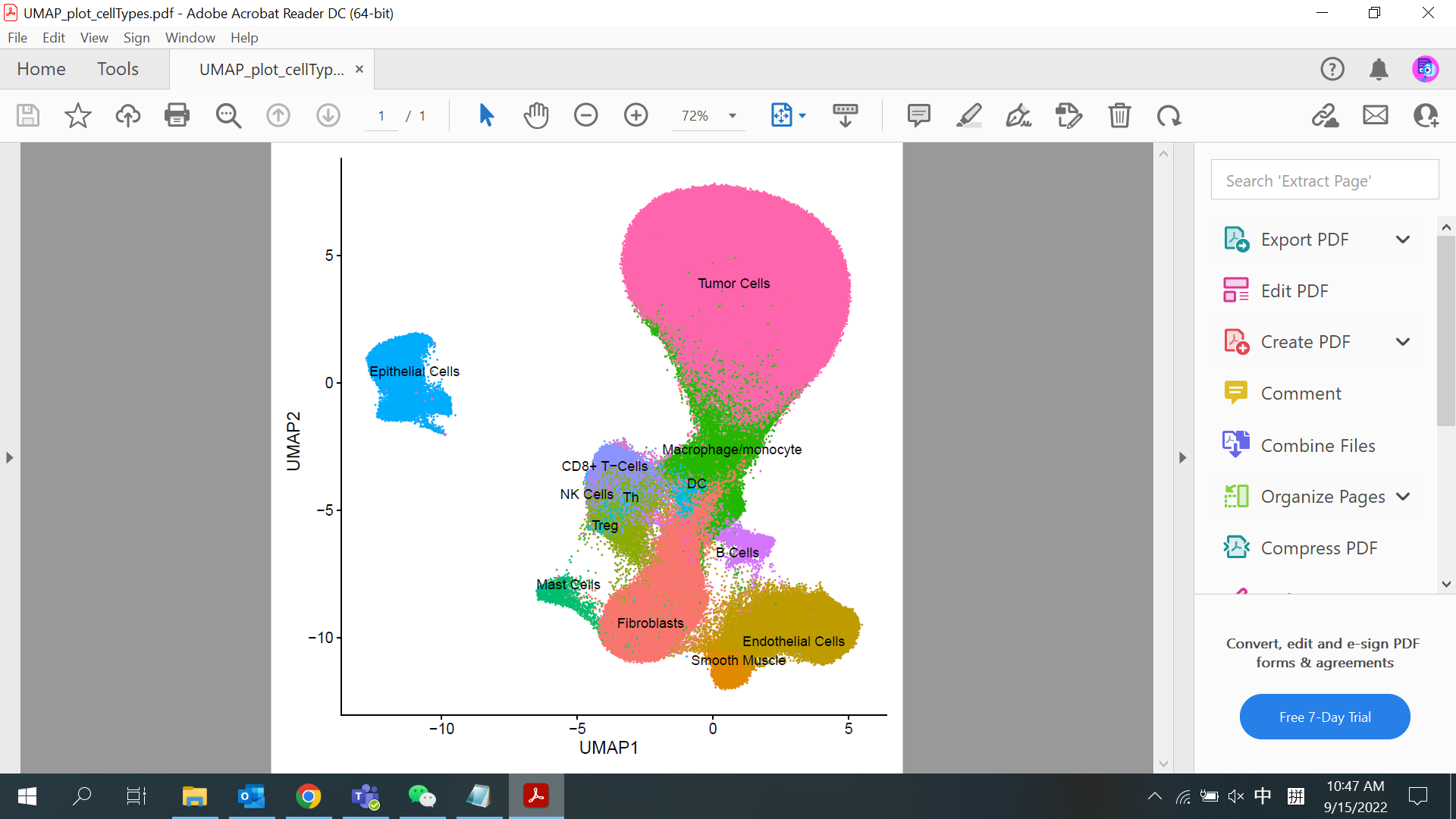

***Sup. File 2 Fig. 14*** *Umap plot showing the cells and their cell types of the Melanoma dataset.*

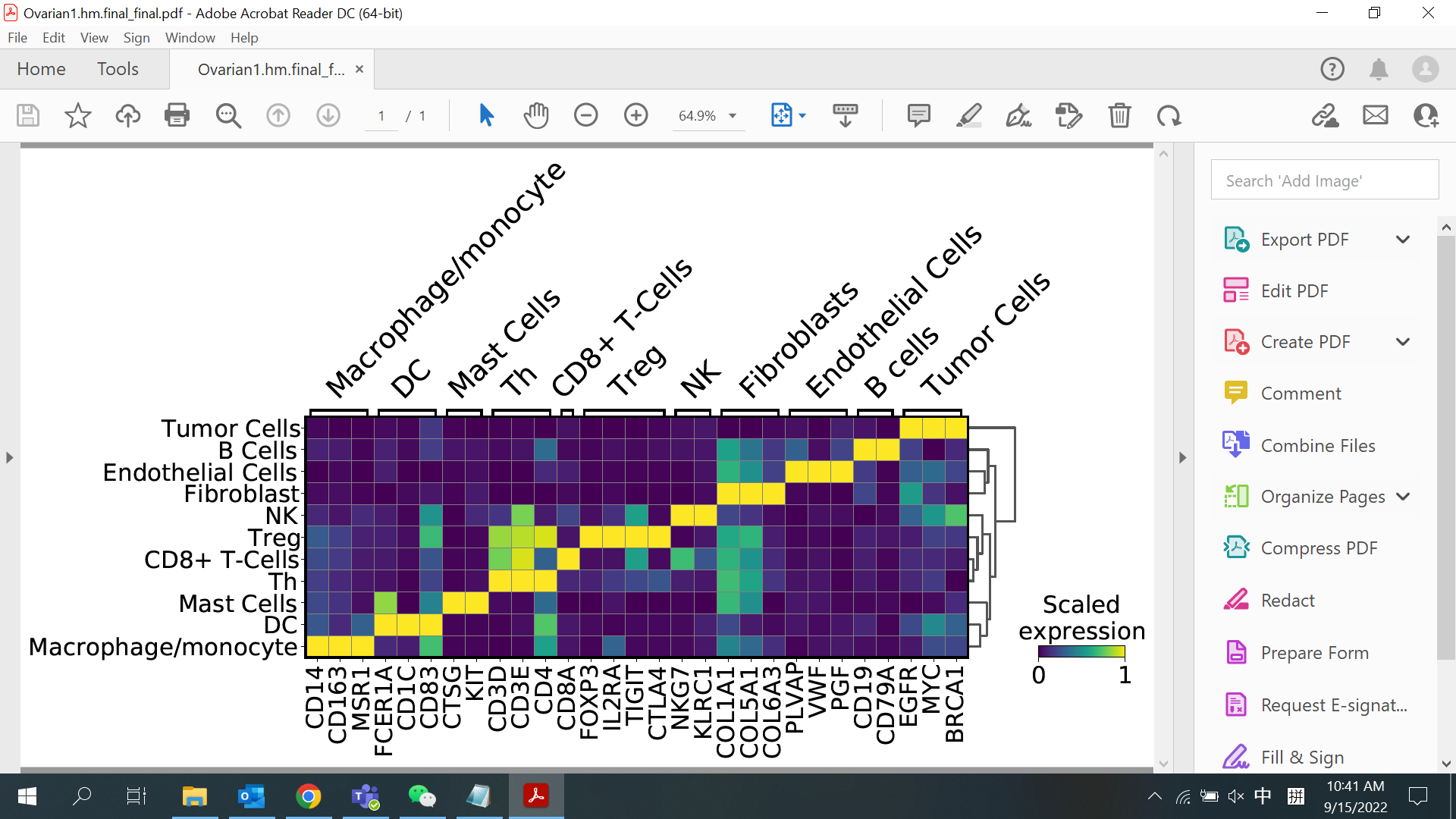

***Sup. File 2 Fig. 15*** *Heatmap showing the expression of the key marker genes for all cell types detected in the Ovarian Cancer dataset*

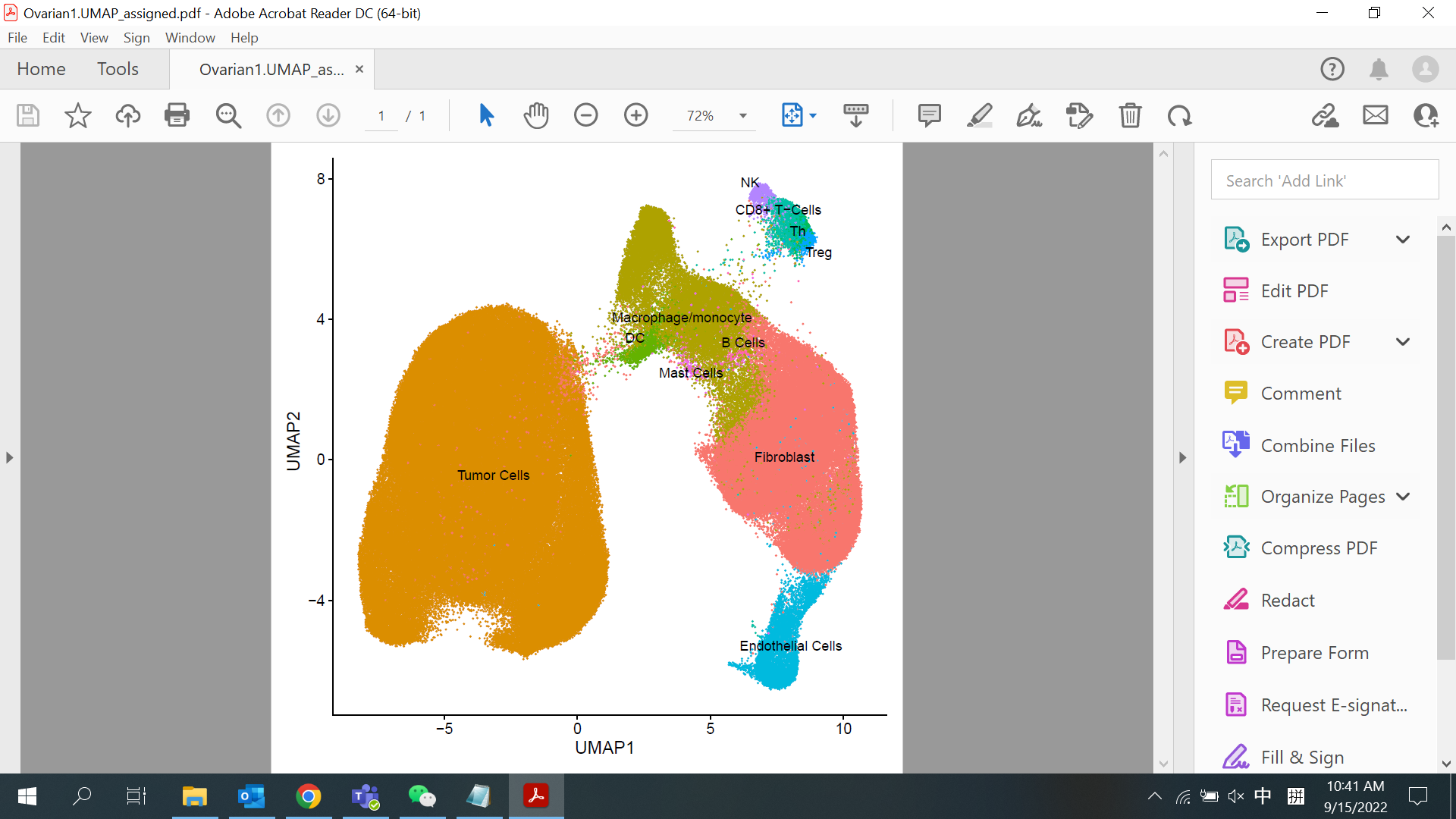

***Sup. File 2 Fig. 16*** *Umap plot showing the cells and their cell types of the Ovarian Cancer dataset.*

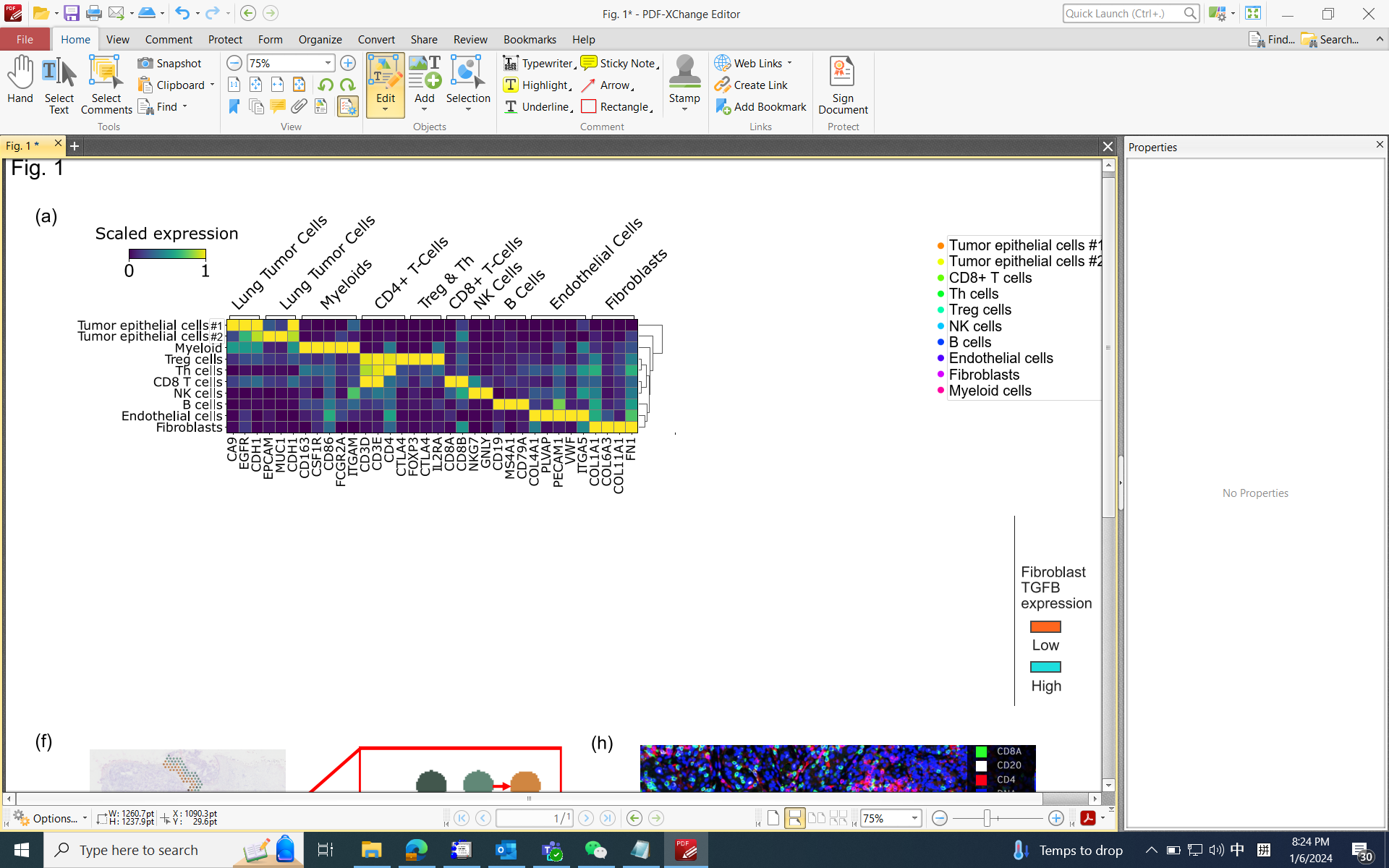

***Sup. File 2 Fig. 17*** *Heatmap showing the expression of the key marker genes for all cell types detected in the Lung Cancer dataset*

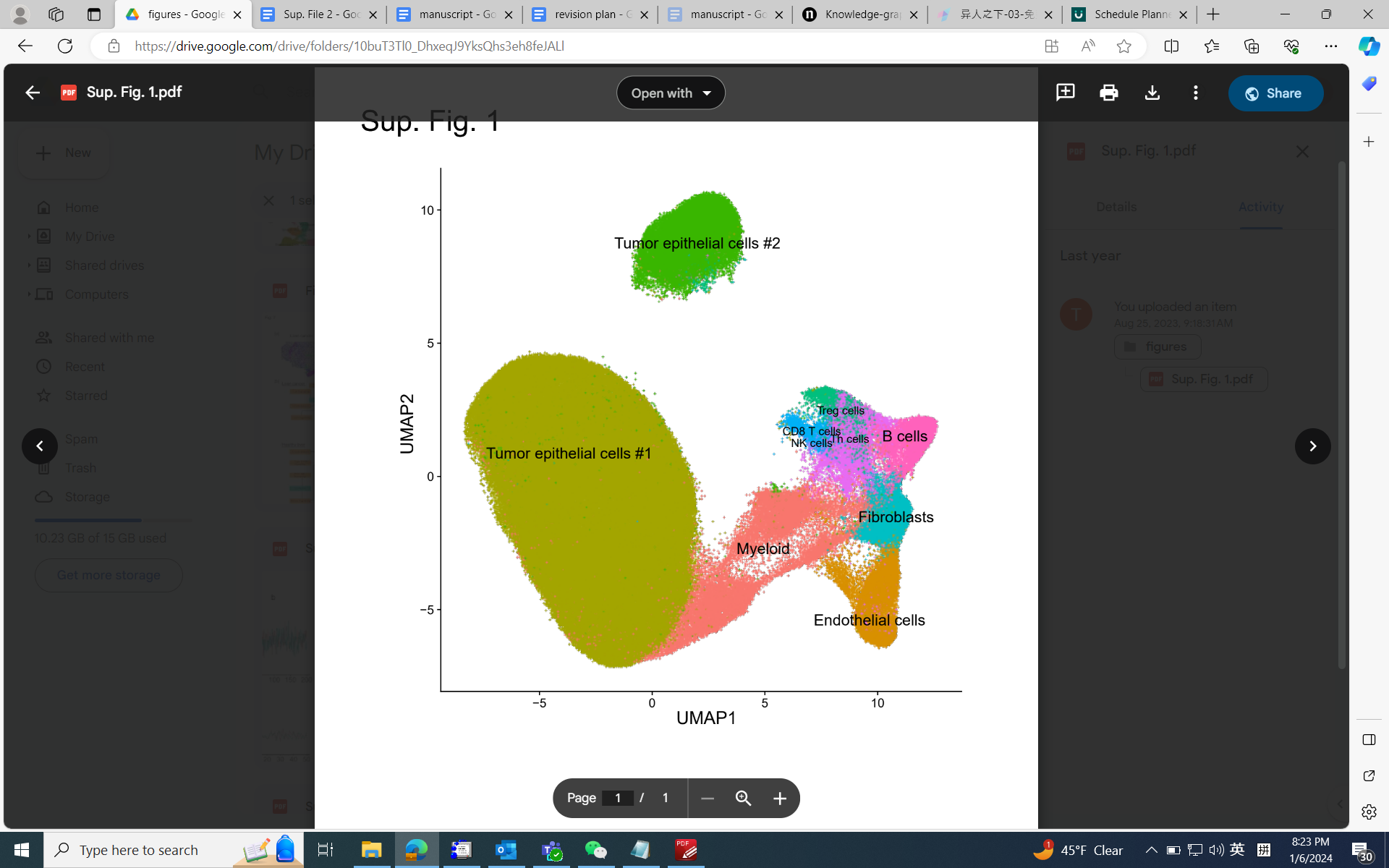

***Sup. File 2 Fig. 18*** *Umap plot showing the cells and their cell types of the Lung Cancer dataset.*

**Validation of the EMT signature in the prostate cancer scRNA-seq data**

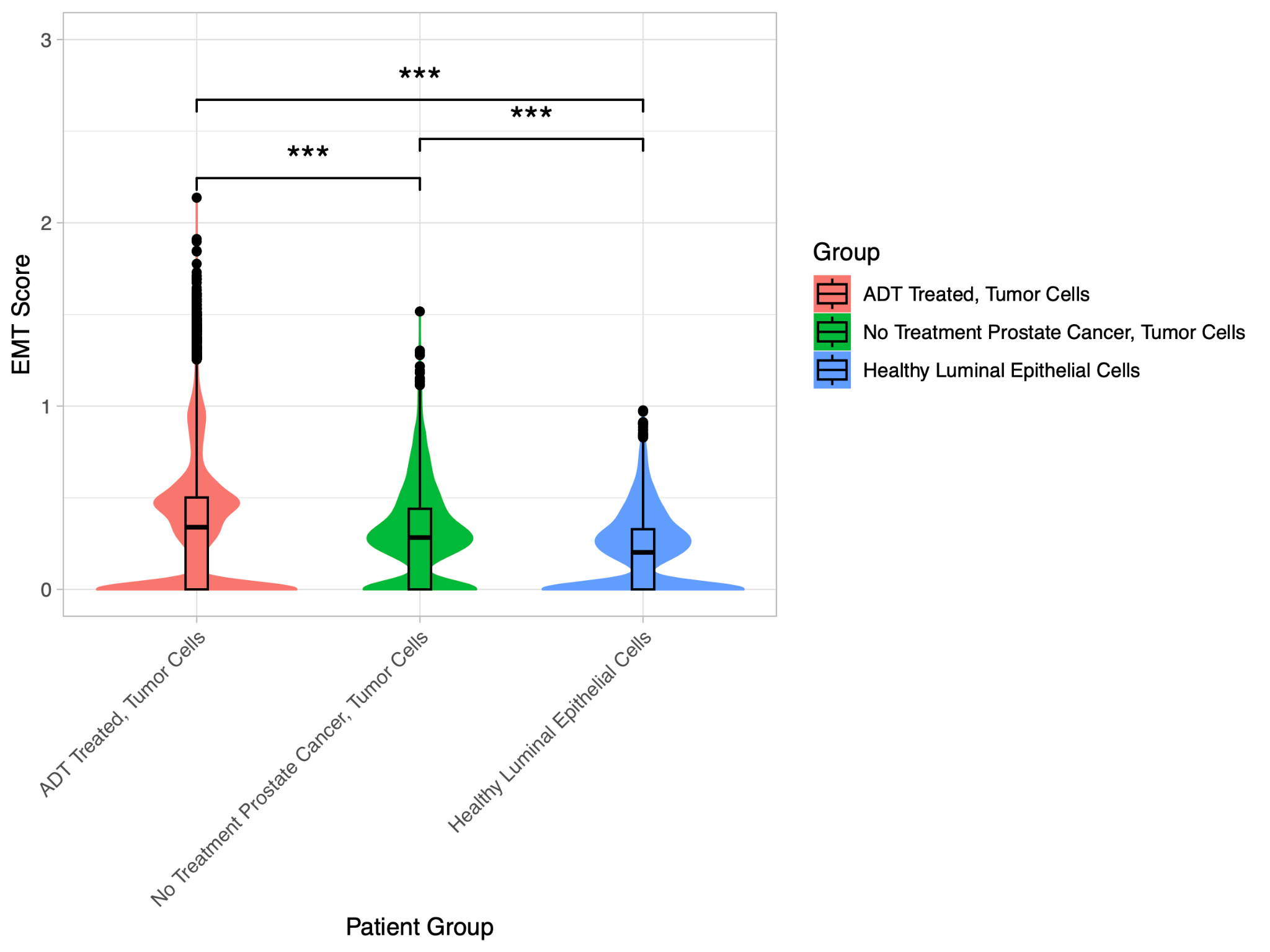

***Sup. File 2 Fig. 19*** *EMT scores in epithelial cells of different patient groups*

To validate that the EMT scores calculated with our gene set are reflective of the true EMT process in the epithelial cells, we examined the expression of this gene signature across three different patient groups in our scRNA-seq data. We compared the EMT scores of tumor cells from androgen deprivation therapy (ADT) treated patients (N=21), tumor cells from untreated prostate cancer patients (N=4), and luminal epithelial cells from healthy donors (N=4). ADT treatment enhances EMT in tumor cells through the Zeb1 transcription factor and the androgen receptor [(*1*)](https://sciwheel.com/work/citation?ids=611891&pre=&suf=&sa=0). Therefore, the EMT scores should be higher in the ADT-treated patients than in the non-ADT-treated patients, which is indeed the case (**Sup. File 2 Fig. 19**). We also expect that the EMT scores of tumor cells from the cancer patients should be higher than the normal epithelial cells from healthy donors, due to a number of factors such as secreted ligands from the tumor microenvironment, which is also validated as true (**Sup. File 2 Fig. 19**). The differences achieved statistical significance in both tests (Pval <0.001, FDR adjusted Wilcox test), which confirms that our EMT scoring metric is valid.

**Performance of benchmark software in identifying EMT-inducing cytokine ligands**

Analogous to our analyses performed in **Fig. 3b** with spacia, we performed the same analyses, but based on the CCC inference output of the five benchmark software applications. Since the benchmark tools rely on the previously curated databases, they do not output all possible pairs of interactions like spacia does. For these data points, we have assigned zero interaction scores as the database determines the pairs as not interacting. Similar to our analyses in **Fig. 3b**, we ranked these interactions and performed a T-test against the mean of 250. The tests for CellChat results yielded no significance for fibroblasts (Pval=0.40), endothelial cells (Pval=0.63), and B cells (Pval=0.65). For CellphoneDB, the results are significant for CD8+ T-cells (Pval=0.04), but not for fibroblasts (Pval=0.30), endothelial cells (Pval=0.56), and B cells (Pval=0.36). COMMOT also resulted in no significance for fibroblasts (Pval=0.20), endothelial cells (Pval=0.85), and B cells (Pval=0.75). Similarly, for both SpaTalk and SpatialDM, the T-test results yield no significance for fibroblasts (Pval=0.90,0.83), endothelial cells (Pval=0.60,0.85), and B cells (Pval=0.20,0.75).

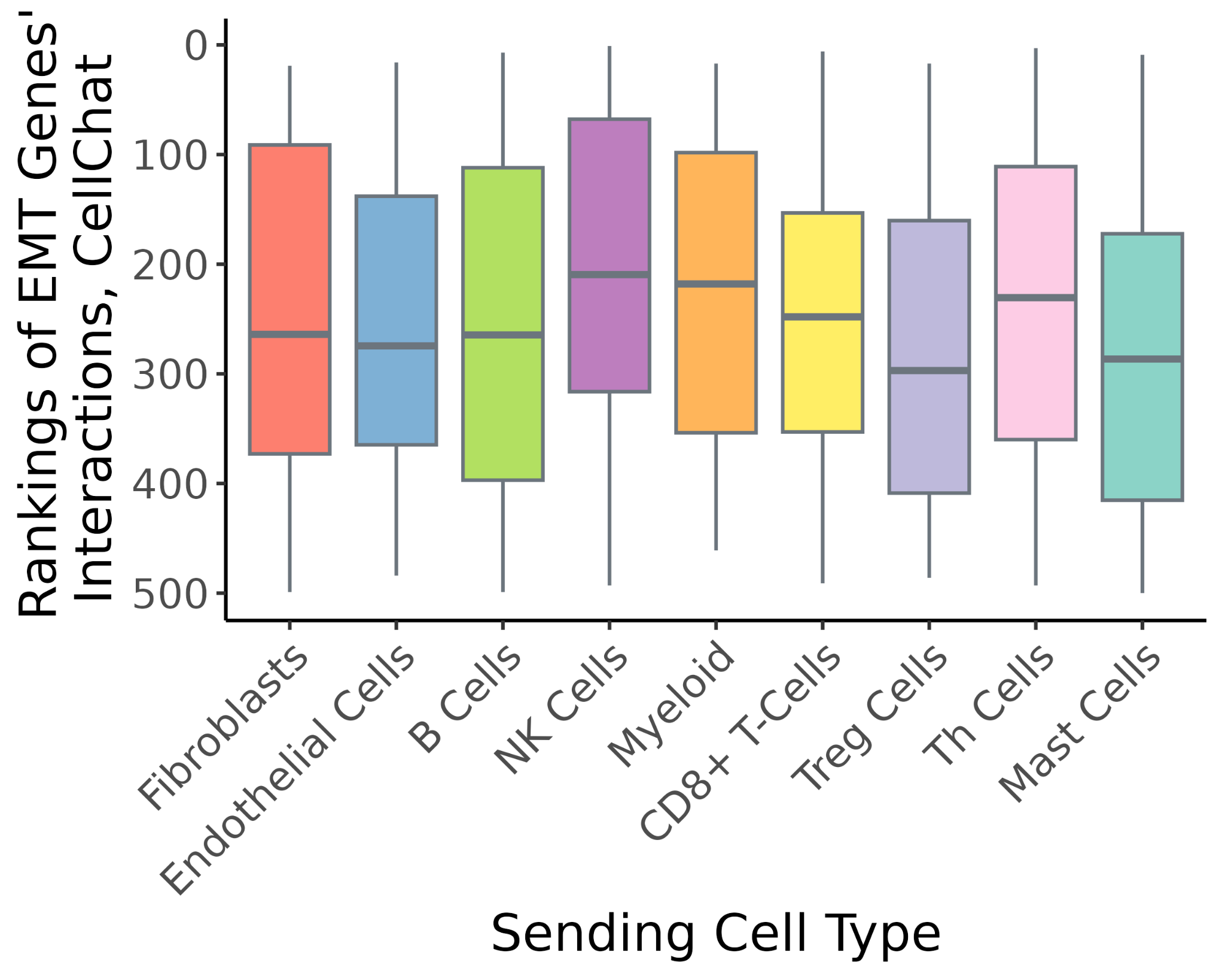

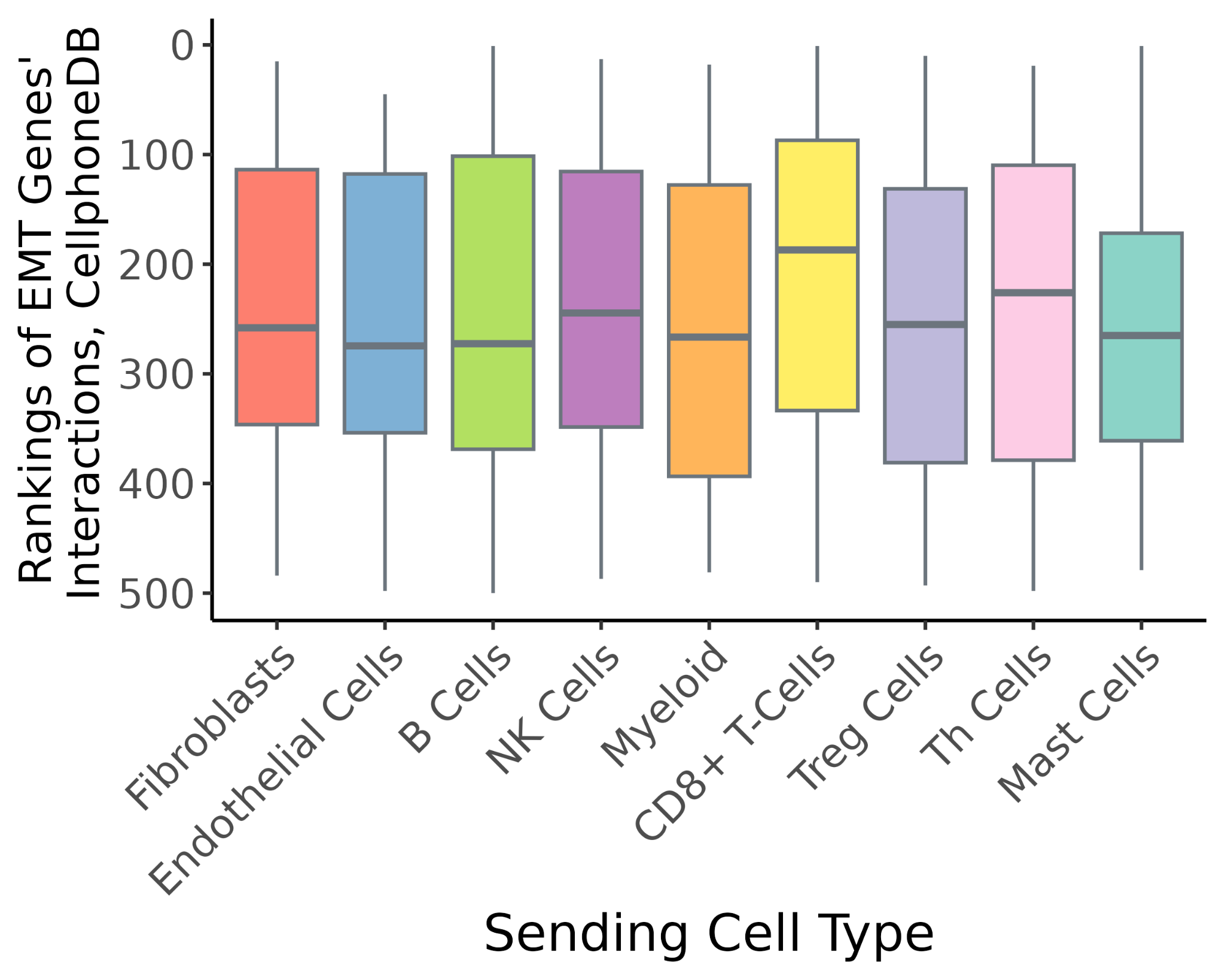

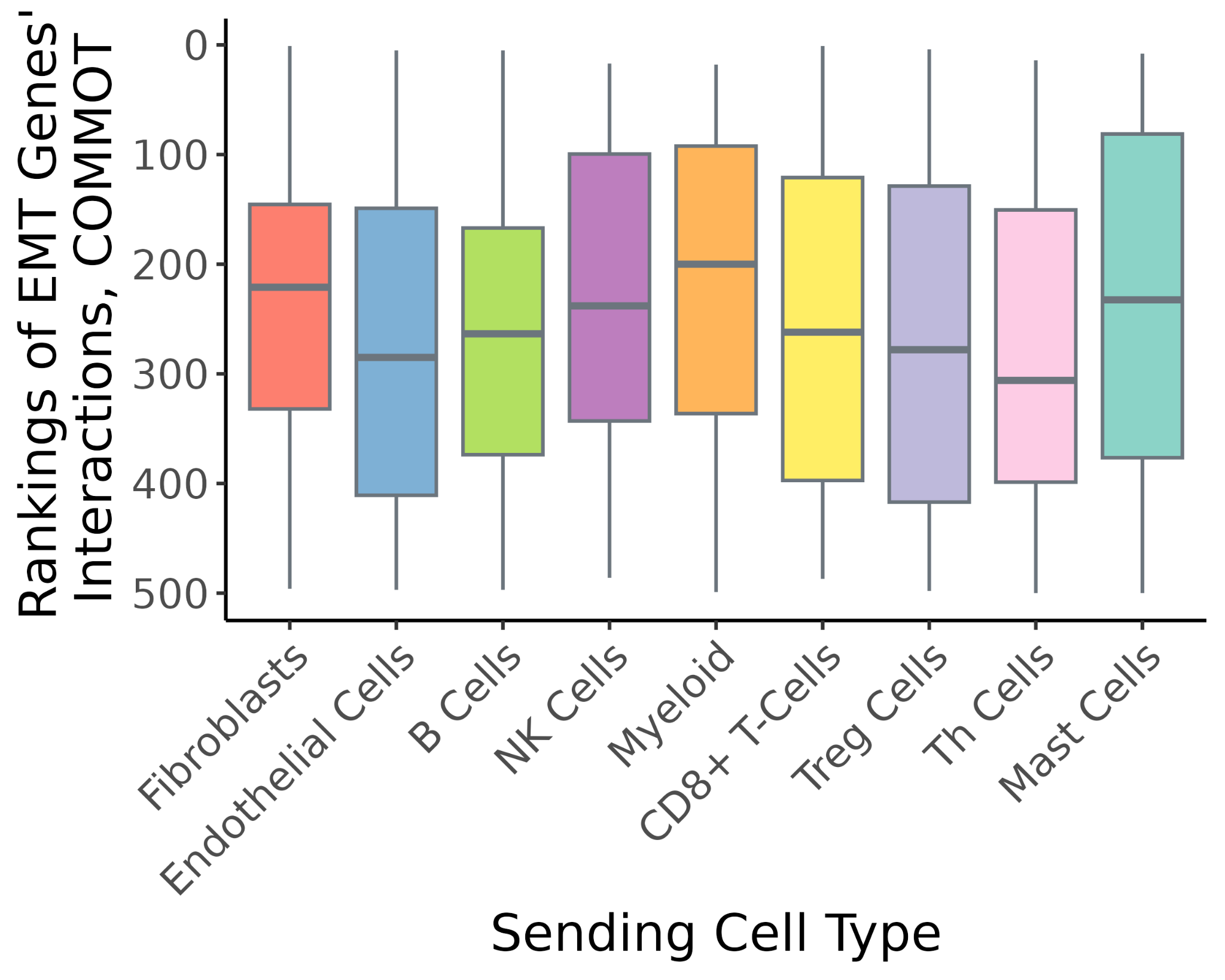

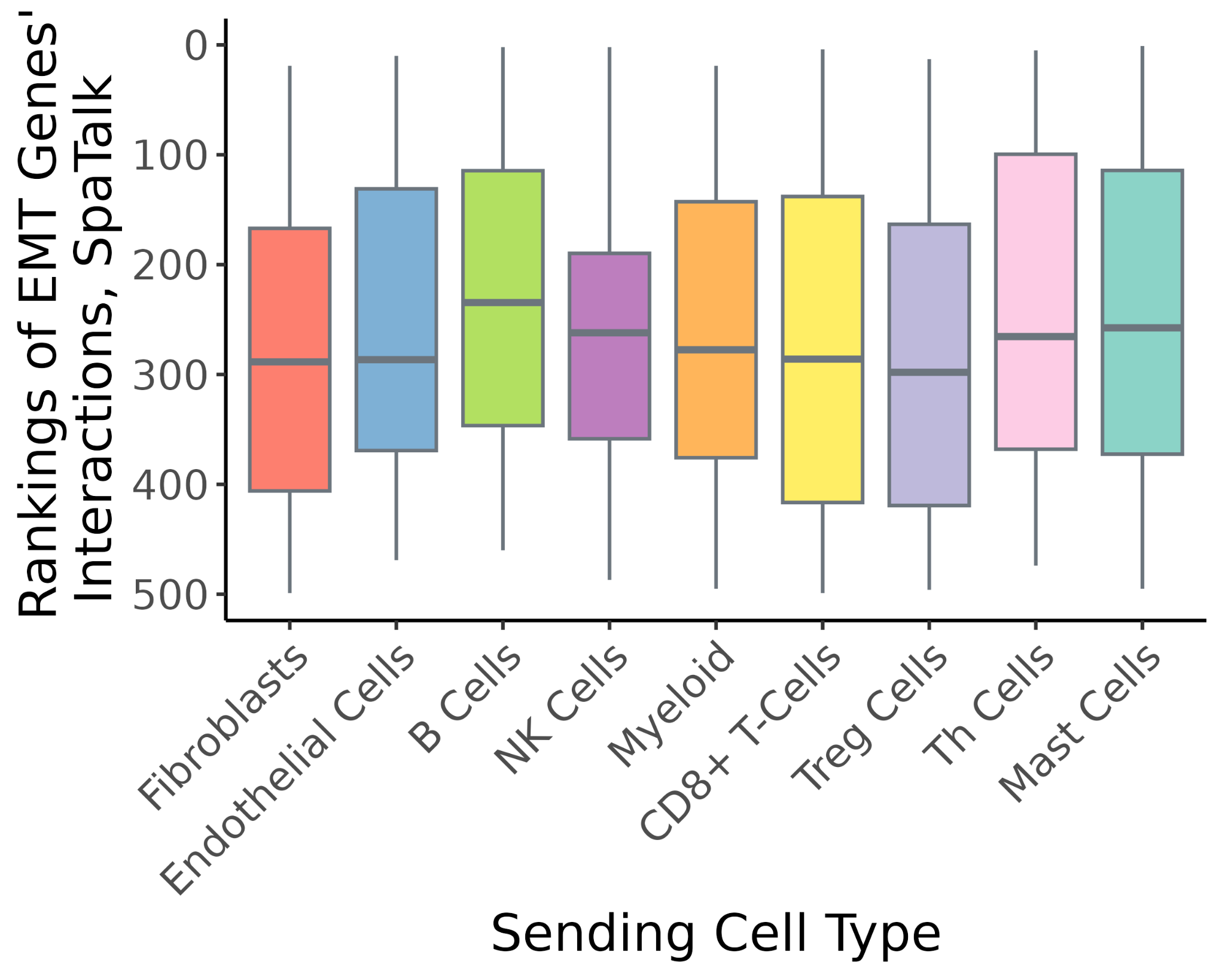

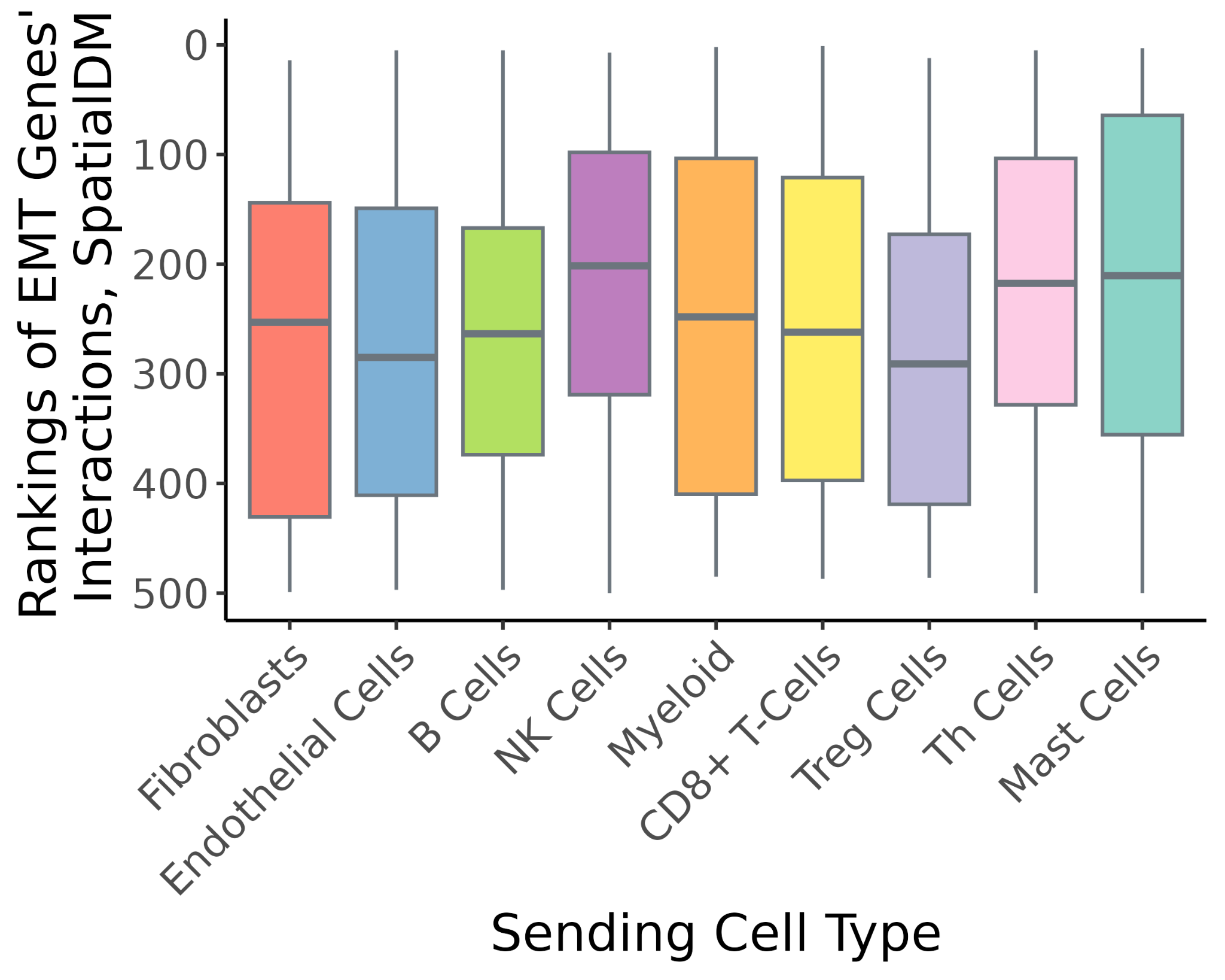

***Sup. File 2 Fig. 20*** *The rankings of the interaction scores for known cytokine ligands that could induce EMT, among all the sending genes input into the five benchmark software applications, for each cell type. A smaller rank refers to a stronger interaction strength (larger absolute interaction score).*

**Full images for one example GeoMX ROI**

To acquire the barcodes separately for each cell type, UV light was directed over a mask pattern defined by the immunofluorescence (IF) markers. In particular, we created a PanCK+ mask for cancer cells (green), a CD45+ mask for immune cells (red) and a PanCK-/CD45- mask for the stromal compartment (light grey). This ROI being shown here only has the tumor mask and the immune mask.

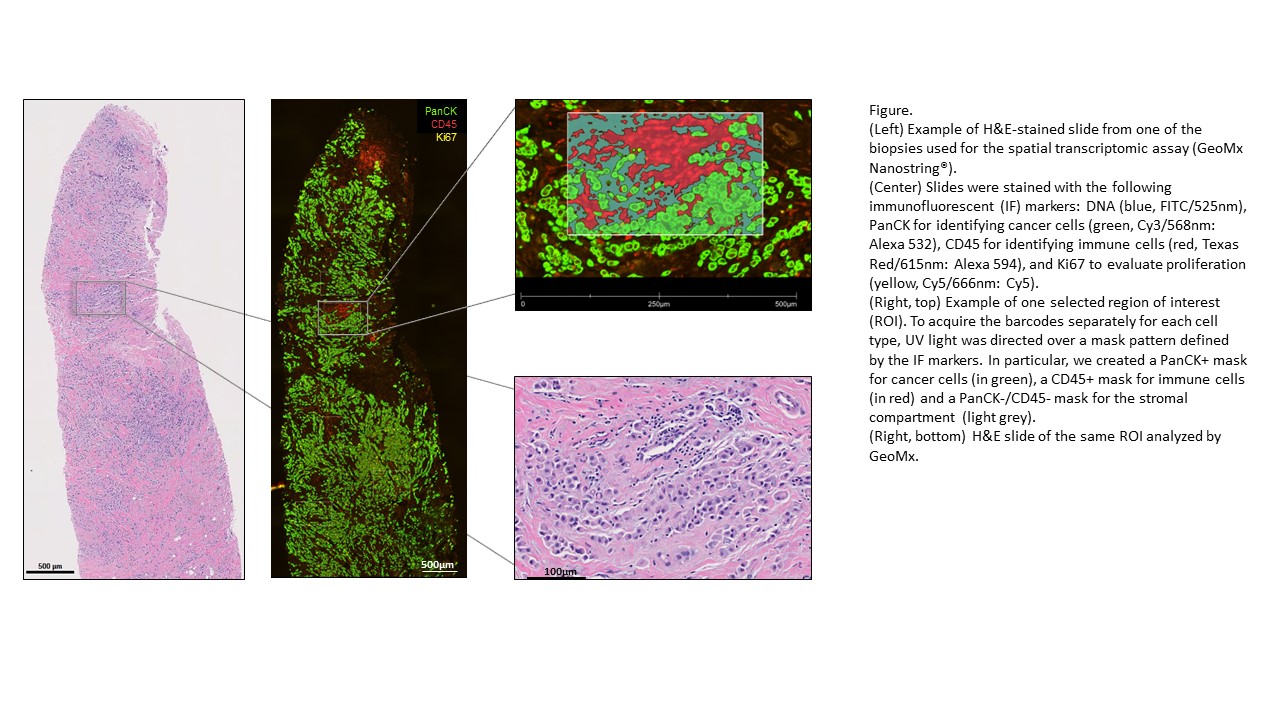

***Sup. File 2 Fig. 21*** *Full images for one example GeoMX ROI. Left: Example H&E-stained slide from one of the biopsies used for the GeoMx assay. Center: Slides were stained with the following IF markers: PanCK for identifying cancer cells (green, Cy3/568nm: Alexa 532), CD45 for identifying immune cells (red, Texas Red/615nm: Alexa 594), and Ki67 to evaluate proliferation (yellow, Cy5/666nm: Cy5). Right, top: One selected region of interest (ROI). Right, bottom: H&E slide of the same ROI analyzed by GeoMx.*

[***Bibliography***](https://sciwheel.com/work/bibliography)

[*1.	Y. Sun, B.-E. Wang, K. G. Leong, P. Yue, L. Li, S. Jhunjhunwala, D. Chen, K. Seo, Z. Modrusan, W.-Q. Gao, J. Settleman, L. Johnson, Androgen deprivation causes epithelial-mesenchymal transition in the prostate: implications for androgen-deprivation therapy. Cancer Res.* ***72****, 527–536 (2012).*](https://sciwheel.com/work/bibliography/611891)
